# Choices of regulatory logic class modulate the dynamical regime in random Boolean networks

**DOI:** 10.1101/2024.12.17.628948

**Authors:** Priyotosh Sil, Suchetana Mitra, Olivier C. Martin, Areejit Samal

## Abstract

Random Boolean networks (RBNs) have been widely explored as models for understanding complex systems, in particular gene regulatory networks (GRNs). Stability (order) and instability (chaos), critical characteristics of any dynamic system, have naturally been a central focus in RBN research. Several measures have previously been introduced to assess stability, e.g., via damage spreading or via statistical properties of attractors such as their number, lengths or basin sizes. Undoubtedly, network topology plays an important role in shaping the dynamics of RBN. Another key factor influencing the dynamics is the Boolean functions (BFs) assigned to each node. In this work, we conduct a systematic investigation to examine the influence of five different classes of BFs on the dynamics of RBNs. By employing various dynamical stability measures, we show that biologically meaningful BFs consistently drive the dynamics toward the stable regime compared to random BFs. This observation holds across networks with varying size, connectivity, and degree distributions. Additionally, we show that the change in the values of our stability measures with increasing connectivity or size of RBNs is different across the BF classes. For most observables, random BFs change the values significantly, pushing the dynamics toward a more chaotic regime with increasing connectivity or network size. In contrast, biologically meaningful BFs show very minimal fluctuations for most stability measures. These findings emphasize the advantage of restricting to biologically meaningful classes in the reconstruction and modeling of biological systems within Boolean framework.

## 1. INTRODUCTION

Gene regulatory networks (GRNs) play a crucial role in orchestrating cellular decision-making processes [1]. Over the past few decades, researchers have made significant strides in unraveling the complexity of these biological networks, shedding light on both their structural organization and dynamical behavior [2–7]. Among the various approaches developed to study GRNs, the *logical modeling* framework has emerged as a particularly powerful tool. Sugita [8, 9], inspired by Jacob and Monod [10], was among the earliest to formalize logical circuits and utilize the Boolean formalism to model GRNs. Soon thereafter, Stuart Kauffman [11, 12] and René Thomas [13, 14] solidified this framework and brought significant attention towards this approach to study the relationships between structure and dynamics of GRNs. In its most common form, logical modeling utilizes Boolean constraints, where genes or their products are considered to be in one of two states: active (*on*) or inactive (*off*).

Kauffman particularly studied the dynamics of Random Boolean Networks (RBNs). He provided some insights on the number and length of attractors as a function of the number (*N*) of nodes in the RBN [11]. Each node in his *N* -node random network is influenced by *k* other random nodes in the network via a randomly chosen logic rule from the set of all Boolean functions (BFs) with *k* inputs. Kauffman suggested that the networks with fixed in-degree (*k*) of 2 were critical networks operating at the *edge of chaos* and hypothesized that the number of attractors in a RBN increases as 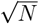. It was also claimed that in RBNs, the typical length of an attractor grows asymptotically as a power of *N* or slower. However, both these conjectures were eventually found to be inaccurate. Specifically, it was shown that both the number of attractors and the mean attractor length grow [15–17] faster than any power law of *N*. Further, Kauffman also found that for *k >* 2, the behavior of the system was chaotic [2].

Subsequently, there has been considerable research on the stability of the dynamics in RBNs when varying network topology as well as the choice of logic rules or BFs [18–21]. For instance, Kauffman *et al*. [18] found that networks with nested canalyzing rules are stable in terms of the total sensitivity of BFs in Boolean networks (BNs) with power law and exponential in-degree distributions. Further, Murrugarra and Laubenbacher [19] analyzed two ensembles of random networks - one employing random rules and the other employing nested canalyzing rules. Their findings revealed that networks using nested canalyzing functions (NCFs) exhibit more robust dynamics, are characterized by fewer attractors whose lengths are also shorter compared to what happens in networks with random rules. Moreover, some of us [22] have shown that NCFs are highly preponderant in the reconstructed biological BNs and NCFs exhibit the minimum average sensitivity for any number of inputs. A meta-analysis of reconstructed BNs for diverse biological systems revealed that those models exhibit more canalyzation and stable dynamics than expected [22–24]. Furthermore, previous studies [25, 26] have also claimed that cells operate in the ordered regime, remaining far from the edge of chaos.

There are several measures that have been utilized to study the stability of dynamics of BNs. In BNs, one can analyze various properties associated with attractors and their basins of attraction, or examine the effect of applying a small initial perturbation and discerning whether the perturbation grows or decreases with time, a process referred to as damage spreading. Specifically, the Derrida coefficient (*δ*) [27] measures the average spreading after one time step and is the same as the so-called *average sensitivity* [26, 28] *of the network (the average of the average sensitivities of the BFs at each node in the BN). δ* is one of the most utilized damage spreading measure [26, 29, 30], and a value greater than 1 of this measure indicates a chaotic dynamics while a value less than 1 indicates stable dynamics. From the point of view of dynamical systems theory, it is appropriate to evaluate the damage spreading at long times, and in that limit, the mean damage is denoted by the measure final Hamming distance or *h*^∞^ [26, 31]. *In the present context, the quantity h*^∞^ can be considered to be the most natural measure of stability when studying BNs.

In studying the impact of BFs on stability of RBNs, significant attention has been given on canalyzing functions (CFs) [18, 20, 21]. Some previous efforts have also touched on the properties when restricting to its NCF subclass [18, 19, 22]. In this work, we perform a detailed and systematic investigation to compare the effect of five different classes of BFs, namely *Effective Functions* (EFs) [32], *Effective Unate Functions* (EUFs) [33], *Read-once Functions* (RoFs) [34], *Nested Canalyzing Functions* (NCFs) [4, 29, 35] and *Link Operator Functions* (LOFs) [36–38] on the stability of RBNs. We also investigate whether the observations change with topological variations, be it through modifications of the average number of inputs, the in-degree and out-degree distributions, or the network size. Specifically, we construct and utilize three different network topological ensembles. In the first ensemble, we fix the in- and out-degrees to be identical for all nodes; in the second, we fix the in-degree for all nodes but allow fluctuations of the out-degree (Poissonian out-degree distribution); in the third, we allow both the in- and out-degree to follow the Poisson distribution. In each ensemble, we consider three different values of network size and four different values of average degree per node (see Methods). Moreover, to probe the stability of BNs, we have considered several measures. One of our sets of measures is associated to the attractors and/or their basins of attraction (see Methods). We also include the recently proposed measure of approximability [39, 40]. Thereafter, we compare the properties of these measures to the established measures of damage spreading *δ* and *h*^∞^, motivated in part by the need to find proxies for *h*^∞^ because it is computationally expensive to determine.

## 2. METHODS

### 2.1. Boolean network modeling framework

A Boolean network (BN) consists of a set of nodes connected by directed edges. Each node in a BN can be in one of two states, “on” (1) or “off” (0). In a BN with *N* nodes, we denote by *x*_*i*_(*t*) the state of the *i*^th^ node at time *t*. Here, *i* ranges from 1 to *N* and *x*_*i*_(*t*) is a binary variable that takes values 0 or 1. At time *t*, the state of the BN can be represented by a vector **X**(*t*) = (*x*_1_(*t*), *x*_2_(*t*), …, *x*_*N*_ (*t*)). The temporal evolution of a BN is determined by its Boolean functions (BFs) (or *logical update rules* or *regulatory logic*), in tandem with an update scheme. BFs may be classified based on several of their properties and we will focus in this work on 5 classes (presented in the next subsection). Update schemes can be either *synchronous* [2] or *asynchronous* [14]. In the synchronous case, all the nodes of the BN are updated concurrently, whereas in the asynchronous case, each node is updated at its particular time. Here we will only consider the synchronous case. More succinctly, 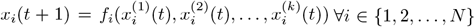, where *f*_*i*_ is the BF at node *i* and 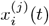 refers to the *j*^th^ input to the BF. Applying the *f*_*i*_ to all nodes takes the network from the state **X**(*t*) to the state **X**(*t* + 1). The dynamics of BNs yield *attractors*. Under synchronous update, attractors can be of two types: *fixed point attractors* or *cyclic attractors*. Fixed point attractors are states that do not change on further update, i.e., **X**(*t*) = **X**(*t* + 1), while cyclic attractors are trajectories whose period is greater than 1.

### 2.2. A hierarchy of classes of Boolean functions

In this work, we particularly employ BFs that are known to have biological relevance [22]. In the ensuing definitions, a BF with *k* inputs is said to be a “*k*-input BF”.

#### 2.2.1. Effective functions

A *k*-input BF is an effective function (EF) [32] if and only if:

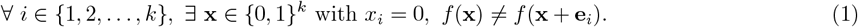

Here **e**_*i*_ ∈ {0, 1} ^*k*^ denotes the unit vector associated to the component of index *i*. Simply put, all the inputs of an EF are effective in the sense that in some context the output changes when that input changes.

#### 2.2.2. Effective and unate functions

Effective and unate functions (EUFs) [33] are effective BFs that are also unate, i.e. the BF’s output is monotonic in each of its inputs. A BF is monotonically increasing in its input *x*_*i*_ if:

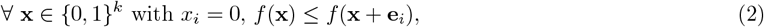

and is monotonically decreasing if:

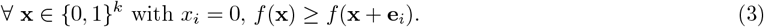

#### 2.2.3. Read-once functions

A *k*-input BF is said to be *read-once* [34] function (RoF) if by using only the operations of disjunction, conjunction and negation, the BF can be written so that each of the *k* variables appears exactly once in that expression. Mathematically, this may be succinctly stated as:

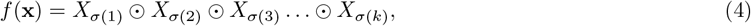

where 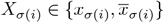 represents the AND (∧) or OR (∨) operator, *σ* represents the permutations on the set 1, 2, …, *k* and *σ*(*i*) is the *i*^th^ element of the set. Note that, for legibility of Eq. 4, we have omitted all parentheses necessary to completely specify the BF. But given that writing, clearly RoFs are a subset of EUFs.

#### 2.2.4. Nested canalyzing functions

A *k*-input BF is a nested canalyzing function (NCF) [4] if:

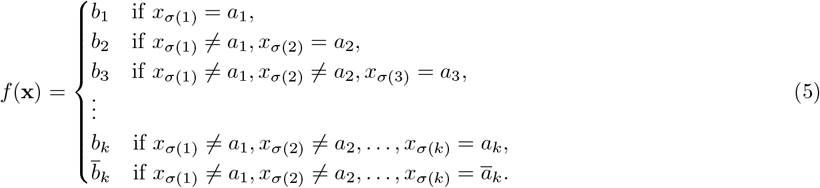

In the above equation, *a*_1_, *a*_2_, …, *a*_*k*_ are the canalyzing input values and *b*_1_, *b*_2_, …, *b*_*k*_ are the canalyzed output values for inputs *σ*(1), *σ*(2), …, *σ*(*k*) in the permutation *σ* of the *k* inputs. Clearly, NCFs are a subset of RoFs.

#### 2.2.5. Link operator functions

A *k*-input BF is a link operator function (LOF) [36–38] if it can be represented in one of the following 4 forms:

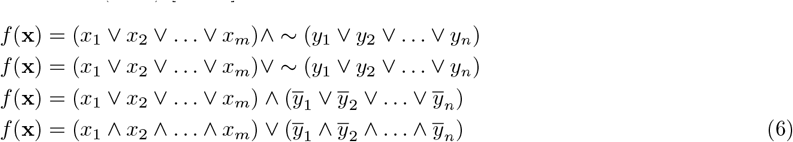

Note that LOFs are also a subset of RoFs but are neither a subset nor a superset of NCFs.

### 2.3. Stability of the Boolean network dynamics: Derrida’s damage spreading and generalizations

A common way to test for stability of dynamics in BNs is to determine how two trajectories, initially separated by a single bit change, diverge with time. This is formalized via the so-called Derrida map [41] connecting the magnitude of the distance between the two trajectories (sometimes called *damage*) at time *t* + 1 to that at time *t*. If the initial damage tends not to spread, the system is considered to be *stable*, whereas the opposite case is taken as a signature of *chaotic* behavior. For this work, we consider three measures of stability [26] based on damage spreading: (a) the Derrida coefficient (*δ*), (b) the final Hamming distance (*h*^∞^), and (c) the fragility (*ϕ*).

The Derrida coefficient, *δ*, is just the average damage size after a single step of the dynamics. If *δ >* 1, the damage is likely to spread to a finite fraction of the network, while if *δ <* 1, one expects the damage to die out, in which case the system may be considered to be in the ordered regime. Note that *δ* is equal to the average network sensitivity of the system [26, 42] and hence depends only on the BFs and not on the network architecture.

The measure “final Hamming distance”, *h*^∞^, quantifies the long-term average separation between trajectories. It is mathematically defined as

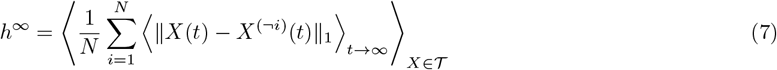

Here, *X*(*t*) is the network state at time *t* without the initial perturbation while *X*^(*¬i*)^(*t*) is the state that is obtained by evolving the state that initially had the (single bit) perturbation. 𝒯 corresponds to the set of all trajectories. The symbol ⟨·⟩_*X*∈*T*_ denotes the average taken over all possible initial network states and ‖*X*(*t*_*f*_) − *X*^(*¬i*)^(*t*_*f*_)‖_1_ represents the L1-norm, i.e., the Hamming distance, between the states *X*(*t*_*f*_) and *X*^(*¬i*)^(*t*_*f*_) at time *t*_*f*_. Finally, ⟨‖*X*(*t*) − *X*^(*¬i*)^(*t*)‖_1 *t*→∞_⟩ denotes a time average taken in the limit of long times.

Given that *h* _∞_ is sensitive to phase shifts arising in cyclic attractors, Park *et al*. [26] proposed a modification to Eq. 7, leading to what they call “fragility”, defined by:

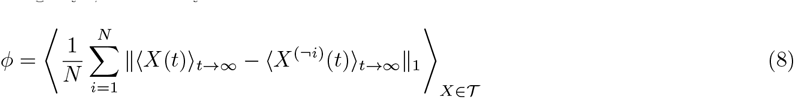

The difference between *ϕ* and *h*^∞^ is that in the case of *ϕ* the time average is taken inside the norm, a choice that removes the sensitivity to the phase shift in cycles.

### 2.4. Other proxies for stability based on properties of the attractors

In this work we analyze 7 different observables related to the attractors and their basins of attraction which are as follows.

i. The total number of attractors: This is obtained by summing the number of fixed points and the number of cyclic attractors.
ii. Fraction of fixed point attractors: This is the ratio of the number of fixed point attractors to the total number of attractors. If there are a total of *n* attractors, out of which *m* are fixed point attractors, then this quantity is given by *m/n*.
iii. Fraction of states in the basins of fixed points: This is the ratio of the number of states converging to fixed point attractors to the total number of states, 2^*N*^. If for an *N* -node BN, there are *m* fixed-point attractors with the basin sizes (number of states in the basin of attraction) *W*_1_, *W*_2_, …, *W*_*m*_ respectively, then the quantity we consider is given by 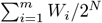.
iv. Average attractor length: The length of an attractor refers to the number of states that constitute the attractor. Fixed points are attractors of length 1, whereas cyclic attractors can have lengths ranging from 2 to 2^*N*^ in an *N* -node BN.
v. Maximum attractor length: This quantity refers to the length of the largest attractor.
vi. Weighted average attractor length: If for an *N* -node BN, there are *n* attractors, respectively with attractor lengths *L*_1_, *L*_2_, … *L*_*n*_ and basin sizes *W*_1_, *W*_2_, … *W*_*n*_ (summing up to 2^*N*^), then the weighted average attractor length is given by

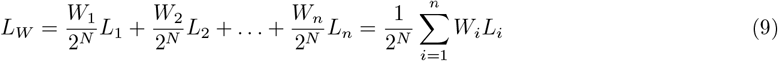
vii. Fraction of states on attractors: For an *N* -node BN, this quantity refers to the fraction of the network states belonging to attractors. If there are *n* attractors with attractor lengths *L*_1_, *L*_2_, … *L*_*n*_, then the quantity is given by 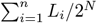

### 2.5. From polynomial approximations of regulatory rules to the “approximability” of network dynamics

In continuous dynamical systems, major insights follow from linearizing the equations of motion. In the case of BF, the only way to have a linear update rule is for it to depend on a single variable; BFs of multiple variables are therefore intrinsically nonlinear. Nevertheless, some BFs may be more nonlinear than others. A question thus immediately comes to mind: do the different classes of BFs (e.g., EFs, EUFs, …) have significantly different levels of nonlinearities? To address this question, we follow the framework of Manicka *et al*. [39] who in effect use the Glauber formula [43] to write a BF, *B* as a sum of polynomial terms:

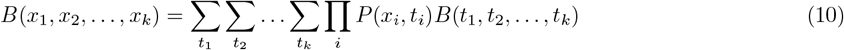

Where

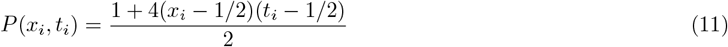

is a Kronecker delta function on Boolean variables, but written so it can also be applied to real variables (Note that the product of these Kronecker functions is analogous to the use of Lagrange interpolation polynomials). Eq. 10 then directly provides a polynomial representation of the BF to which one can apply the standard tools of continuous dynamical systems. To proceed, the polynomial of Eq. 10 is Taylor expanded about the mid-point of the hypercube (all *x*_*i*_ = 1*/*2) and then truncated at the desired order. The order 0 case leads to the constant approximation, the order 1 case to the linear approximation, etc. As a result, the degree of nonlinearity of a BF can be quantified by the error introduced when using the degree 1 approximation. More generally, for any degree used in the truncation, one can define an error via the square of the difference between the value of the BF and that of the truncated polynomial when averaging over all 2^*k*^ possible input values. We call this the mean error of truncation and it depends only on the BF and the truncation degree. The mean error of truncation of a BF when the truncation degree is 1 will be used as a measure of nonlinearity of that BF, thereby allowing us to compare the nonlinearities across the different classes of BFs.

Manicka *et al*. [39] were interested in how well a truncated dynamics could approximate the true Boolean dynamics at the level of the whole network. Rather than consider the approximation for a single time step, they focused on the long-term behavior, imposing that the error be evaluated after 500 time steps. They call this the MAE (for mean approximation error) where the mean is obtained by averaging over as many random initial conditions as possible and over all the variables of the network. For the MAE results provided in the present work, we have followed exactly that prescription from which we compare the behavior of the different classes of BFs.

### 2.6. Ensembles of RBNs: constraining network topologies and restricting Boolean functions to a class

Past studies on random Boolean networks (RBNs) used random BFs at each node but made more specific choices for the network architecture, i.e., who drives whom, generically referred to as the topology of the network. The two most common choices are as follows. In the first, one assigns (oriented) interactions between randomly selected nodes, subject to a given average number *λ* of edges per node. This procedure corresponds to the Erdös-Reyni [44] ensemble (but with oriented edges), hereafter denoted P-P as when the number of nodes becomes large, the distribution of in-degrees (*k*_*in*_) and of out-degrees (*k*_*out*_) becomes Poissonian of parameter *λ*. In the second, one imposes that both the in- and the out-degrees be fixed, corresponding to what is called the *k*-regular graph ensemble, hereafter denoted R-R. That choice was motivated [45] by the desire to ensure that the nodes had comparable update rules since two rules with different numbers of inputs are difficult to compare. In addition to those two ensembles of network architectures, in this work we consider a mixed case where one fixes the in-degree to one particular value [12], leaving free the out-degree so it stays Poissonian. We will call it the R-P ensemble and it will allow us to test the relevance of fluctuations in the out-degree on the network dynamics.

In addition to controlling network architecture, for our study we also control the type of BFs used at the nodes of the network. Specifically, within each of the 3 topological ensembles introduced (P-P, R-R and R-P), we will not consider random logical update rules but instead rules that have particular properties such as unateness or are canalyzing. Thus, we will impose that each node in the network is assigned a rule at random within the considered class (respectively EF, EUF, RoF, NCF, and LoF). Note that for all practical purposes, the EF class essentially corresponds to random BFs, the only constraint being that those BFs be effective; when the number of inputs is 3 or more, this constraint is nearly always satisfied for a random BF [22].

As a technical point, the assignment of BFs to nodes requires that we be able to randomly draw BFs with a uniform measure within each of the 5 classes considered. Unfortunately, this becomes computationally infeasible for certain classes such as EUF when the in-degree becomes larger than 6. Thus, in the case of the P-P ensemble, we can be confronted with situations where the in-degree is too large to guarantee a uniform drawing. If so, we resort to a sampling that only approximates the uniform case but that has nevertheless been shown to deviate little from uniformity [42].

#### Implementation

We generate the ensembles of RBNs as follows: (i) We use 1000 networks in each case. (ii) We consider networks with number of nodes *N* = 12, 16 and 20 and analyse different properties associated with the attractors and MAE of the network. (iii) *δ, h*^∞^ and *ϕ* are computed for networks with *N* = 12. (iv) The in-degrees (or mean in-degrees in case of Poisson in-degree distribution) considered are 2, 3, 4 and 5. Nevertheless, in the case of Poisson degree distribution, the in-degree is kept within the range 1 and 7. The value 1 is to ensure that the node has a BF attached to it (this corresponds to the so called zero-truncated Poisson distribution) while 7 is to ensure that we can uniformly (or almost uniformly) sample BFs in each class as explained above. (v) The classes of functions considered are EF, EUF, RoF, NCF and LOF. (vi) We generate 1000 models each for every class per network to study different network attractor properties. (vii) We generate 10 models each for every class per network topology for the computation of *δ, h*^∞^, *ϕ* and MAE.

### 2.7. The Jensen–Shannon distance and statistical tests for comparing distributions

Since we use an ensemble approach, each of the observables can be thought of as a random variable. We seek to characterize the associated distributions and to compare them as there are usually visible differences when examining their violin plots. A standard way to compare two distributions is to implement a binning and measure the Kullback-Leibler divergence of the resulting histograms:

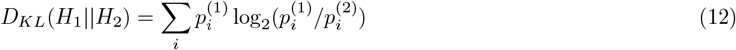

where 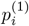 and 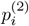 are the respective probabilities of falling in bin *i* according to the histograms *H*_1_ and *H*_2_. *D*_*KL*_|| (*H*_1_ *H*_2_) vanishes when the two histograms are identical and it rises as the difference between the two histograms increases. Note that in general *D*_*KL*_(*H*_1_ || *H*_2_) ≠ *D*_*KL*_(*H*_2_ || *H*_1_) = 0, *i*.*e*., this divergence is not a distance and not even symmetric. Thus, to follow common practice, we use in all our comparisons a symmetric quantity, the so-called Jensen–Shannon distance:

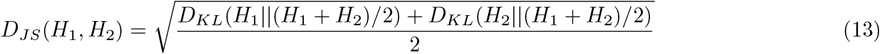

This has the remarkable property of really providing a distance, that is, the order of the arguments does not matter and this measure satisfies the triangle inequality. Eq. 13 allows us to quantify how different two distributions are via a *distance* that conveniently lies in the range [0, 1].

Given any two empirical distributions, i.e., finite samples each drawn from their own distribution, one can test whether they might have been drawn from the same distribution. The Python function scipy.stats.ks 2samp provides a p-value for that hypothesis using the standard Kolmogorov-Smirnov test. What is more interesting is the prospect of testing the possibility that the first distribution dominates the second one, i.e., its cumulative (or *integrated*) distribution is everywhere higher than that of the other. A p-value for that hypothesis is provided by the scipy.stats.ks 2samp function by requiring the test to be one-sided.

## 3. RESULTS AND DISCUSSION

### 3.1. Stability is enhanced along our hierarchy of Boolean function classes

The measure *h*^∞^ [26] quantifies the long-term average separation between trajectories with an initial Hamming distance of 1. In Fig. 1, each point is associated with a graph chosen at random within the R-R topology ensemble (*k*_*in*_ = *k*_*out*_ = 4 and *N* = 12). The value on each axis is the average of *h*^∞^ over 10 different realizations of the BF rules when taken at random from the corresponding labeled class. Broadly, we observe 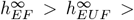 (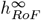 or 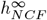 or 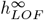), with the last three classes RoF, NCF and LOF not following any particular ordering. This hierarchy of *h*^∞^ indicates that progressing from EF to EUF to any of the three remaining classes (RoF, NCF or LOF) steers the dynamics towards the ordered regime (or away from the chaotic regime).

**FIG. 1.**
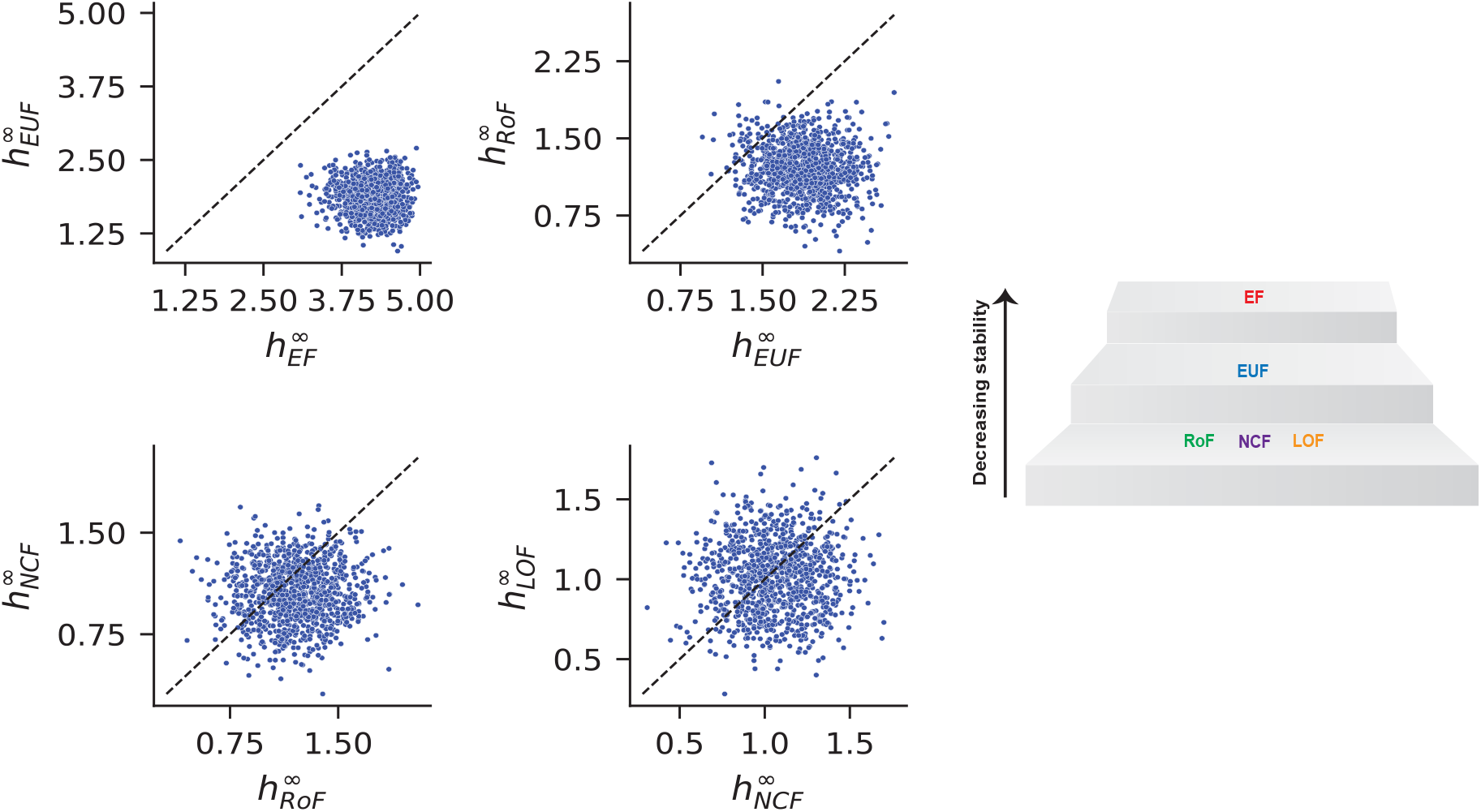
Comparison of mean values of *h*^∞^ when using different classes of BFs on identical networks. For each of the 1000 different networks with *k*-regular (R-R) topology (*k*_*in*_ = *k*_*out*_ = 4, *N* = 12), 10 models were randomly generated, for each of the 5 different ensembles of models (EF, EUF, RoF, NCF and LOF). The values of *h*^∞^ were then averaged over the 10 models and plotted. Each sub-figure provides comparisons across two successive classes of the putative hierarchy. The dashed line is the X=Y boundary. The right panel shows a visual illustration of stability hierarchy when one uses the value of *h*^∞^.

As shown in Fig. 2, such a hierarchy is also evident when using other measures considered to probe stability, whether they are based on damage spreading (*δ, ϕ*), attractor properties (see the 7 observables defined in Methods), or linear and nonlinear MAEs (see Methods). Indeed, these observables show a monotonic trend along the hierarchy. For instance the number of attractors decreases along the hierarchy while the fraction of fixed point attractors increases, both observables behaving as expected when dynamical systems become more stable. In all cases we see that the classes RoF, NCF and LOF have very similar distributions while the EF and EUF stand apart. Given the values of *h*^∞^ for EF and EUF classes, it is evident their associated dynamics is significantly more chaotic than the ones arising when using the other classes of BFs (see *h*^∞^ = 1 boundary separating chaotic from stable dynamics in Fig. 2(i)).

**FIG. 2.**
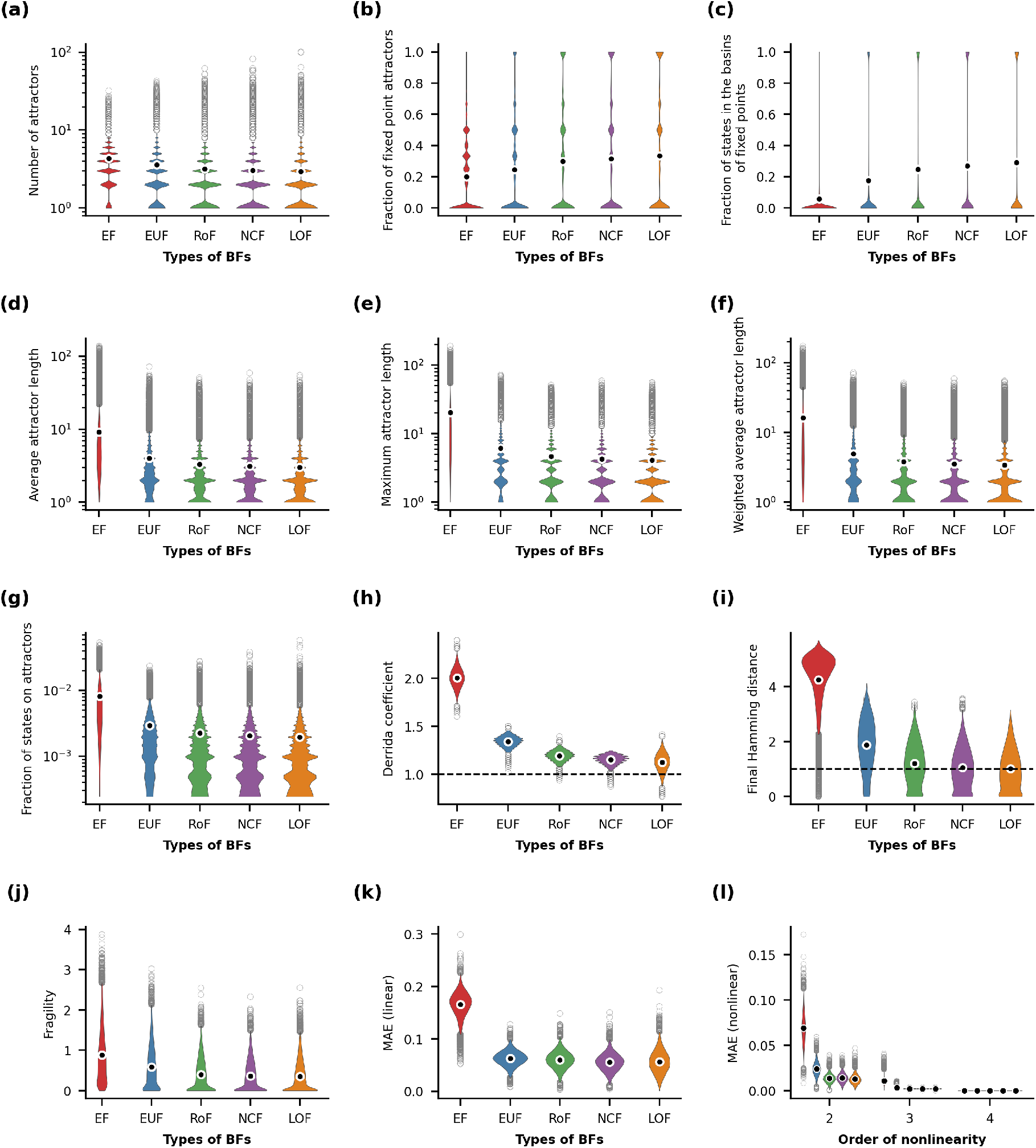
Distributions of stability observables when using different classes of BFs. Within each sub-figure, the violins represent the distributions of the corresponding observable according to the EF, EUF, RoF, NCF and LOF class used for the BFs. For each of the 1000 different networks with R-R topology (*k*_*in*_ = *k*_*out*_ = 4, *N* = 12), 1000 models were generated for the observables in (a)-(g) and 10 models were generated for the observables in (h)-(l). The distributions of the observables were then plotted. Outliers of each distribution are shown as gray rings while the means are shown as black dots encircled with white rings.

In practice, computing either *h*^∞^ or *ϕ* requires iterating the dynamics a very large number of times, and so it is reassuring that other easier-to-compute measures provide similar results. A common proxy for *h*^∞^ is the Derrida coefficient *δ* which measures the damage spreading after a single time step (with *δ* = 1 separating the chaotic and stable dynamical behaviors). In Fig. 2(h), we see that *δ* has distributions with particularly low overlaps when comparing our 5 classes of BFs. To probe this property, we show in SI Fig. S1 the analog of Fig. 1 upon using *δ*. There we see that, for *δ*, one has the more extended hierarchy *δ*_*EF*_ *> δ*_*EUF*_ *> δ*_*RoF*_ *>* (*δ*_*NCF*_ or *δ*_*LOF*_) but where there is no systematic ordering between the NCF and LOF classes. The low overlap of distributions is also visible in SI Fig. S2(a) which displays the joint values of *h*^∞^ and *δ*, and moreover, this observation is also evident from a comparison of the marginal distributions displayed as box plots. Unexpectedly, the correlation between *h*^∞^ and *δ* is relatively weak. Furthermore, while *h*^∞^ is directly tied to the long-term dynamics of the system and depends on the network connectivity, *δ* depends only on the BFs. In fact, *δ* is identical to the so called *average network sensitivity* which does not capture information about the network connectivity [26, 42]. Based on our observations, using *δ* as a proxy for *h*^∞^ is not satisfactory. Interestingly, it may be preferable to use for instance the weighted average attractor length *L*_*W*_ (see Eq. 9) that has a higher correlation with *h*^∞^ and provides similar hierarchy prediction (see SI Fig. S2(b)).

SI Figures S3-S8 reveal that the monotonic trend for *δ, h*^∞^ and *ϕ* as observed in Fig. 2(h)-(j) are consistent irrespective of the value of *k* and the type of topology. For the 7 observables associated to attractors, we see a monotonic trend as shown in Fig. 2(a)-(g). Depending on the observable considered, the trend is either monotone increasing or decreasing. For the two observables “Fraction of fixed point attractors” and “Fraction of states in the basins of fixed points”, the trend is monotone increasing, whereas for the other 5 observables the trend is decreasing. But, in all cases, progressing from EF to EUF to other classes results in increasing stability, i.e., one satisfies the stability hierarchy EF *<* EUF *<* RoF *<* NCF *<* LOF. SI Figures S9-S49 illustrate the distributions of various observables associated to the attractors, the Jensen-Shannon distances between different distribution pairs, and p-values corresponding to the one-sided Kolmogorov-Smirnov (K-S) test. From these plots, it is evident that for all 7 observables associated to attractors, the monotonic trend holds consistently across all values of *N, k* and topologies considered. In each case, for every pair of distributions, we have computed the Jensen-Shannon distances and conducted the one-sided K-S test. Notably, the extremely low p-values from the K-S test in each case strongly support the stability hierarchy. When considering the Jensen-Shannon distances between the distributions for any of the 7 observables associated to attractors, we obtain smaller values for any pair among the RoF, NCF and LOF classes compared to those involving EF or EUF classes. Further, both the linear and nonlinear MAEs exhibit a consistent hierarchy across varying values of *N, k* and types of topology (see SI Figures S50-S52). For the EF class, we obtain consistently the highest MAE, indicating that this class of logic rules is the least approximable. In contrast, the average of both linear or nonlinear MAEs for other BF classes is found to be lower compared to the EF class across varying *N, k* and topology. Still, we find a slight difference in the obtained hierarchy for linear and nonlinear MAEs. Specifically, for linear MAE the obtained hierarchy is EF *>* EUF *>* RoF *>* LOF *>* NCF, while for quadratic and higher-order MAE the obtained hierarchy is EF *>* EUF *>* RoF *>* NCF *>* LOF, irrespective of *N, k* and topology (see SI Figures S50-S52).

Previous studies have demonstrated that the extent of *canalyzation* of logic rules or BFs can influence the the stability of dynamics [18–22]. Specifically, canalyzation can steer the dynamics of the system towards a more ordered regime. In this work, we have considered 5 classes of functions: EF, EUF, RoF, NCF and LOF. The relationship among these classes is as follows: EUF is a subclass of EF, RoF is a subclass of EUF, and both NCF and LOF are subclasses of RoF. Notably, LOF is a subclass of NCF only for *k* = 2 and 3. For other values of *k*, NCF and LOF are only overlapping. For each of the 5 considered classes, there is a non-null intersection with the class of canalyzing functions (CFs) [2]. However, apart from NCFs, no other class is a proper subset of CF. Our results reveal that imposition of the unateness condition itself on BFs significantly drives the dynamics of BNs away from the chaotic regime. Furthermore, applying the rules from the RoF, NCF and LOF classes leads to much greater stability in the dynamics of RBNs. Although, the values of the stability measures for RoF, NCF and LOF classes are quite similar in most cases, the one-sided K-S test reveals that there is a consistent trend, namely we find the stability ordering RoF *<* NCF *<* LOF. It is important to note that both NCF and LOF are subclasses of RoF, but NCF is a special subclass of CF too. In contrast, there are two types of LOFs (AND-pairs and OR-pairs) that do not always belong to the CF class. Specifically, for *k* = 4 and 5, 25% and 36.36% respectively of the total LOFs are not CFs, rather they are *collectively canalyzing* functions [46]. A *k*-input BF is *collectively canalyzing* if the output of the function becomes fully determined by fixing a specific subset of *i* inputs (1 *< i < k*); however, the output remains undetermined when fewer than *i* inputs are fixed. Furthermore, while we were exploring the *approximability* measure proposed in [39] across different classes of BFs, an independent study [40] appeared which also showed that increased canalyzation in RBNs can explain their increased approximability. On similar lines, we show here that NCFs indeed result in minimum linear MAE, i.e., maximum approximability. However, for quadratic MAE, we find that the minimum value for approximability is achieved by LOFs.

### 3.2. The dependence of stability on network average degree is specific to the different classes of functions

From the earliest works on damage spreading [2, 27], it was apparent that a network’s average degree has a strong influence on the system’s dynamics. Specifically, the higher the average degree, the greater the damage spreading in the network. In Fig. 3, each sub-figure displays the mean values of an observable when using the different BF classes, as a function of *k* (R-R topology with *N* = 12). For random choice of functions (∼ EF class), in almost all cases the stability decreases as *k* increases. These trends are as expected: if one increases the number of inputs to a generic BF, there are more possibilities of changing the input, and thus, of spreading any damage. Consequently, the dynamical behavior of a network is pushed towards the chaotic regime as *k* increases. More importantly, for all *k* values in Fig. 3, the stability hierarchy EF *<* EUF *<* (RoF or NCF or LOF) is maintained. At larger values of *N*, the same pattern continues to hold (see SI Figures S53 and S54). Consider for instance the weighted average attractor length (*L*_*W*_) observable that was found to be well correlated with *h*^∞^. Fig. 4 shows that the stability hierarchy EF *<* EUF *<* (RoF or NCF or LOF) is consistently observed across different values of *N* and *k* investigated here. The sub-figures depicting the mean values of *L*_*W*_ (see Fig. 3(f), SI Figures S53(f) and S54(f)) indicate a general trend of increasing values of *L*_*W*_ with increasing *k* (R-R topology) for the EF class across different values of *N* (12, 16 and 20). However, for the classes EUF, RoF, NCF and LOF, the variation in the mean *L*_*W*_ with increasing *k* is less pronounced. Now, higher *L*_*W*_ is associated with more chaotic dynamics. Thus, increasing *k* pushes the dynamics more towards the chaotic regime in case of EF class (random rules). But, for logic rules in the classes such as RoF, NCF or LOF, increasing *k* has only a small effect on the dynamics. This difference also arises for other observables like average attractor length, maximum attractor length, or fraction of states on attractors. Indeed, for each of these observables, there is a significant change in the mean value with increasing *k* for the EF ensemble but not for the other ensembles (EUF, RoF, NCF and LOF).

**FIG. 3.**
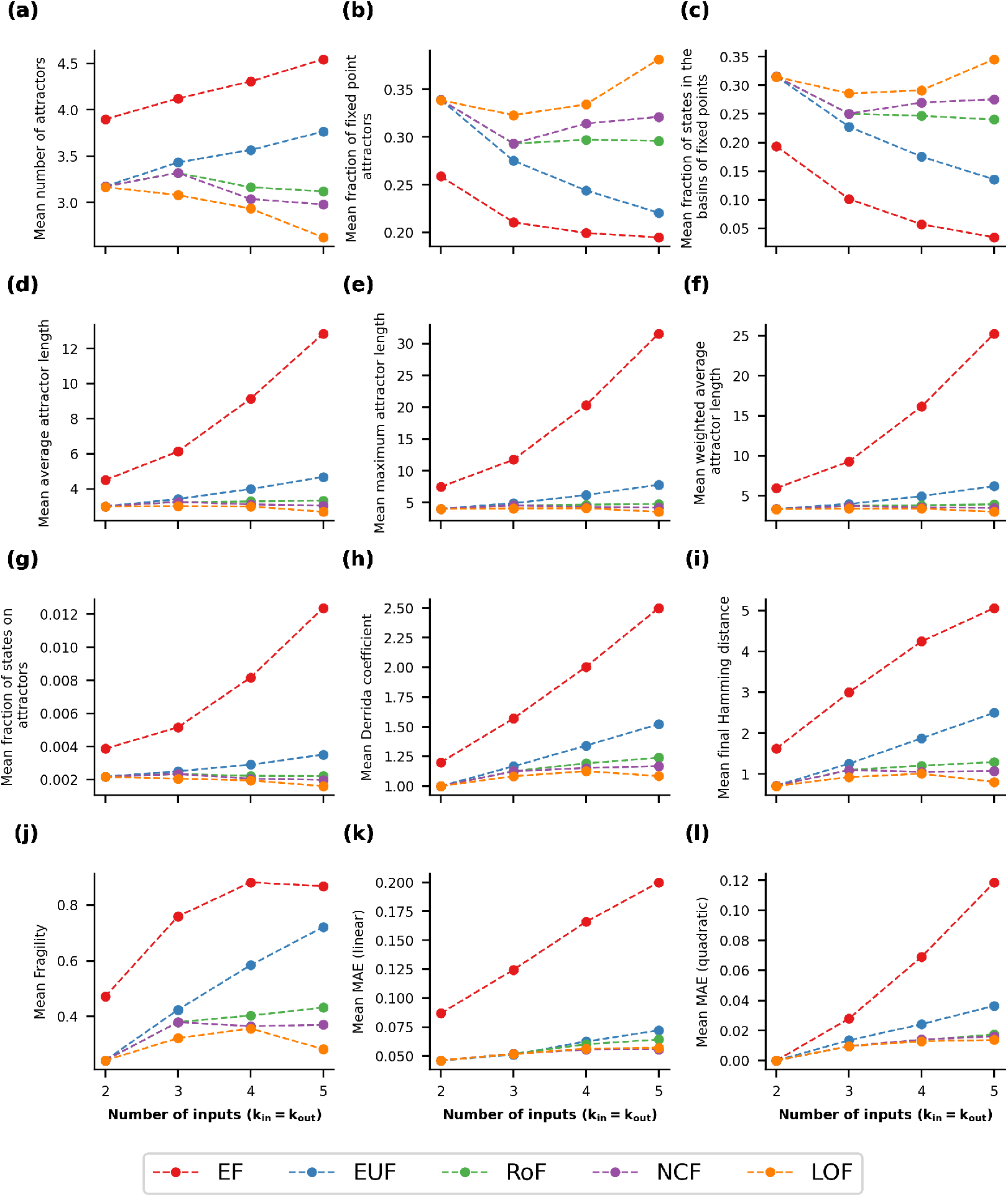
Mean values of our 12 stability observables as a function of *k*_*in*_(= *k*_*out*_) for different classes of BFs, in the case of R-R networks at *N* = 12. Within each sub-figure, the 5 curves correspond to the five different classes of BFs. The data displayed represent the mean value of the corresponding observable at a given value of *k*_*in*_ for the R-R network topology.

**FIG. 4.**
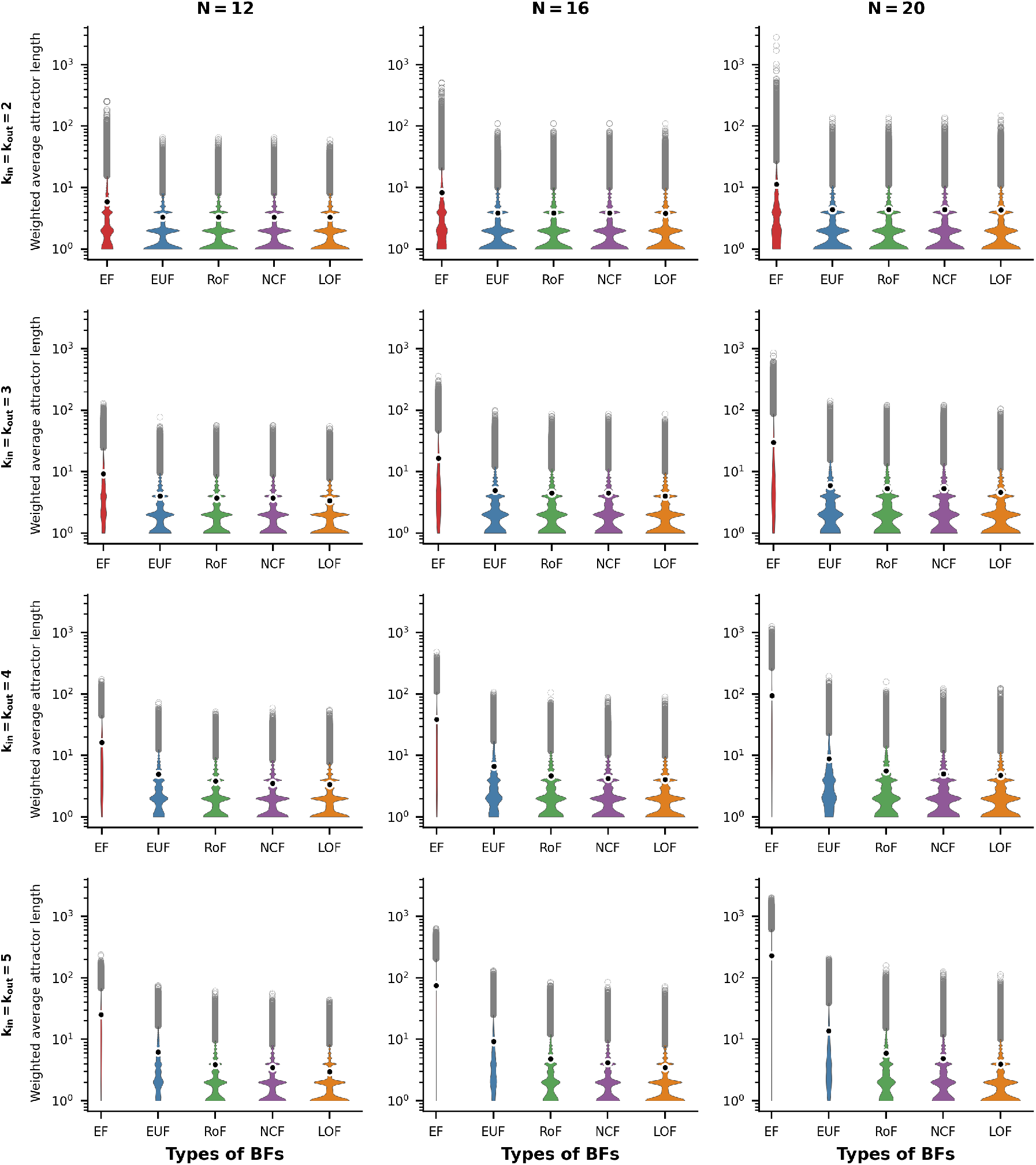
Distributions of *L*_*W*_ (the weighted average attractor length) for R-R networks. The violin plots display the distribution of the weighted average attractor length for the R-R network topology. The sub-figures are organized into 3 columns, corresponding to the network sizes *N* = 12, 16 and 20 respectively. Each row represents a different number of inputs per node, ranging from 2 to 5. Within each sub-figure, there are five violins representing the distributions of the weighted average attractor length for the different ensembles of models, namely EF, EUF, RoF, NCF and LOF. Each violin distribution is based on 10^6^ data points. Outliers of each distribution are shown as gray rings while the means are shown as black dots encircled with white rings. Note the logarithmic scale for the y-axis.

For the observables linear MAE and quadratic MAE, there is a visible trend of increasing mean value with increasing *k* for every ensemble (see Fig. 3, SI Figures S53 and S54). However, the rate of increase is much higher in the EF ensemble than in the other ensembles. We observe a more interesting trend for the observables number of attractors, fraction of fixed point attractors and fraction of states in the basins of fixed point attractors (see Fig. 3, SI Figures S53 and S54). For these observables, the EF and EUF classes show an expected trend: the mean values of the number of attractors increases with increasing *k* while the mean values of the fraction of fixed point attractors and the fraction of states in the basins of fixed point attractors decreases with increasing *k*. However, one sees the opposite trend in the mean values of theses observables with increasing *k* for the LOF and NCF classes. Even though for the LOF and NCF classes the trend is not completely monotonic, it is evident that these classes have a tendency to push the dynamics more towards the ordered regime for higher *k* (at least in terms of these three observables). SI Figures S55-S60 indicate that the above-mentioned observations hold true in the other network topologies we have considered.

From these results, it is evident that the stability of RBNs is significantly influenced by the choice of logic rules as one increases the average degree or connectivity of the network. When random rules are employed, increasing *k* drives the dynamics further towards the chaotic regime. However, by restricting to biologically meaningful classes of BFs namely, EUF, RoF, NCF and LOF (which are prevalent in reconstructed Boolean models of biological systems [22]), we find that there is no significant change in the values of the stability measures with increasing *k*. In other words, our results reinforce the beneficial dynamical consequences of restricting to biologically meaningful classes of BFs like RoFs, NCFs and LOFs when reconstructing and modeling GRNs corresponding to biological processes using the Boolean framework.

### 3.3. The stability hierarchy is robustly maintained across network size

Similarly to the investigation reported in Fig. 3, the mean values of stability observables can be analyzed as a function of network size *N*. Fig. 5 displays the mean values of the stability observables for different BF classes as a function of *N* (R-R topology with *k*_*in*_(= *k*_*out*_) = 4). The nine sub-figures in Fig. 5 display nine different observables; however *h*^∞^ and *ϕ* are omitted from this analysis due the associated computational demand for larger *N*. Additionally, the observable *δ* is excluded as it is solely dependent on the BFs and is independent of network sizes. For the four observables, namely number of attractors, fraction of fixed point attractors, fraction of states in the basins of fixed point attractors and fraction of states on attractors, the stability decreases with increasing *N* regardless of the classes of BFs. In particular, the mean values of the number of attractors increases while the mean values of the fraction of fixed point attractors, the fraction of states in the basins of fixed point attractors and the fraction of states on attractors decreases as *N* increases. Interestingly, we observe a distinct trend for the observables linked to the attractor lengths. For the EF class, the mean values of the three observables namely, average attractor length, maximum attractor length and weighted average attractor length, exhibit an increasing trend as *N* increases. However, we do not observe any significant change in the mean values of these observables for the BF classes EUF, RoF, NCF and LOF. On the other hand, we observe that the mean values of MAE (linear or quadratic) remain almost invariant with increasing *N* irrespective of the choice of BF class. Note that, for any given value of *N*, the stability hierarchy EF *<* EUF *<* (RoF or NCF or LOF) is consistently maintained as discussed in sub-section 3.1. Moreover, for R-R network topology, the above-mentioned observations hold true for other values of *k* as well (see SI Figures S61-S63). Lastly, the above-mentioned observations also hold true even if one analyzes other network topologies (see SI Figures S64-S71).

**FIG. 5.**
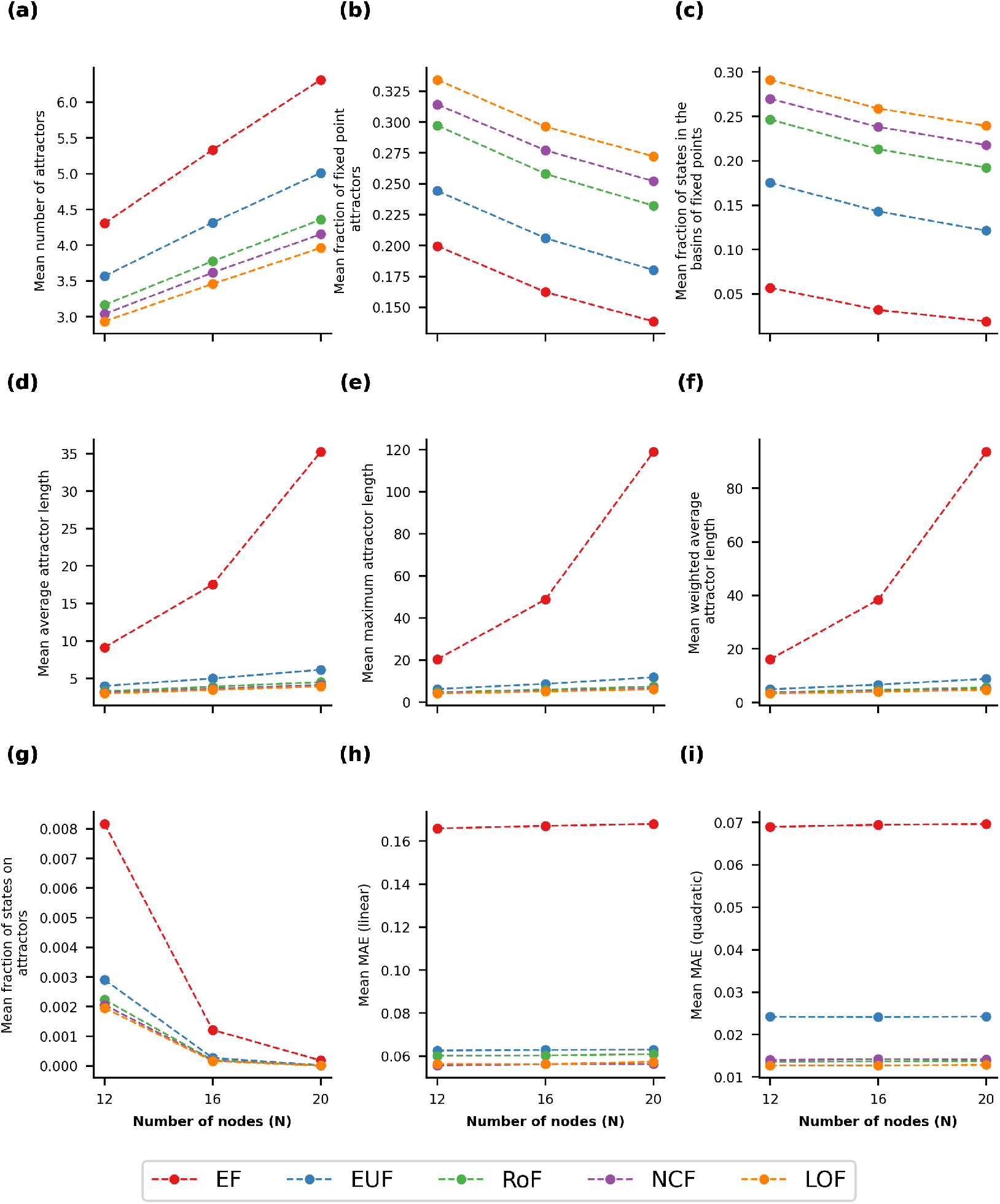
Mean of the stability observables for different values of *N* in the case of the R-R topology with *k*_*in*_ = *k*_*out*_ = 4. Within each sub-figure, the 5 curves correspond to the 5 different classes of BFs. The data displayed represent the mean value of the corresponding observable at a given value of *N* for the R-R network topology with *k*_*in*_ = *k*_*out*_ = 4

Previous research on RBNs has investigated how observables such as the number of attractors or the typical length of attractors are influenced by the increase in BN size [11, 15–17]. To the best of our knowledge, this is the first systematic study on five classes of BFs to show that the stability hierarchy among different classes of functions (EF *<* EUF *<* (RoF or NCF or LOF)) is consistent across different BN sizes. However, with growing BN size, there is a significant influence of the choice of logic rules on the stability of BNs. Furthermore, the observed effect on the stability can vary depending on which observable one considers. For example, with increasing BN size, the observable “number of attractors” shows a very similar trend across different classes of BFs. In contrast, the observables associated with the length of attractors show a significantly different trend for the biologically meaningful functions (EUF, RoF, NCF and LOF) in comparison to the case of random functions (EF). With EF rules, the lengths of the attractors (whether average, weighted average, or maximum) grow very fast with increasing BN size and push the dynamics towards the chaotic regime. In contrast, the use of biologically meaningful functions prevents the typical cycle lengths from going very high. Therefore, these results provide additional support for restricting update rules to the biologically meaningful classes RoFs, NCFs and LOFs when reconstructing and modeling GRNs corresponding to biological processes using the Boolean framework.

## 4. CONCLUSION

Since the advent of RBNs to model GRNs [8–12], much work has focused on the dynamics of RBNs to determine which characteristics of such systems ensure stability or robustness [18–21]. In particular, various measures have been introduced that quantify in different ways the degree of stability achieved in these discrete dynamical systems [26, 39, 41]. The aim of our work was to comprehensively and holistically analyze the effects of both network structure and logical rules on the dynamics of RBNs. We considered both aspects using many different stability measures ranging from damage spreading (see section 2.3) to various attractor properties (see section 2.4). We have also utilized the recently introduced measure of approximability (MAE) as described in section 2.5.

We find that all stability measures follow the expected hierarchy: EF *<* EUF *<* (RoF or NCF or LOF), in the order of least to most stable. That hierarchy is particularly robust, holding irrespective of network size (*N*) and connectivity (*k*), or even of the topological ensemble considered. Unexpectedly, the Derrida Coefficient (*δ*), which is commonly considered as a proxy for the long-term damage spreading measure (*h*^∞^), is found to be quite weakly correlated with *h*^∞^ and thus may not be a satisfactory substitute to infer long-term damage spreading dynamics. In contrast, we find that the weighted average attractor length (*L*_*W*_) is quite highly correlated with *h*^∞^, so it can be used when necessary as a substitute for *h*^∞^.

Next we focused on the average connectivity (*k*) of the network. As expected, the system typically gets pushed towards chaotic dynamics as one increases *k*, however the effects are quite variable across the classes of BFs. Specifically, the random class of functions (EFs) seem to be far more affected by the change in *k* compared to the other classes EUF, RoF, NCF and LOF which have been shown to be more biologically meaningful [22]. For some attractor properties (namely the fraction of fixed point attractors and the fraction of states in the basins of fixed point attractors), we find that the overall trend of decrease in stability is not shared by the classes LOF and NCF at high *k*. Nevertheless, the patterns observed are robust, being insensitive to network topology and network sizes.

Lastly, we examined the effects on the dynamics of varying network size (*N*) when considering the network attractor properties and the MAE. There is significant variation depending upon the observable considered. For example, the average number of attractors shows an increasing trend upon increasing the size of the network across all BF classes. In contrast, the observable “average length of attractors” only shows an increasing trend with network size for the EF rules, whilst not showing significant difference for the other classes of BFs. Such class-specific behaviour is similar to that observed when examining the repercussions of varying the average connectivity of the network.

To sum up, we can conclude that the networks with biologically meaningful classes of functions can largely withstand variations to the topology and the structure of the network and still manage to preserve the extent of stability of the system. A important limitation of this work is its restriction to rather modest values of *N* ; indeed many of the observables we considered become computationally intractable at large *N*. Thus, some trends such as the dependence on *k* seem out of reach if one wants to determine the large *N* limit; indeed, large *k* values require large *N* values if finite size effects are to be small. However even at *N* = 20 we have been unable to obtain long-term damage spreading measures such as *h*^∞^ and *ϕ* due to constraints on our computational resources. Hence an immediate improvement would be to find a way to provide reliable measurements or better proxies for larger sized networks and to verify if the trends remain the same. One could also try to explore the effects of performing our analyses under asynchronous [14] dynamics and see if one can still reach similar conclusions.

## ACKNOWLEDGMENTS

Olivier C. Martin and Areejit Samal would like to acknowledge funding from Indo-French Centre for the Promotion of Advanced Research (IFCPAR/CEFIPRA) via Collaborative Scientific Research Programme (CSRP) project number 7004-1. Areejit Samal acknowledges funding from the Department of Atomic Energy, Government of India via Apex project to The Institute of Mathematical Sciences (IMSc), Chennai, India. IPS2 benefits from the support of Saclay Plant Sciences-SPS (ANR-17-EUR-0007). Priyotosh Sil would like to acknowledge IFCPAR/CEFIPRA for support through the award of a Raman-Charpak fellowship. We are thankful to Ajay Subbaroyan for the insightful feedback, and Priyotosh Sil is also grateful to him for computational assistance.

## AUTHOR CONTRIBUTION

Designed the research: P.S., S.M., O.C.M., A.S.; Performed the research: P.S., S.M., O.C.M., A.S.; Performed the computations: P.S., S.M.; Wrote the paper: P.S., S.M., O.C.M., A.S.

