## Supplementary material for "Choices of regulatory logic class modulate the dynamical regime in random Boolean networks": SI Fig

### SUPPLEMENTARY INFORMATION
### FOR
### Choices of regulatory logic class modulate the dynamical regime in random Boolean networks

Priyotosh Sil,[1,2] Suchetana Mitra,[3,4] Olivier C. Martin,[3,4,*] and Areejit Samal[1,2,*]

[1] *The Institute of Mathematical Sciences (IMSc), Chennai 600113, India*
[2] *Homi Bhabha National Institute (HBNI), Mumbai 400094, India*
[3] *Université Paris-Saclay, CNRS, INRAE, Univ Evry, Institute of Plant Sciences Paris-Saclay (IPS2), 91405 Orsay, France*
[4] *Université Paris-Cité, CNRS, INRAE, Institute of Plant Sciences Paris-Saclay (IPS2), 91405 Orsay, France*

* To whom correspondence should be addressed:


#### SUPPLEMENTARY FIGURES

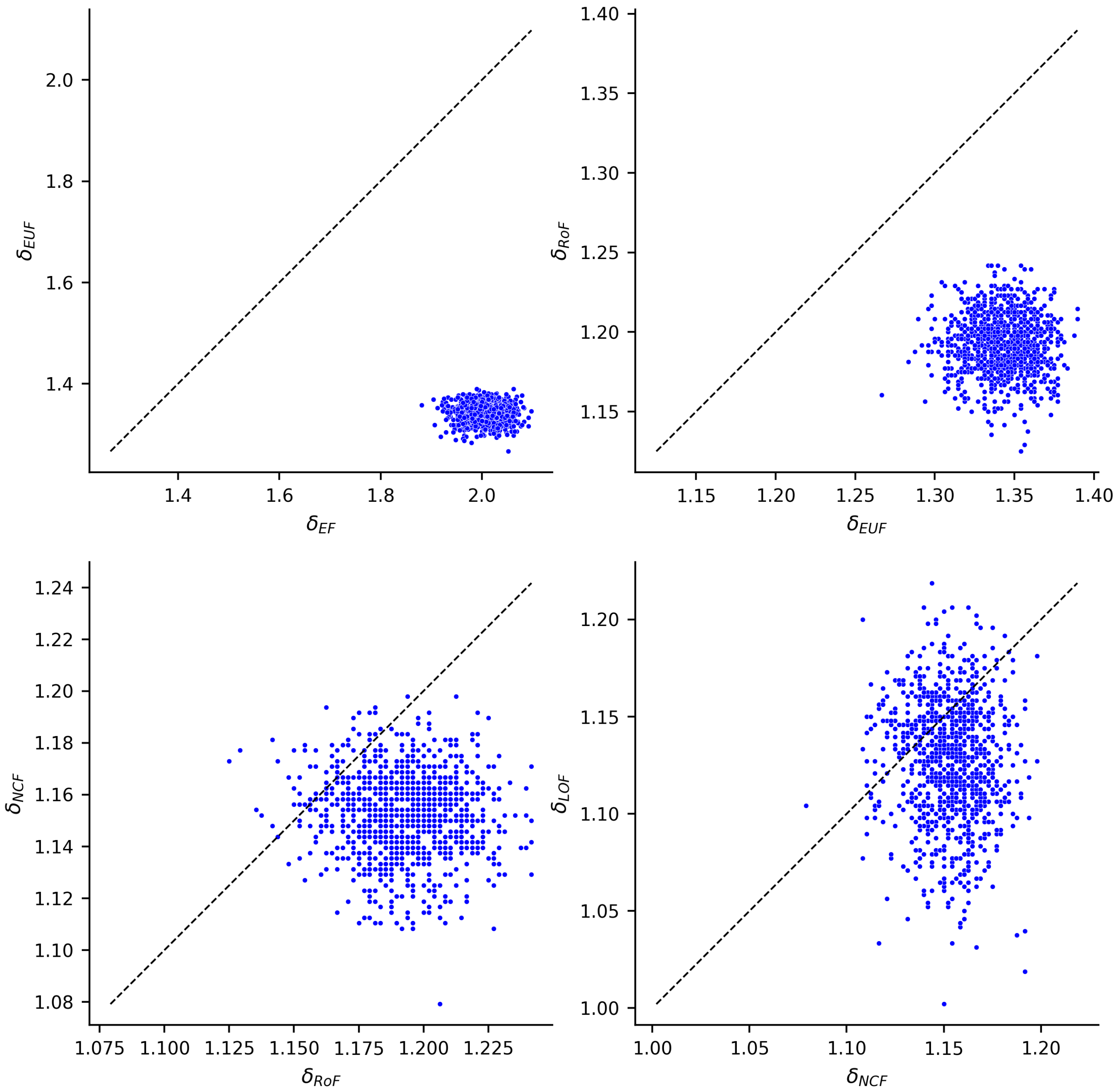


FIG. S1. **Comparison of mean values of $\delta$ when using different classes of BFs on identical networks.** For each of the 1000 different networks with $k$-regular (R-R) topology ($k_{in} = k_{out} = 4$, $N = 12$), 10 models were randomly generated, for each of the 5 different ensembles of models (EF, EUF, RoF, NCF and LOF). The values of $\delta$ were then averaged over the 10 models and plotted. Each sub-figure provides comparisons across two successive classes of the putative hierarchy. The dashed line is the X=Y boundary.

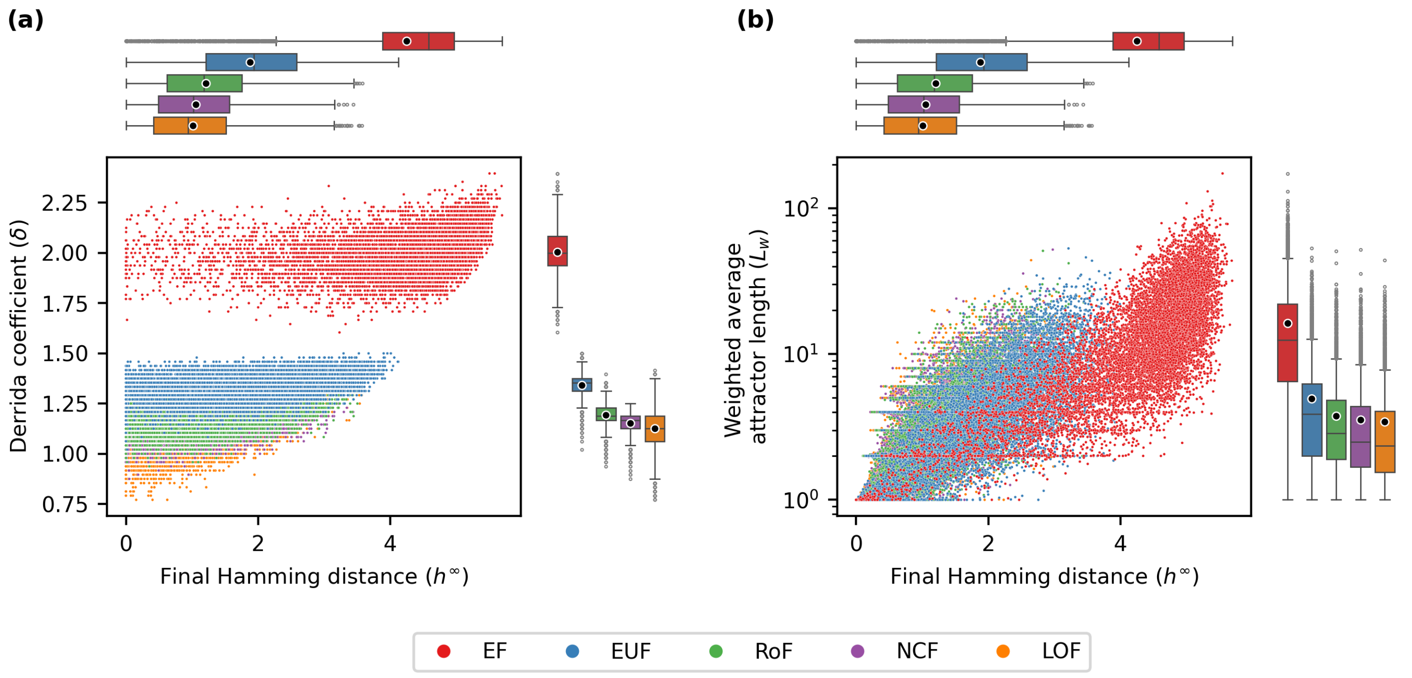


FIG. S2. **Comparing the distributions of $h^{\infty}$ against $\delta$ and $h^{\infty}$ against $L_w$.** For each of the 1000 different networks with $k$-regular (R-R) topology ($k_{in} = k_{out} = 4$, $N = 12$), 10 models were randomly generated, for each of the 5 different ensembles of models (EF, EUF, RoF, NCF and LOF). The quantities $h^{\infty}$, $\delta$ and $L_W$ were computed for each instance. The sub-figure **(a)** illustrates the joint values of $\delta$ and $h^{\infty}$ along with their marginal distributions for different ensembles. Similarly, the sub-figure **(b)** presents the joint values of $\delta$ and $L_W$ across ensembles.

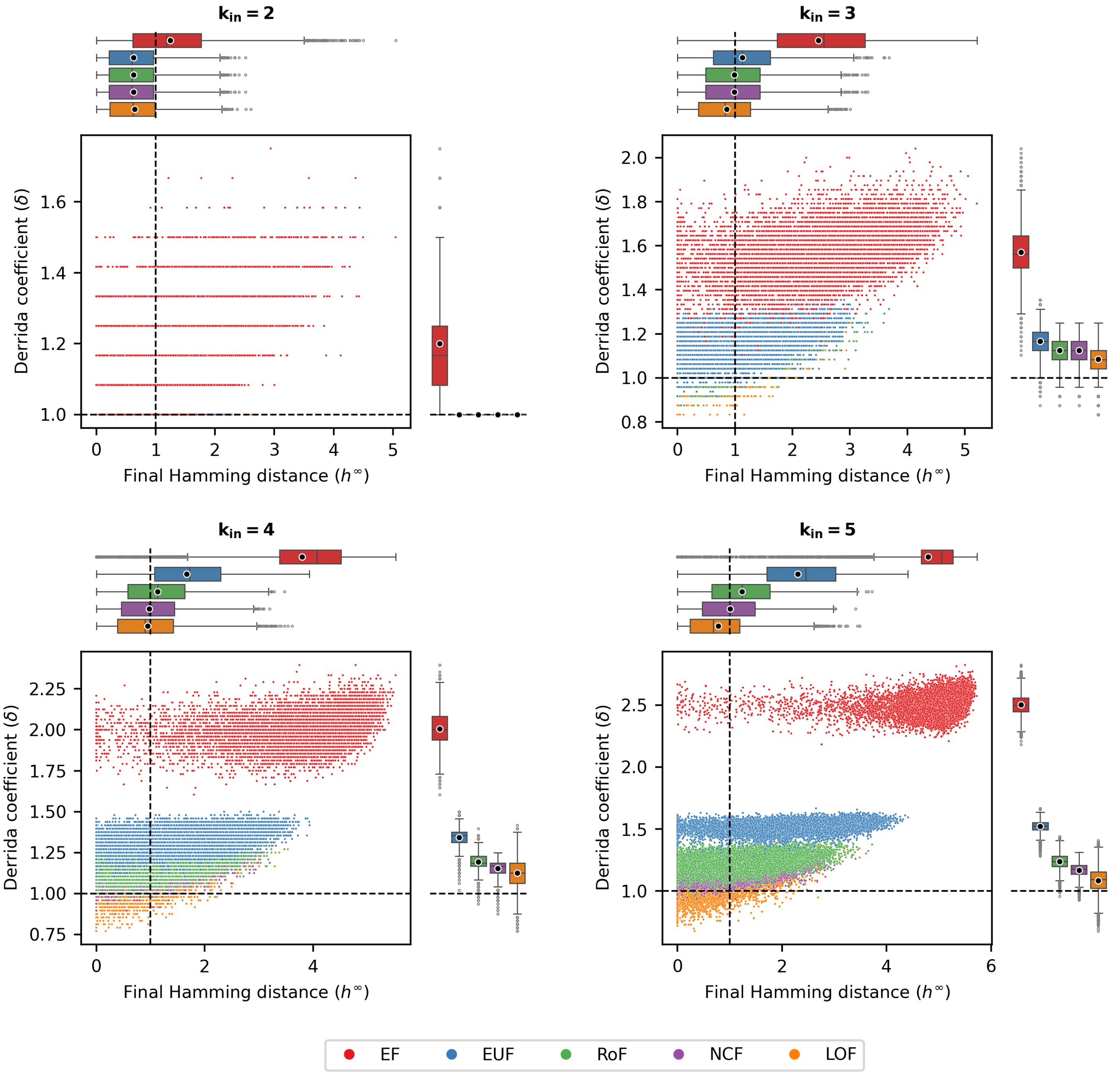


FIG. S3. **The distributions of $h^\infty$ against $\delta$ for R-P networks.** For each of the 1000 different networks with R-P topology ($N = 12$), 10 models were randomly generated, for each of the 5 different ensembles of models (EF, EUF, RoF, NCF and LOF). The quantities $h^\infty$, $\delta$ were computed for each instance. Each sub-figure corresponds to a different number of inputs per node, ranging from 2 to 5. In each sub-figure, the horizontal and vertical dashed black lines correspond to $\delta = 1$ and $h^\infty = 1$ respectively.

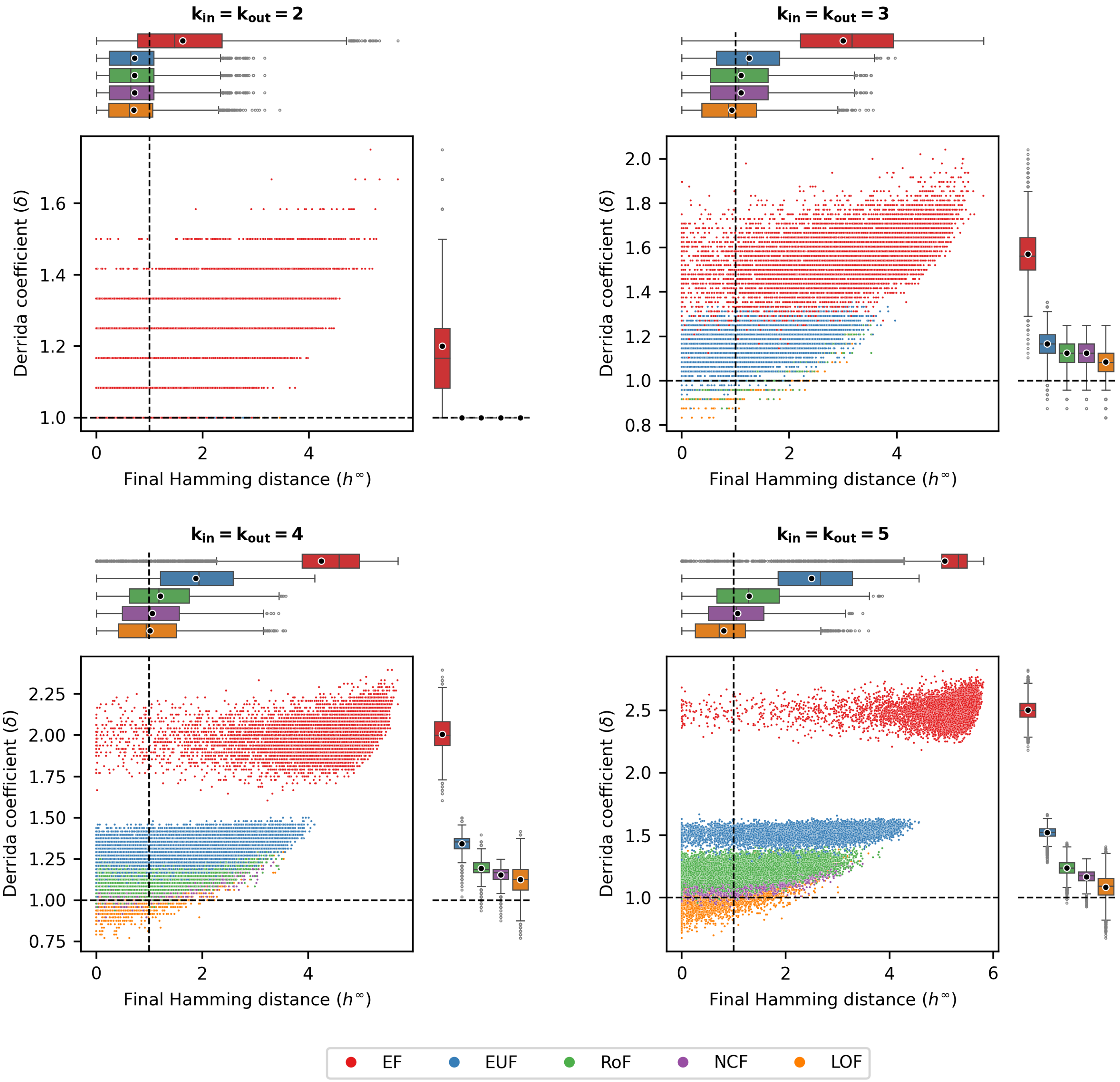


FIG. S4. **The distributions of $h^\infty$ against $\delta$ for R-R networks.** For each of the 1000 different networks with R-R topology ($N = 12$), 10 models were randomly generated, for each of the 5 different ensembles of models (EF, EUF, RoF, NCF and LOF). The quantities $h^\infty$, $\delta$ were computed for each instance. Each sub-figure corresponds to a different number of inputs per node, ranging from 2 to 5. In each sub-figure, the horizontal and vertical dashed black lines correspond to $\delta = 1$ and $h^\infty = 1$ respectively.

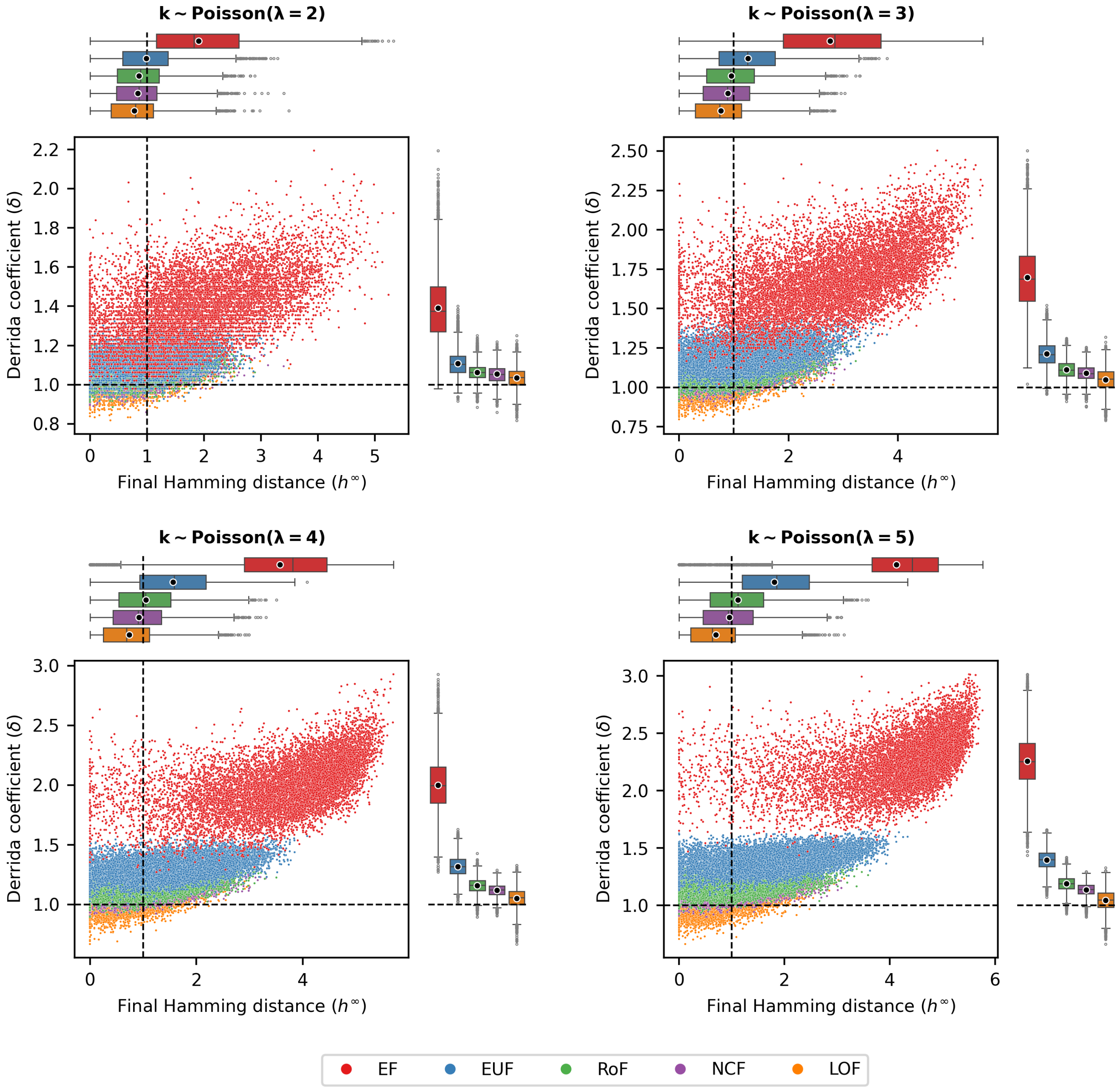


FIG. S5. **The distributions of $h^{\infty}$ against $\delta$ for P-P networks.** For each of the 1000 different networks with P-P topology ($N = 12$), 10 models were randomly generated, for each of the 5 different ensembles of models (EF, EUF, RoF, NCF and LOF). The quantities $h^{\infty}$, $\delta$ were computed for each instance. Each sub-figure corresponds to a different average number of inputs per node, with the degrees drawn from Poisson distributions with means 2, 3, 4 and 5. In each sub-figure, the horizontal and vertical dashed black lines correspond to $\delta = 1$ and $h^{\infty} = 1$ respectively.

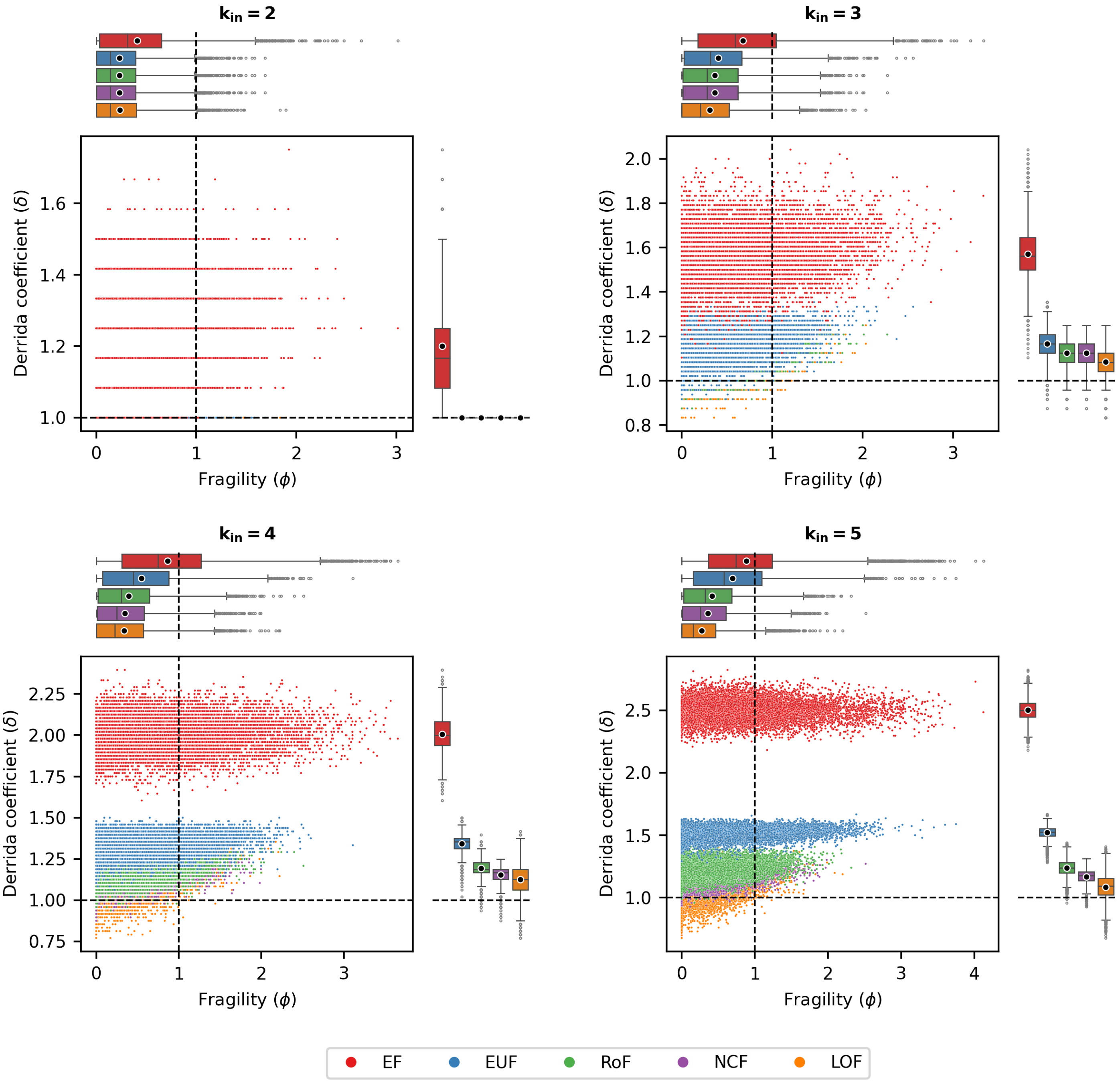


FIG. S6. **The distributions of $\phi$ against $\delta$ for R-P networks.** For each of the 1000 different networks with R-P topology ($N = 12$), 10 models were randomly generated, for each of the 5 different ensembles of models (EF, EUF, RoF, NCF and LOF). The quantities $\phi$, $\delta$ were computed for each instance. Each sub-figure corresponds to a different number of inputs per node, ranging from 2 to 5. In each sub-figure, the horizontal and vertical dashed black lines correspond to $\delta = 1$ and $\phi = 1$ respectively.

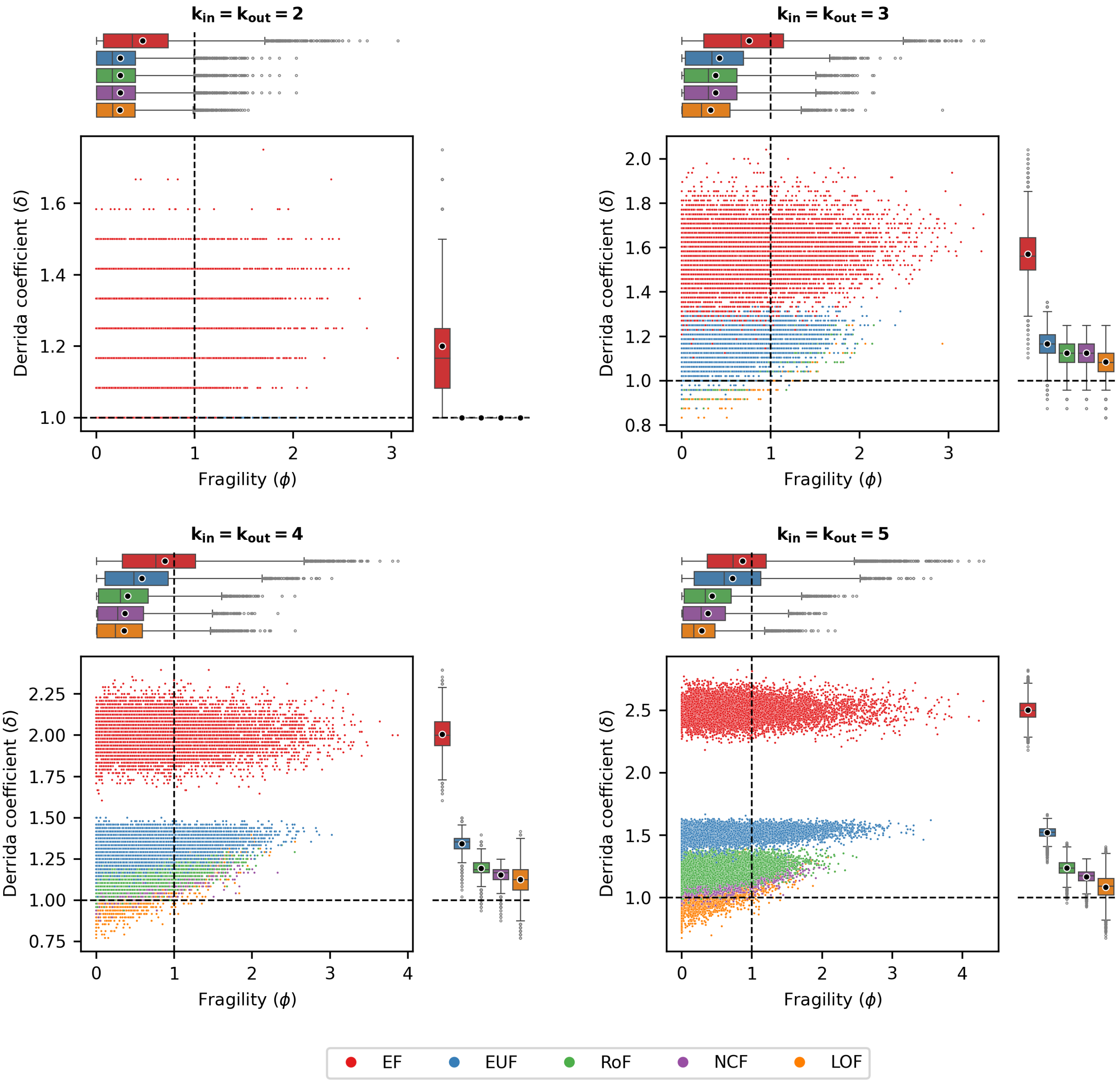


FIG. S7. **The distributions of $\phi$ against $\delta$ for R-R networks.** For each of the 1000 different networks with R-R topology ($N = 12$), 10 models were randomly generated, for each of the 5 different ensembles of models (EF, EUF, RoF, NCF and LOF). The quantities $\phi$, $\delta$ were computed for each instance. Each sub-figure corresponds to a different number of inputs per node, ranging from 2 to 5. In each sub-figure, the horizontal and vertical dashed black lines correspond to $\delta = 1$ and $\phi = 1$ respectively.

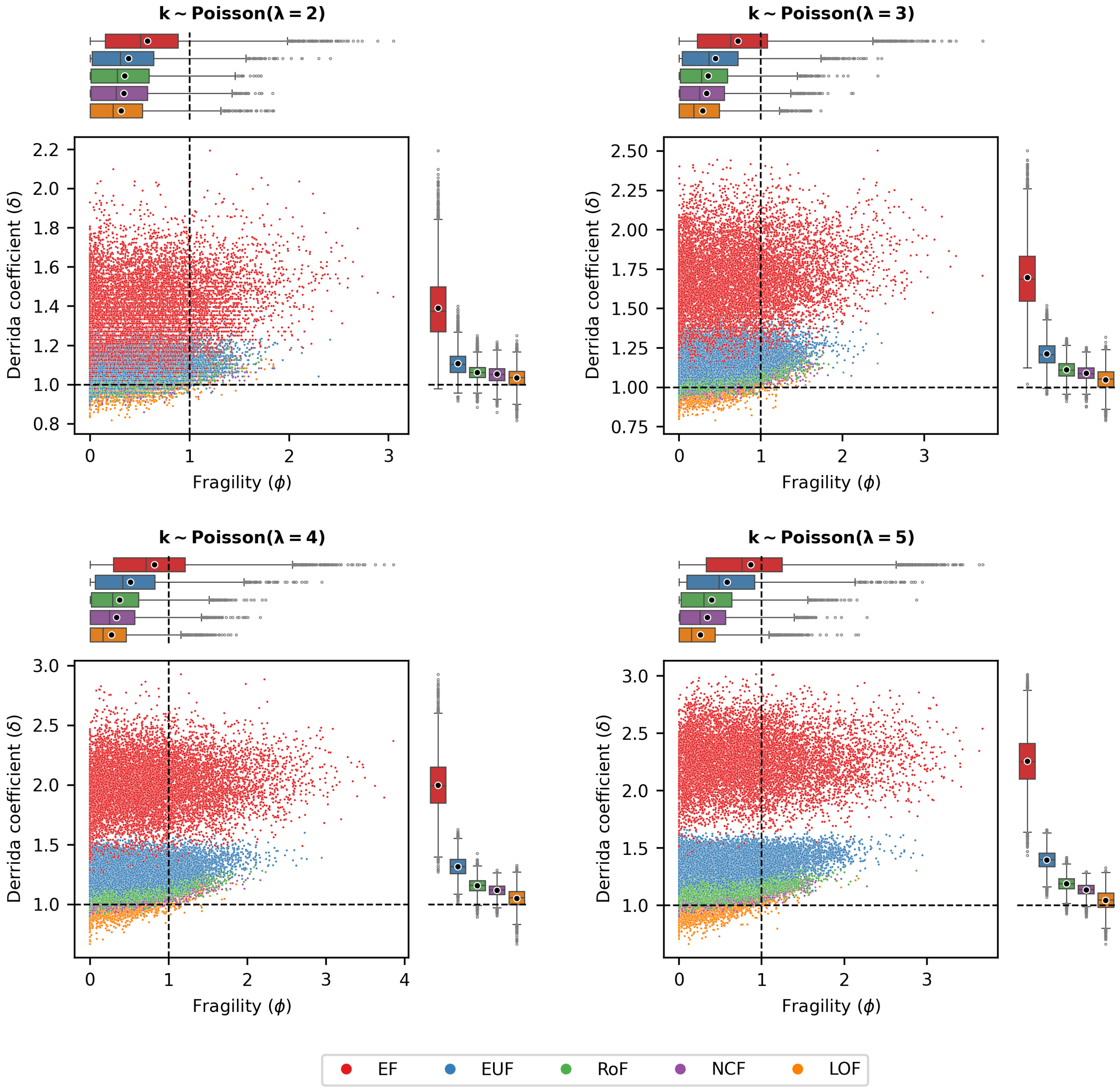


FIG. S8. **The distributions of $\phi$ against $\delta$ for P-P networks.** For each of the 1000 different networks with P-P topology ($N = 12$), 10 models were randomly generated, for each of the 5 different ensembles of models (EF, EUF, RoF, NCF and LOF). The quantities $\phi$, $\delta$ were computed for each instance. Each sub-figure corresponds to a different average number of inputs per node, with the degrees drawn from Poisson distributions with means 2, 3, 4 and 5. In each sub-figure, the horizontal and vertical dashed black lines correspond to $\delta = 1$ and $\phi = 1$ respectively.

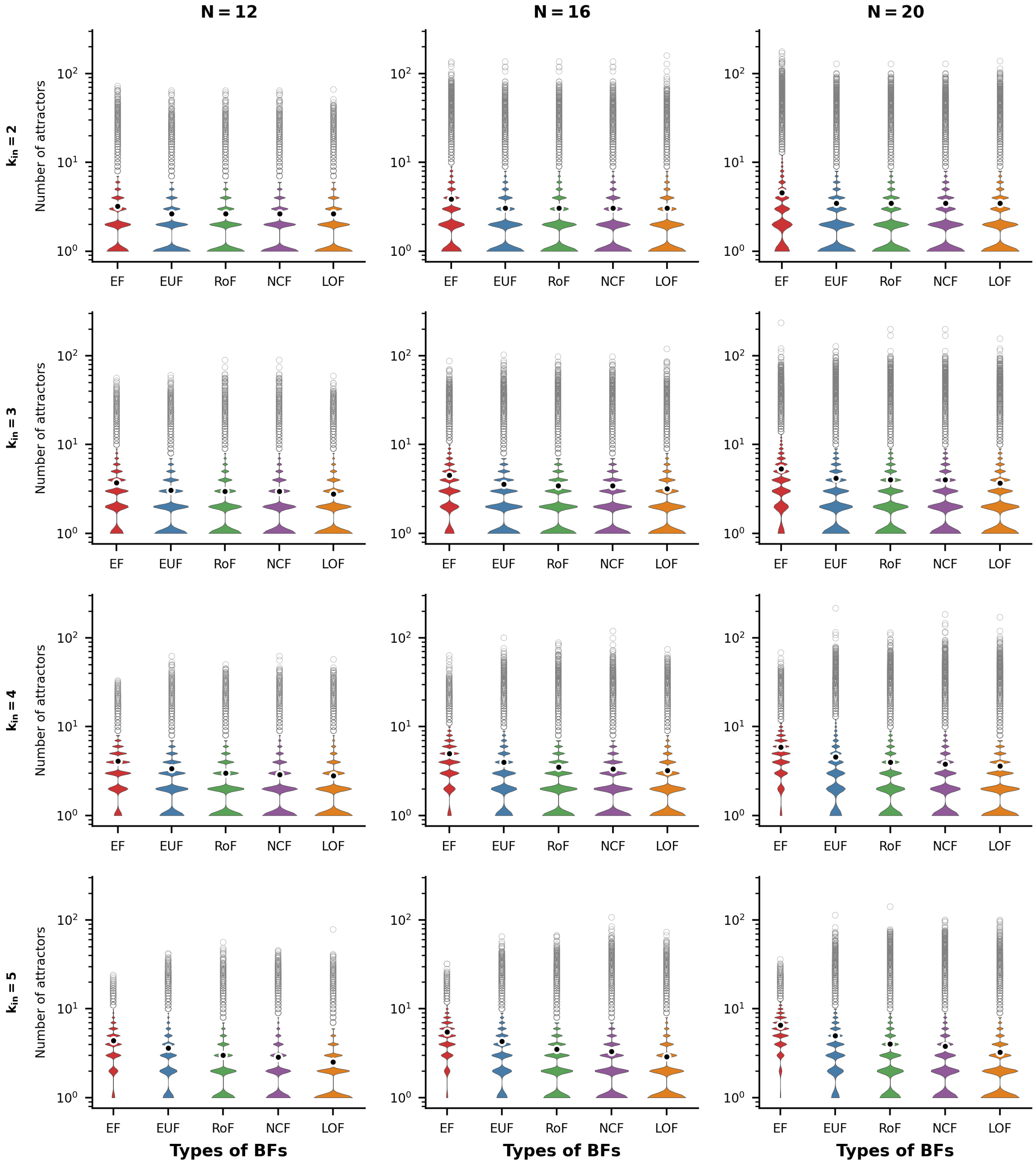


FIG. S9. **Distributions of number of attractors for R-P networks.** The violin plots display the distribution of the number of attractors for the R-P network topology. The sub-figures are organized into 3 columns, corresponding to the network sizes $N = 12, 16$ and 20 respectively. Each row represents a different number of inputs per node, ranging from 2 to 5. Within each sub-figure, there are five violins representing the distributions of the number of attractors for the different ensembles of models, namely EF, EUF, RoF, NCF and LOF. Each violin distribution is based on $10^6$ data points. Outliers of each distribution are shown as gray rings while the means are shown as black dots encircled with white rings. Note the logarithmic scale for the y-axis.

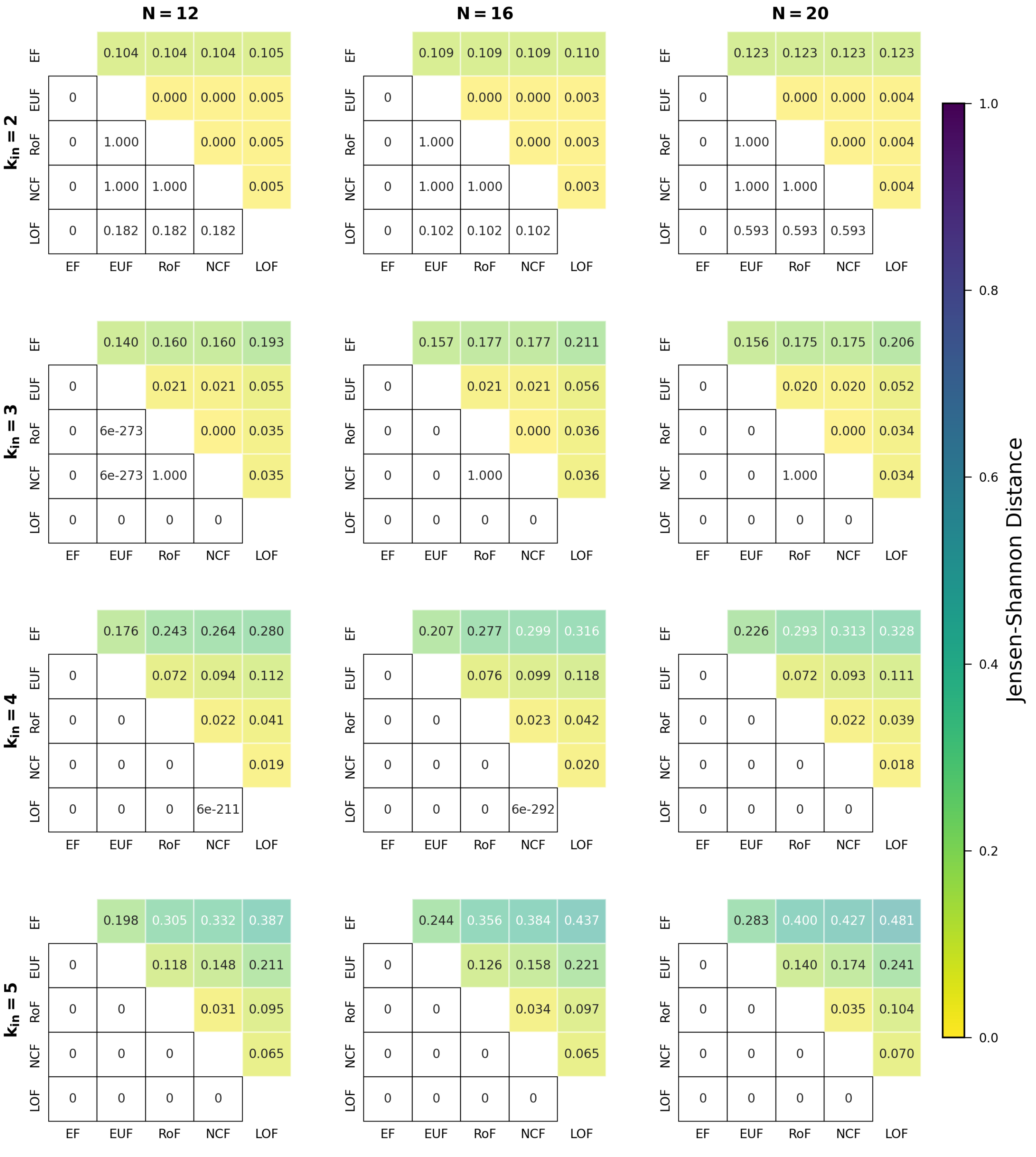

FIG. S10. **Comparing distributions for the number of attractors when using different classes of BFs within the R-P networks.** The different sub-figures are organised according to the values of $k_{in}$ (the in-degree) and $N$ (the number of nodes in the network). At given $k_{in}$ and $N$, we determined the distribution of the number of attractors for each of the 5 classes of BFs. In each sub-figure, the upper triangular part displays the pairwise Jensen-Shannon distances of those distributions. The lower triangular part represents the p-values of the one-sided K-S test (cumulative distribution of the label of column less than that of the row). Entries shown as 0 are actually very small, with values less than $10^{-307}$.

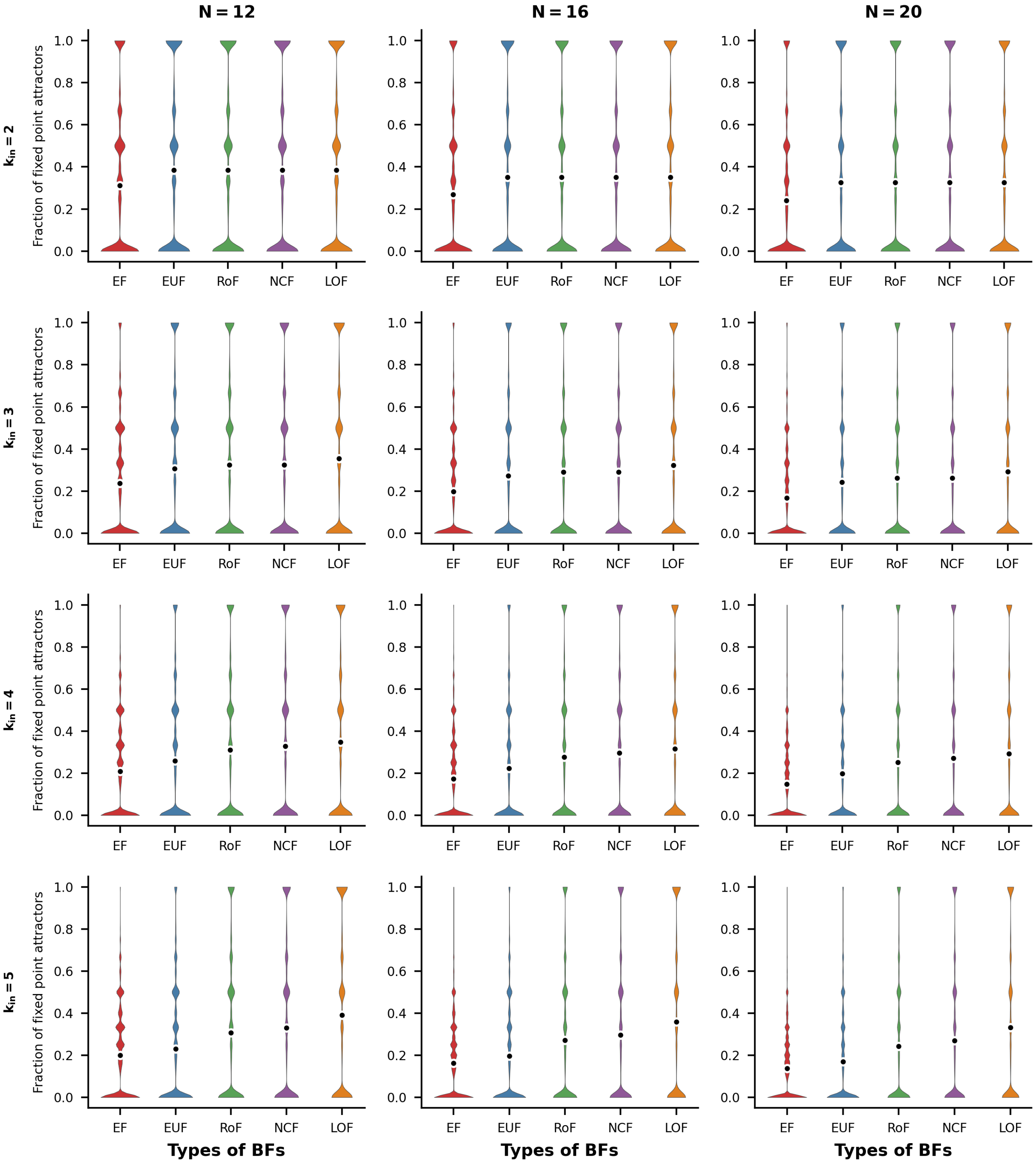


FIG. S11. **Distributions of fraction of fixed point attractors for R-P networks.** The violin plots display the distribution of the fraction of fixed point attractors for the R-P network topology. The sub-figures are organized into 3 columns, corresponding to the network sizes $N = 12, 16$ and 20 respectively. Each row represents a different number of inputs per node, ranging from 2 to 5. Within each sub-figure, there are five violins representing the distributions of the fraction of fixed point attractors for the different ensembles of models, namely EF, EUF, RoF, NCF and LOF. Each violin distribution is based on $10^6$ data points. Outliers of each distribution are shown as gray rings while the means are shown as black dots encircled with white rings.

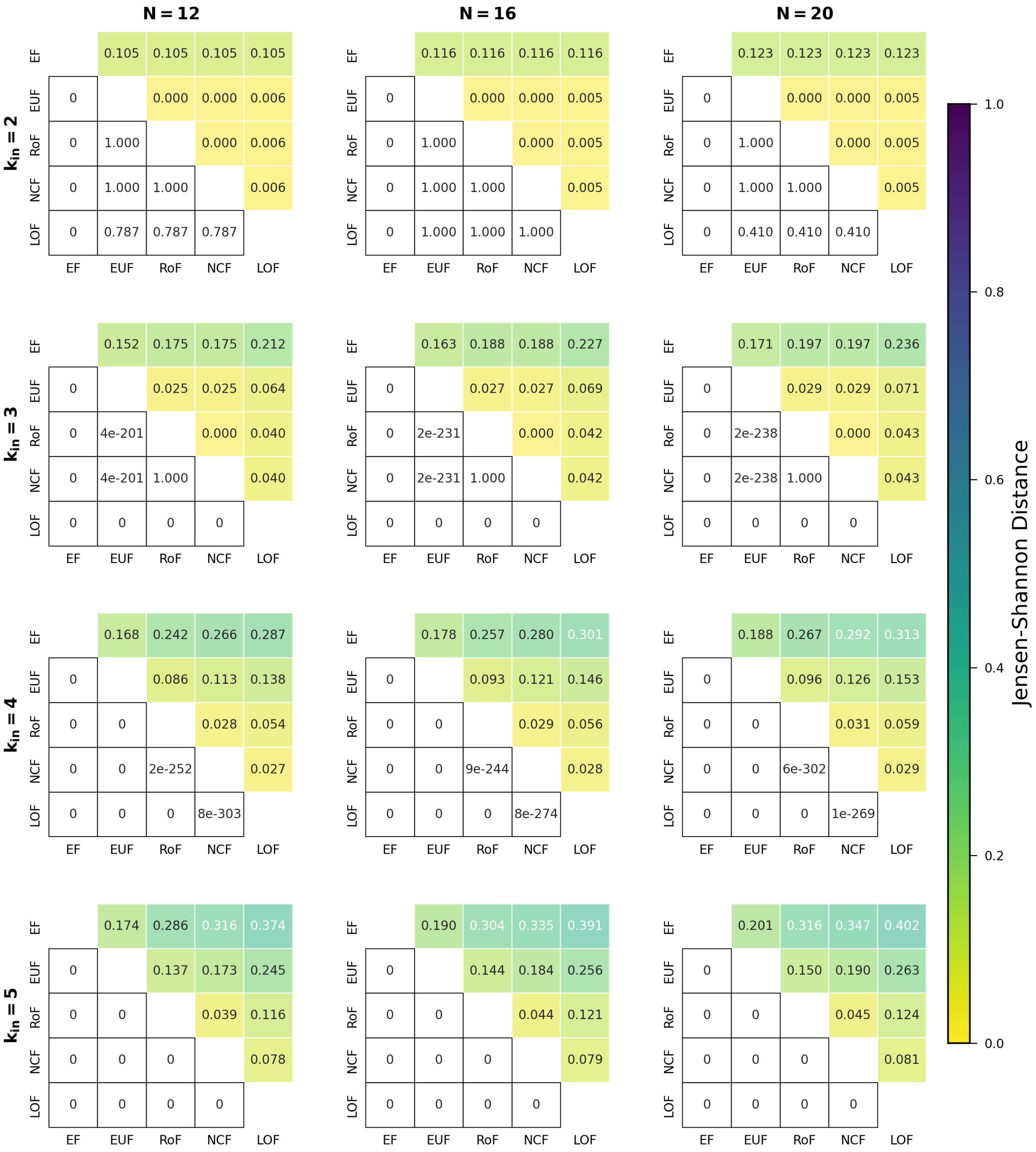


FIG. S12. **Comparing distributions for the fraction of fixed point attractors when using different classes of BFs within the R-P networks.** The different sub-figures are organised according to the values of $k_{in}$ (the in-degree) and $N$ (the number of nodes in the network). At given $k_{in}$ and $N$, we determined the distribution of the fraction of fixed point attractors for each of the 5 classes of BFs. In each sub-figure, the upper triangular part displays the pairwise Jensen-Shannon distances of those distributions. The lower triangular part represents the p-values of the one-sided K-S test (cumulative distribution of the label of column greater than that of the row). Entries shown as 0 are actually very small, with values less than $10^{-307}$.

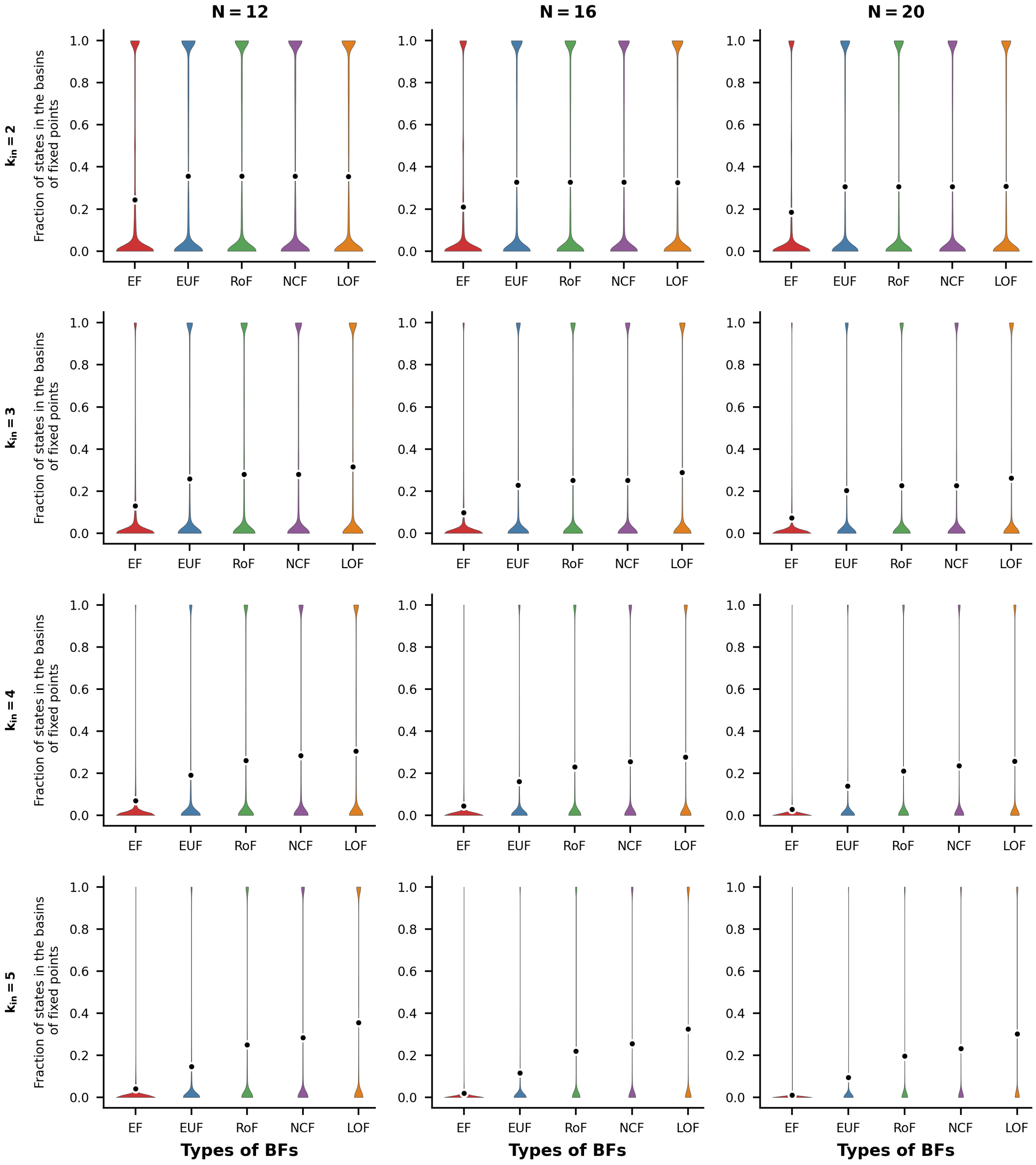


FIG. S13. **Distributions of Fraction of states in the basins of fixed points for R-P networks** The violin plots display the distribution of the fraction of states in the basins of fixed point attractors for the R-P network topology. The sub-figures are organized into 3 columns, corresponding to the network sizes $N = 12, 16$ and 20 respectively. Each row represents a different number of inputs per node, ranging from 2 to 5. Within each sub-figure, there are five violins representing the distributions of the fraction of states in the basins of fixed point attractors for the different ensembles of models, namely EF, EUF, RoF, NCF and LOF. Each violin distribution is based on $10^6$ data points. Outliers of each distribution are shown as gray rings while the means are shown as black dots encircled with white rings.

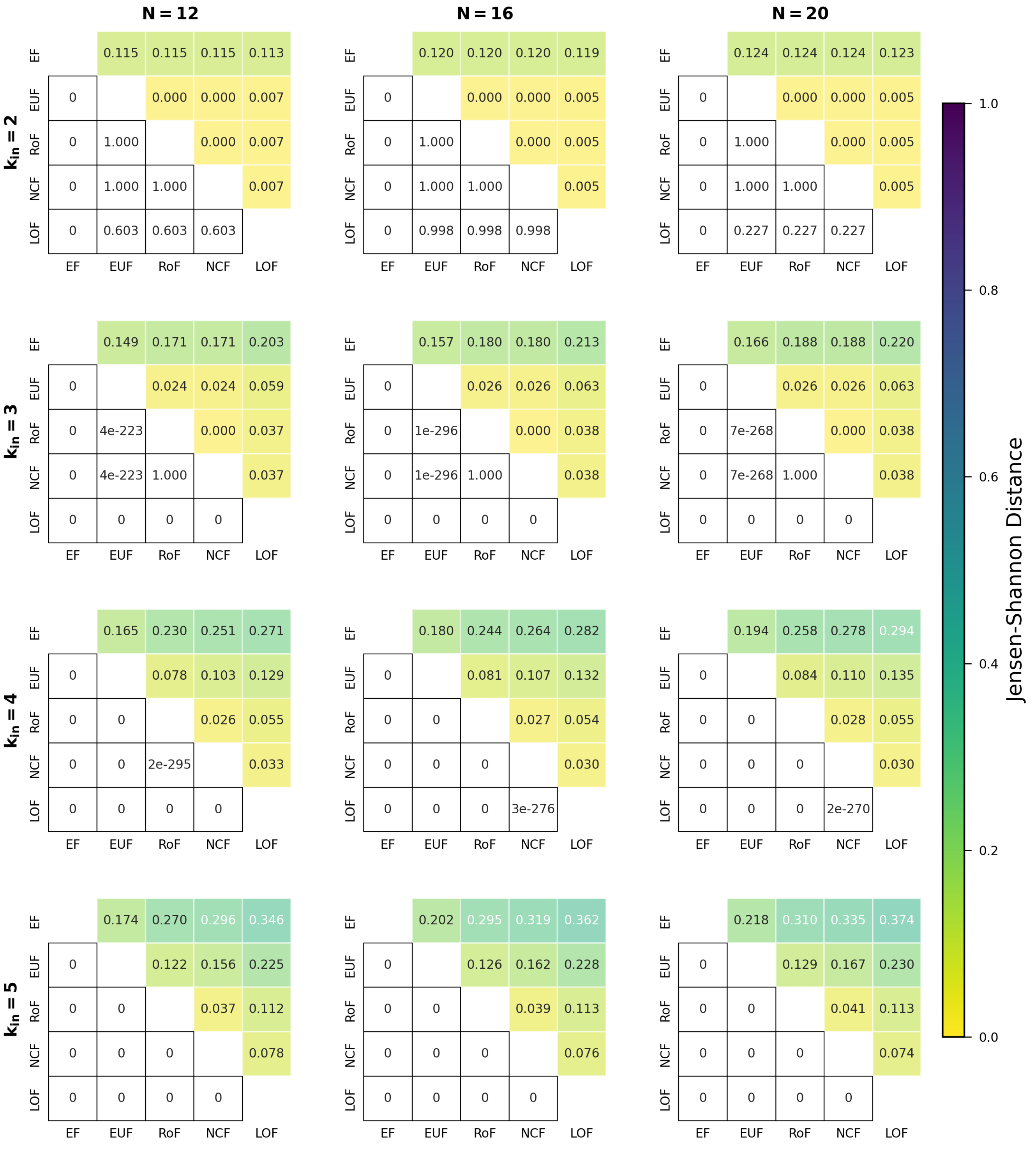


FIG. S14. **Comparing distributions for the fraction of states in the basins of fixed points when using different classes of BFs within the R-P networks.** The different sub-figures are organised according to the values of $k_{in}$ (the in-degree) and $N$ (the number of nodes in the network). At given $k_{in}$ and $N$, we determined the distribution of the fraction of states in the basins of fixed points for each of the 5 classes of BFs. In each sub-figure, the upper triangular part displays the pairwise Jensen-Shannon distances of those distributions. The lower triangular part represents the p-values of the one-sided K-S test (cumulative distribution of the label of column greater than that of the row). Entries shown as 0 are actually very small, with values less than $10^{-307}$.

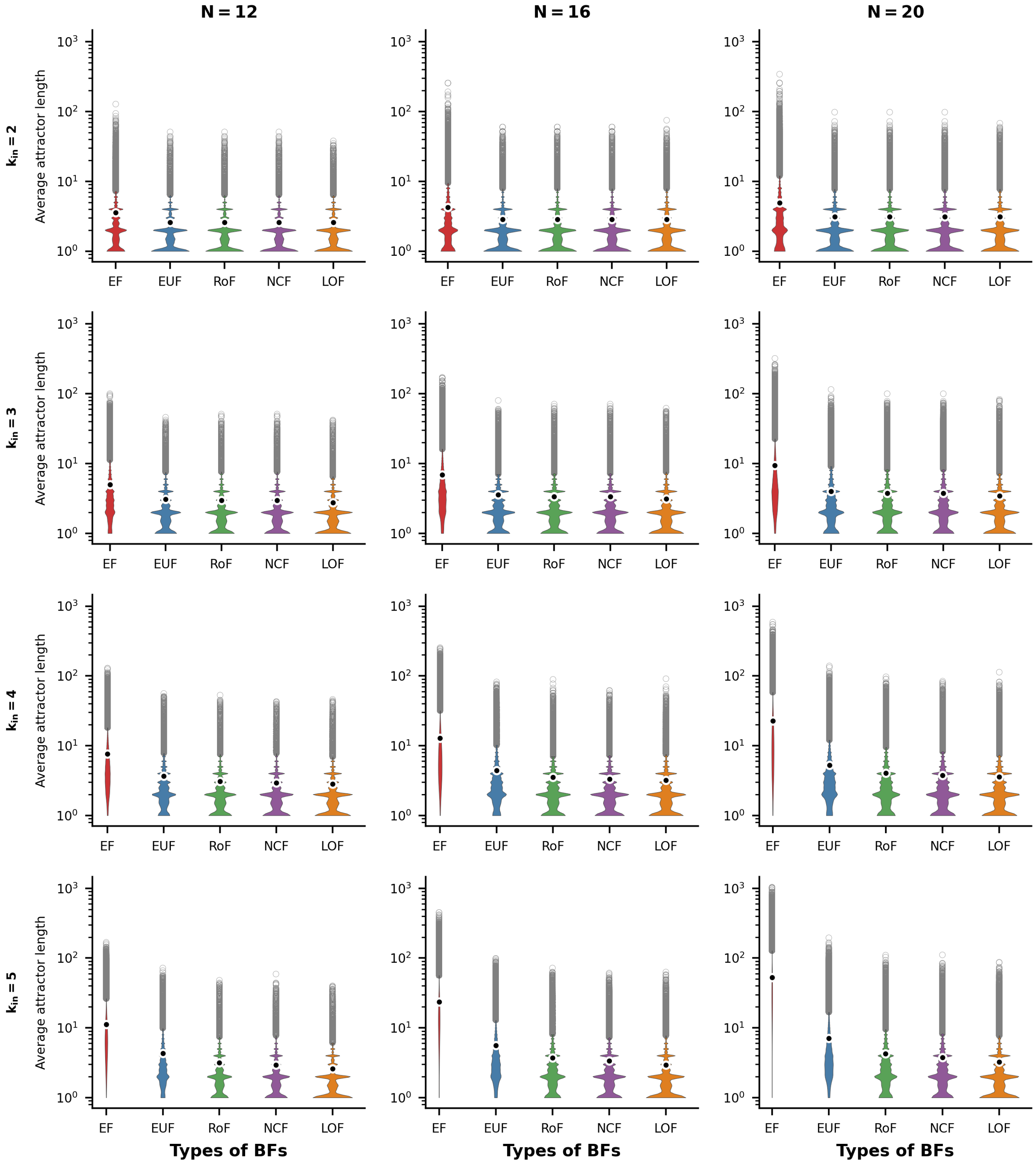


FIG. S15. **Distributions of average attractor lengths for R-P networks.** The violin plots display the distribution of the average attractor length for the R-P network topology. The sub-figures are organized into 3 columns, corresponding to the network sizes $N = 12, 16$ and 20 respectively. Each row represents a different number of inputs per node, ranging from 2 to 5. Within each sub-figure, there are five violins representing the distributions of the average attractor length for the different ensembles of models, namely EF, EUF, RoF, NCF and LOF. Each violin distribution is based on $10^6$ data points. Outliers of each distribution are shown as gray rings while the means are shown as black dots encircled with white rings. Note the logarithmic scale for the y-axis.

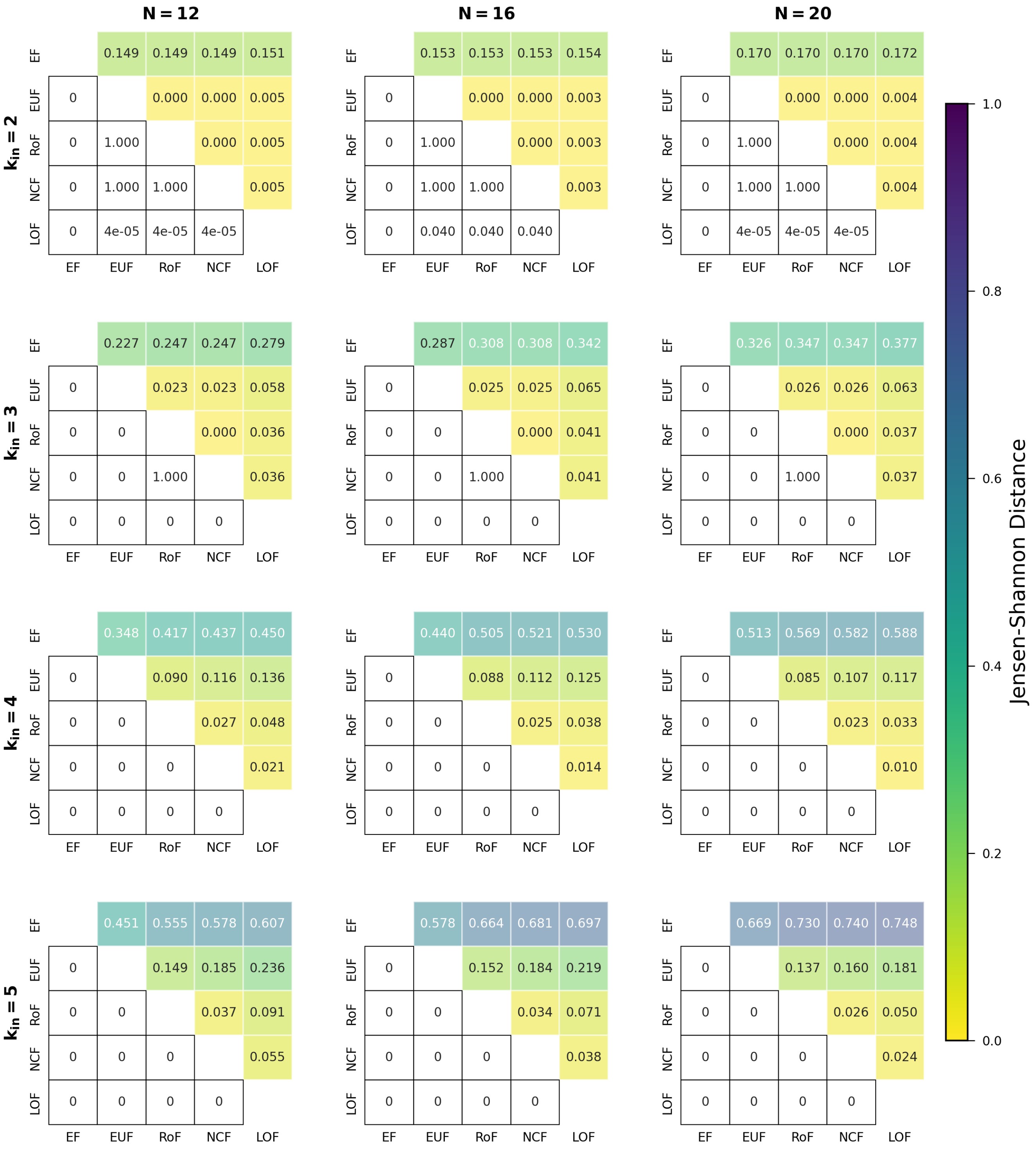


FIG. S16. **Comparing distributions for the average attractor lengths when using different classes of BFs within the R-P networks.** The different sub-figures are organised according to the values of $k_{in}$ (the in-degree) and $N$ (the number of nodes in the network). At given $k_{in}$ and $N$, we determined the distribution of the average attractor lengths for each of the 5 classes of BFs. In each sub-figure, the upper triangular part displays the pairwise Jensen-Shannon distances of those distributions. The lower triangular part represents the p-values of the one-sided K-S test (cumulative distribution of the label of column less than that of the row). Entries shown as 0 are actually very small, with values less than $10^{-307}$.

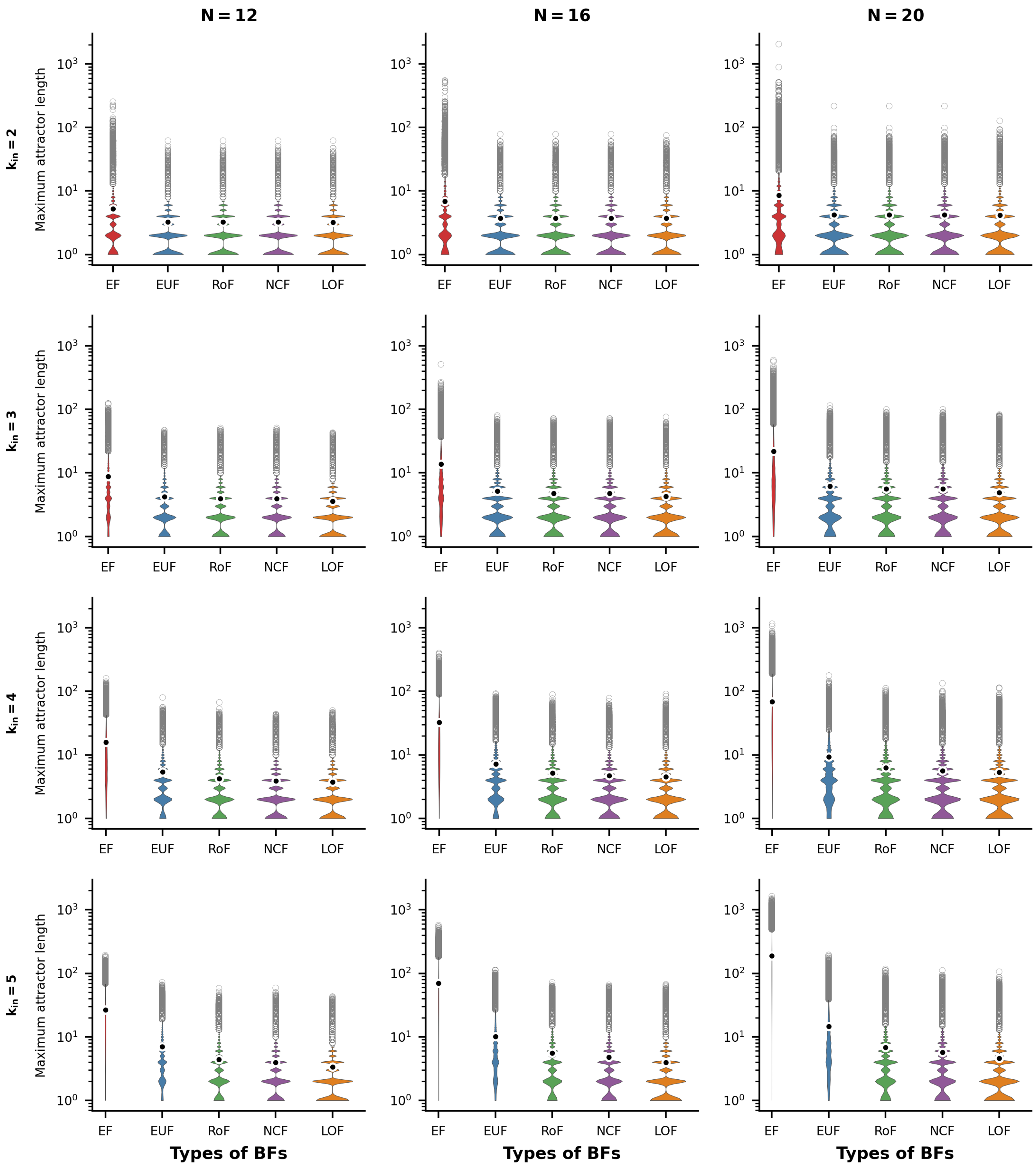


FIG. S17. **Distributions of maximum attractor lengths for R-P networks.** The violin plots display the distribution of the maximum attractor length for the R-P network topology. The sub-figures are organized into 3 columns, corresponding to the network sizes $N = 12, 16$ and $20$ respectively. Each row represents a different number of inputs per node, ranging from 2 to 5. Within each sub-figure, there are five violins representing the distributions of the maximum attractor length for the different ensembles of models, namely EF, EUF, RoF, NCF and LOF. Each violin distribution is based on $10^6$ data points. Outliers of each distribution are shown as gray rings while the means are shown as black dots encircled with white rings. Note the logarithmic scale for the y-axis.

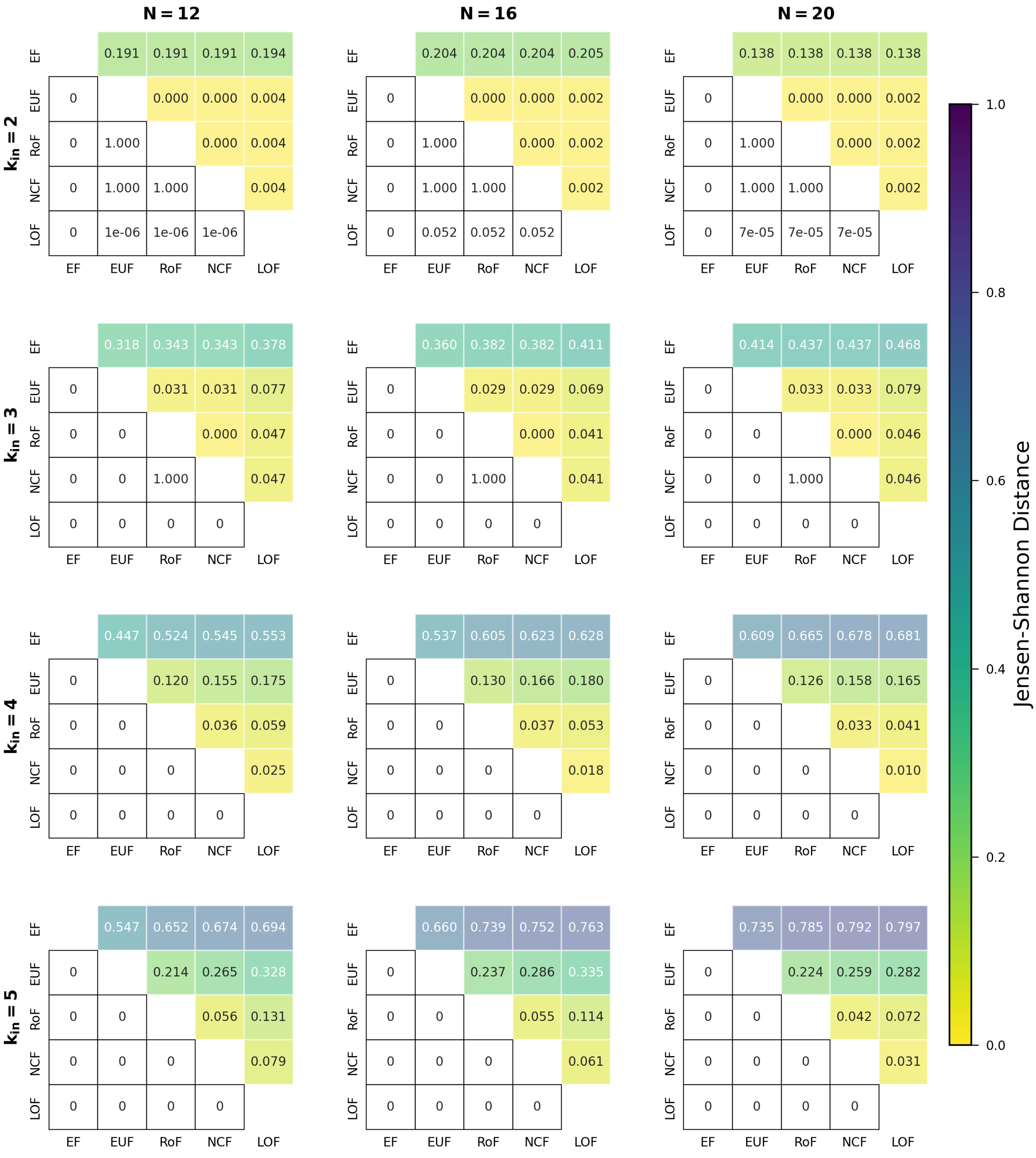

FIG. S18. **Comparing distributions for the maximum attractor lengths when using different classes of BFs within the R-P networks.** The different sub-figures are organised according to the values of $k_{in}$ (the in-degree) and $N$ (the number of nodes in the network). At given $k_{in}$ and $N$, we determined the distribution of the maximum attractor lengths for each of the 5 classes of BFs. In each sub-figure, the upper triangular part displays the pairwise Jensen-Shannon distances of those distributions. The lower triangular part represents the p-values of the one-sided K-S test (cumulative distribution of the label of column less than that of the row). Entries shown as 0 are actually very small, with values less than $10^{-307}$.

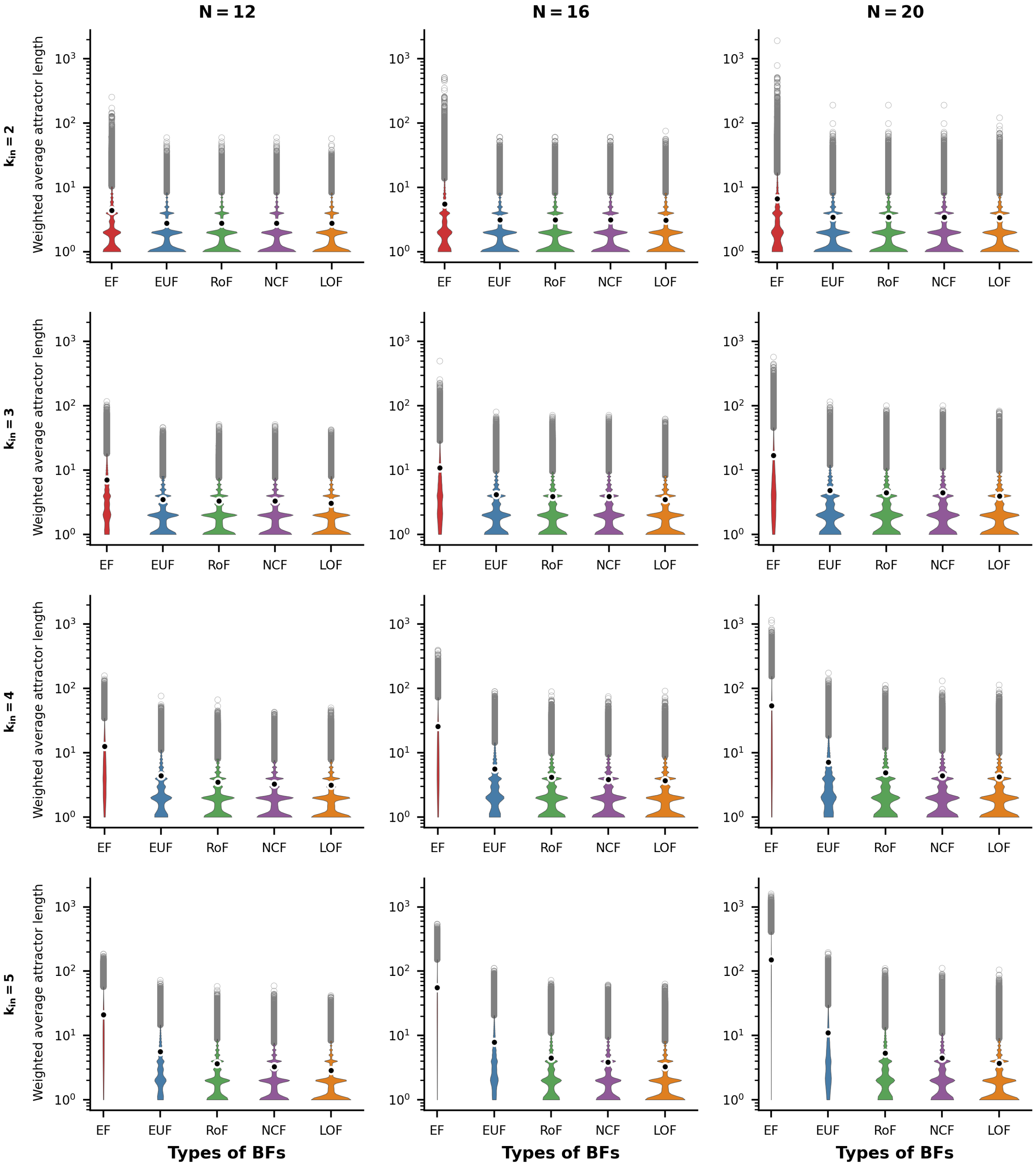


FIG. S19. **Distributions of weighted average attractor lengths for R-P networks.** The violin plots display the distribution of the weighted average attractor length for the R-P network topology. The sub-figures are organized into 3 columns, corresponding to the network sizes $N = 12, 16$ and 20 respectively. Each row represents a different number of inputs per node, ranging from 2 to 5. Within each sub-figure, there are five violins representing the distributions of the weighted average attractor length for the different ensembles of models, namely EF, EUF, RoF, NCF and LOF. Each violin distribution is based on $10^6$ data points. Outliers of each distribution are shown as gray rings while the means are shown as black dots encircled with white rings. Note the logarithmic scale for the y-axis.

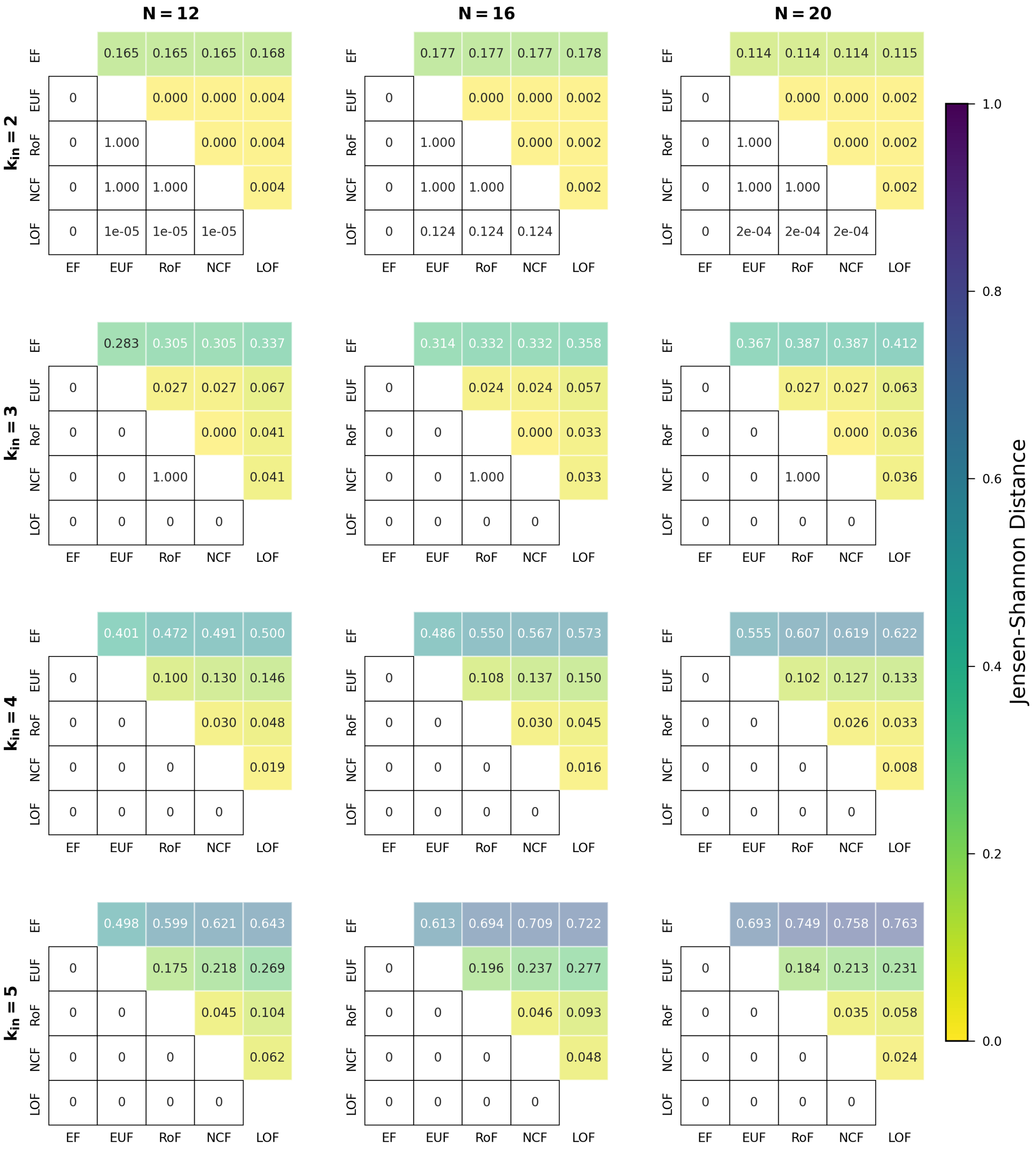


FIG. S20. **Comparing distributions for the weighted average attractor lengths when using different classes of BFs within the R-P networks.** The different sub-figures are organised according to the values of $k_{in}$ (the in-degree) and $N$ (the number of nodes in the network). At given $k_{in}$ and $N$, we determined the distribution of the weighted average attractor lengths for each of the 5 classes of BFs. In each sub-figure, the upper triangular part displays the pairwise Jensen-Shannon distances of those distributions. The lower triangular part represents the p-values of the one-sided K-S test (cumulative distribution of the label of column less than that of the row). Entries shown as 0 are actually very small, with values less than $10^{-307}$.

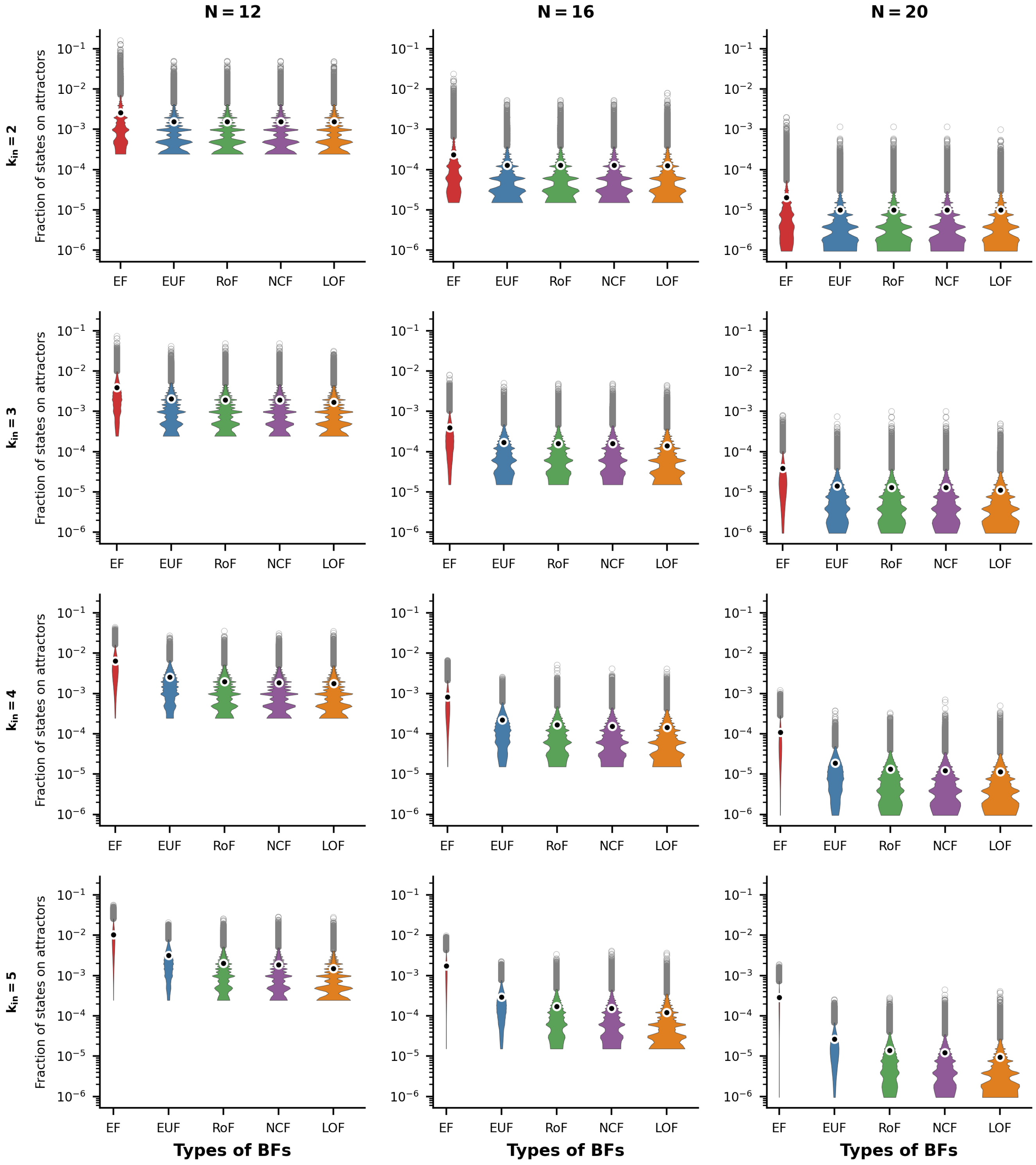


FIG. S21. **Distributions of fraction of states on attractors for R-P networks.** The violin plots display the distribution of the fraction of states on attractors for the R-P network topology. The sub-figures are organized into 3 columns, corresponding to the network sizes $N = 12, 16$ and 20 respectively. Each row represents a different number of inputs per node, ranging from 2 to 5. Within each sub-figure, there are five violins representing the distributions of the fraction of states on attractors for the different ensembles of models, namely EF, EUF, RoF, NCF and LOF. Each violin distribution is based on $10^6$ data points. Outliers of each distribution are shown as gray rings while the means are shown as black dots encircled with white rings. Note the logarithmic scale for the y-axis.

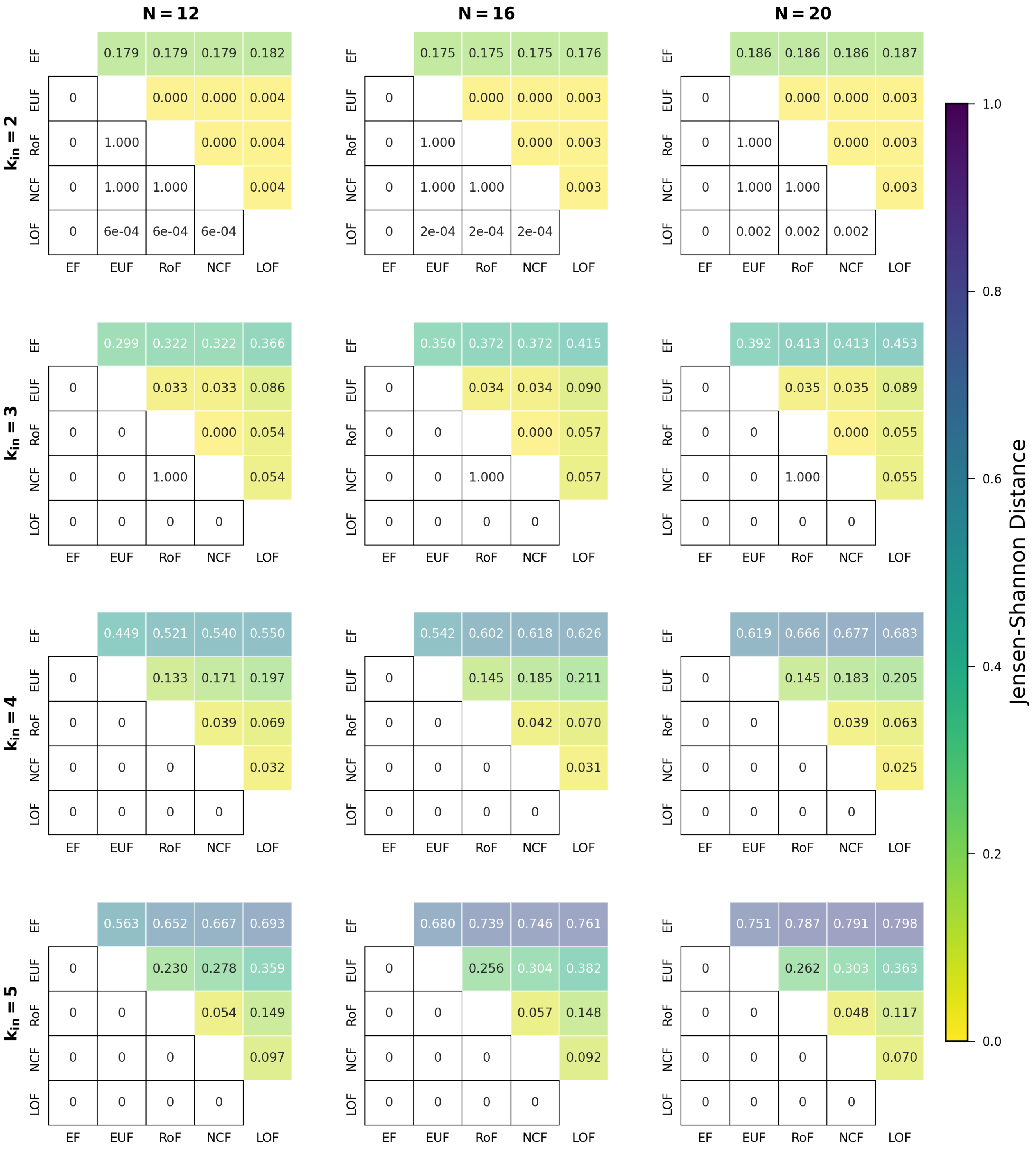

FIG. S22. **Comparing distributions for the fraction of states on attractors when using different classes of BFs within the R-P networks.** The different sub-figures are organised according to the values of $k_{in}$ (the in-degree) and $N$ (the number of nodes in the network). At given $k_{in}$ and $N$, we determined the distribution of the fraction of states on attractors for each of the 5 classes of BFs. In each sub-figure, the upper triangular part displays the pairwise Jensen-Shannon distances of those distributions. The lower triangular part represents the p-values of the one-sided K-S test (cumulative distribution of the label of column less than that of the row). Entries shown as 0 are actually very small, with values less than $10^{-307}$.

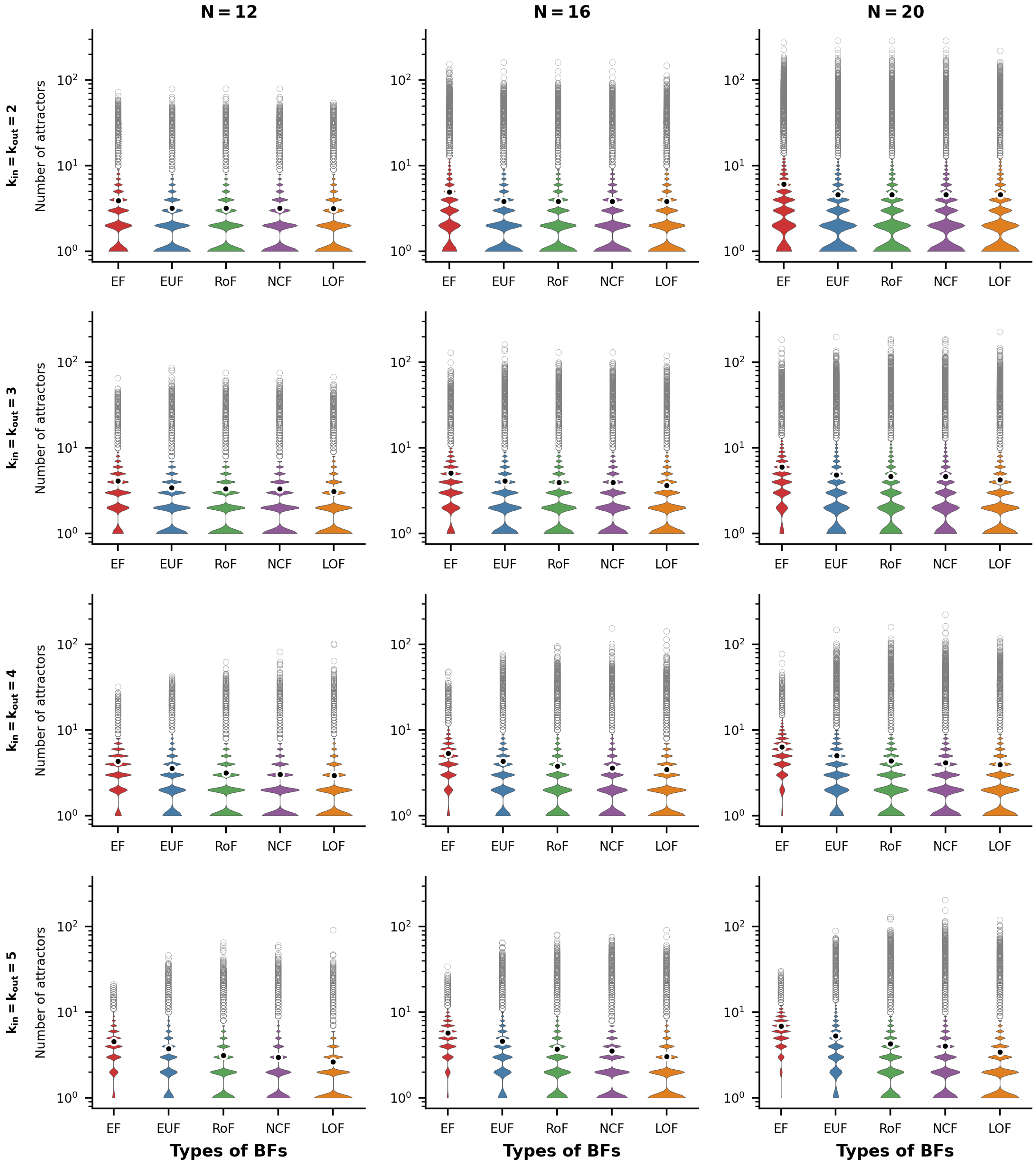


FIG. S23. **Distributions of number of attractors for R-R networks.** The violin plots display the distribution of the number of attractors for the R-R network topology. The sub-figures are organized into 3 columns, corresponding to the network sizes $N = 12, 16$ and 20 respectively. Each row represents a different number of inputs (and outputs) per node, ranging from 2 to 5. Within each sub-figure, there are five violins representing the distributions of the number of attractors for the different ensembles of models, namely EF, EUF, RoF, NCF and LOF. Each violin distribution is based on $10^6$ data points. Outliers of each distribution are shown as gray rings while the means are shown as black dots encircled with white rings. Note the logarithmic scale for the y-axis.

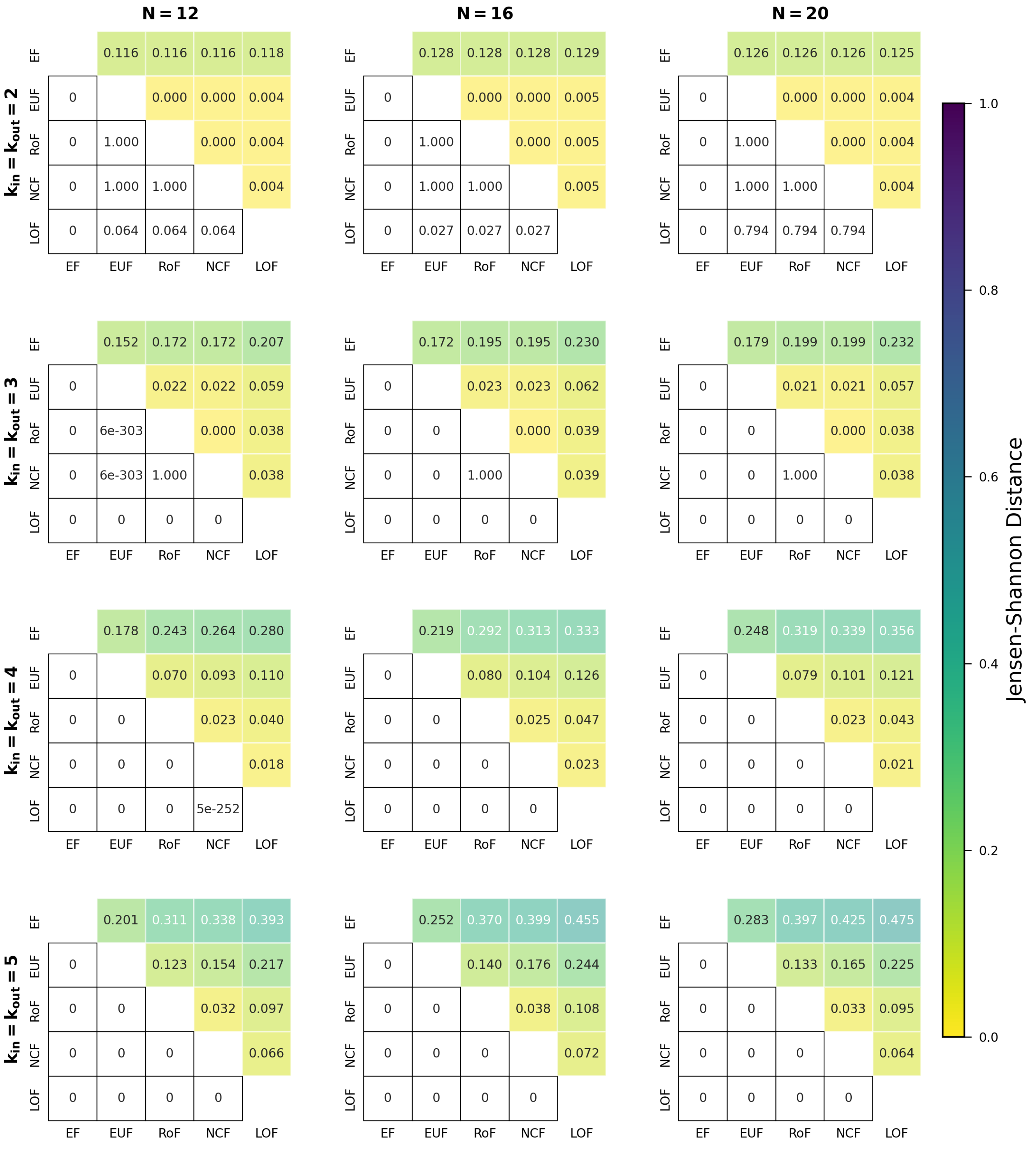


FIG. S24. **Comparing distributions for the number of attractors when using different classes of BFs within the R-R networks.** The different sub-figures are organised according to the values of the in-degree and out-degree ($k_{in}$ and $k_{out}$ both equal), and $N$ (the number of nodes in the network). At given $k_{in}$ (or $k_{out}$) and $N$, we determined the distribution of the number of attractors for each of the 5 classes of BFs. In each sub-figure, the upper triangular part displays the pairwise Jensen-Shannon distances of those distributions. The lower triangular part represents the p-values of the one-sided K-S test (cumulative distribution of the label of column less than that of the row). Entries shown as 0 are actually very small, with values less than $10^{-307}$.

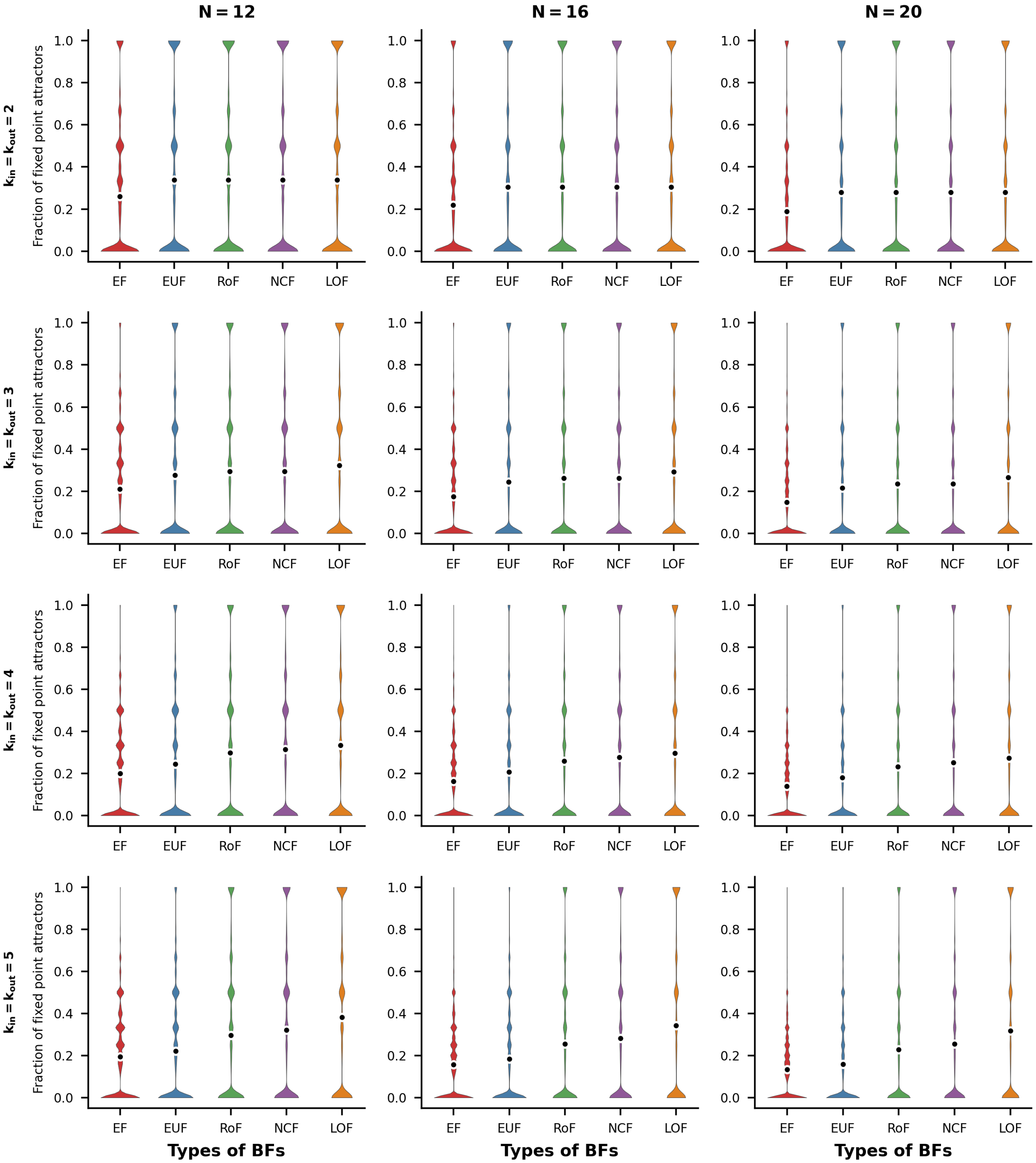


FIG. S25. **Distributions of fraction of fixed point attractors for R-R networks.** The violin plots display the distribution of the fraction of fixed point attractors for the R-R network topology. The sub-figures are organized into 3 columns, corresponding to the network sizes $N = 12, 16$ and 20 respectively. Each row represents a different number of inputs (and outputs) per node, ranging from 2 to 5. Within each sub-figure, there are five violins representing the distributions of the fraction of fixed point attractors for the different ensembles of models, namely EF, EUF, RoF, NCF and LOF. Each violin distribution is based on $10^6$ data points. Outliers of each distribution are shown as gray rings while the means are shown as black dots encircled with white rings.

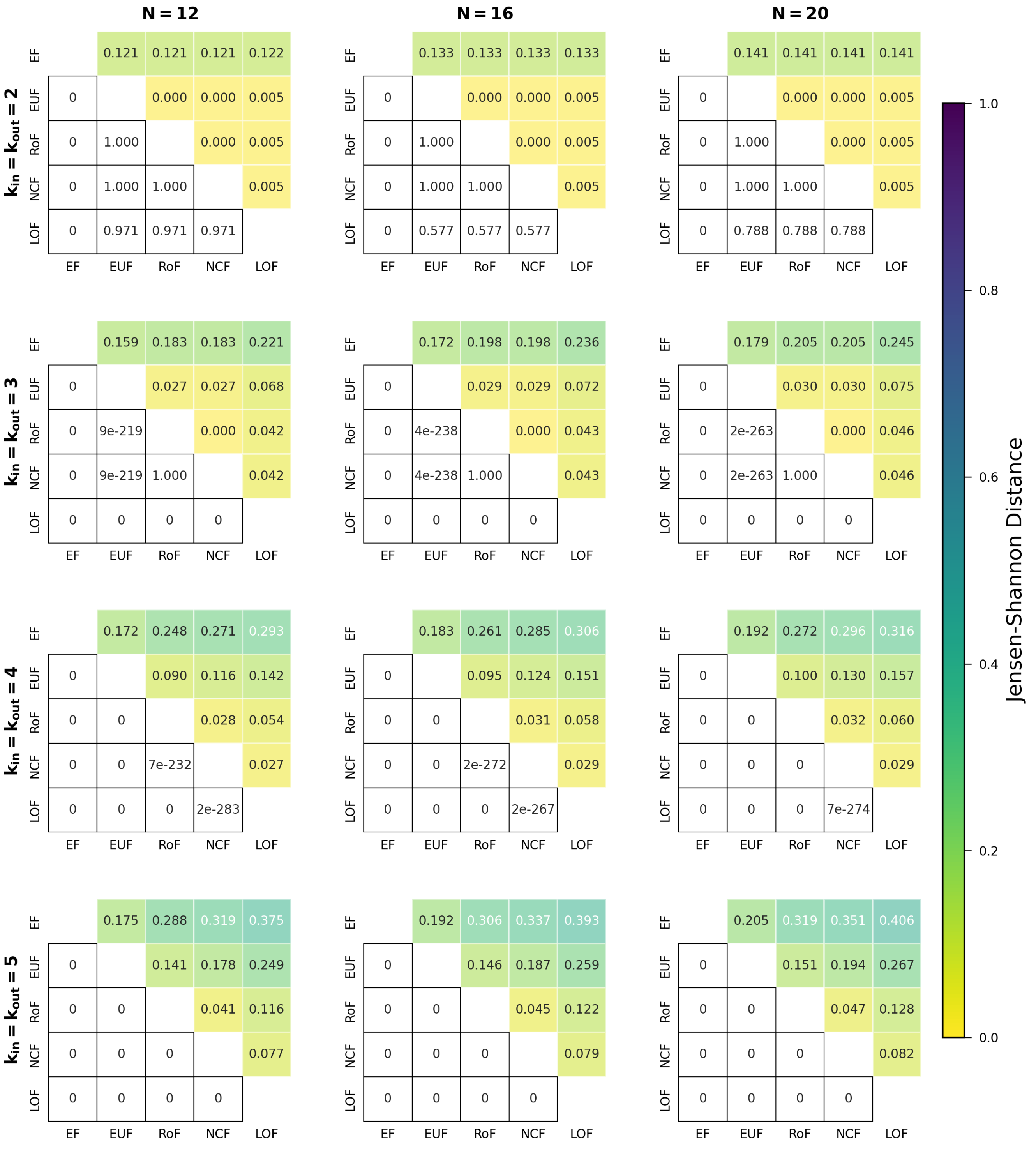


FIG. S26. **Comparing distributions for the fraction of fixed point attractors when using different classes of BFs within the R-R networks.** The different sub-figures are organised according to the values of the in-degree and out-degree ($k_{in}$ and $k_{out}$ both equal), and $N$ (the number of nodes in the network). At given $k_{in}$ (or $k_{out}$) and $N$, we determined the distribution of the fraction of fixed point attractors for each of the 5 classes of BFs. In each sub-figure, the upper triangular part displays the pairwise Jensen-Shannon distances of those distributions. The lower triangular part represents the p-values of the one-sided K-S test (cumulative distribution of the label of column greater than that of the row). Entries shown as 0 are actually very small, with values less than $10^{-307}$.

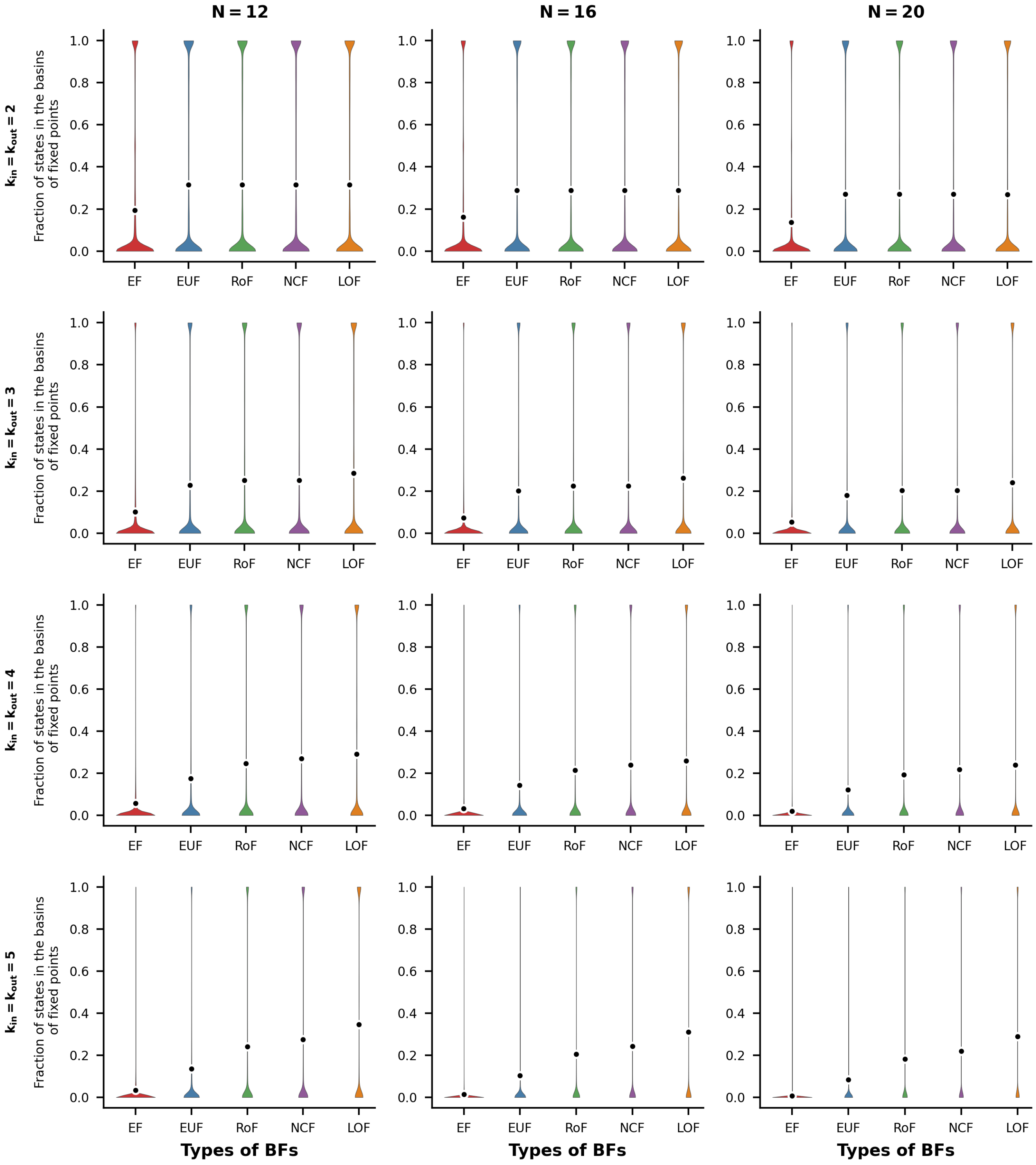


FIG. S27. **Distributions of Fraction of states in the basins of fixed points for R-R networks** The violin plots display the distribution of the fraction of states in the basins of fixed point attractors for the R-R network topology. The sub-figures are organized into 3 columns, corresponding to the network sizes $N = 12, 16$ and 20 respectively. Each row represents a different number of inputs (and outputs) per node, ranging from 2 to 5. Within each sub-figure, there are five violins representing the distributions of the fraction of states in the basins of fixed point attractors for the different ensembles of models, namely EF, EUF, RoF, NCF and LOF. Each violin distribution is based on $10^6$ data points. Outliers of each distribution are shown as gray rings while the means are shown as black dots encircled with white rings.

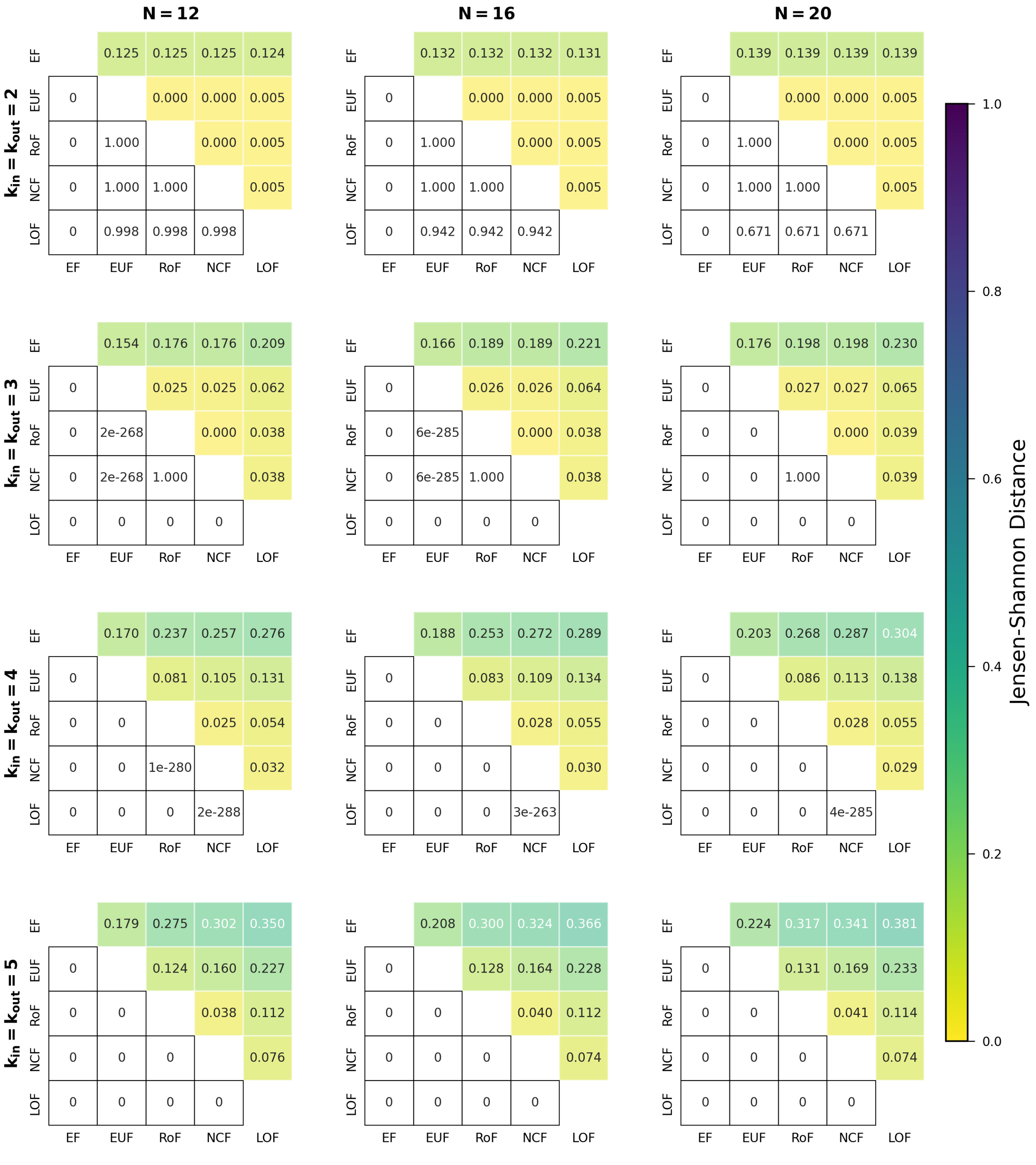

FIG. S28. **Comparing distributions for the fraction of states in the basins of fixed points when using different classes of BFs within the R-R networks.** The different sub-figures are organised according to the values of the in-degree and out-degree,($k_{in}$ and $k_{out}$ both equal), and $N$ (the number of nodes in the network). At given $k_{in}$ (or $k_{out}$) and $N$, we determined the distribution of the Fraction of states in the basins of fixed points for each of the 5 classes of BFs. In each sub-figure, the upper triangular part displays the pairwise Jensen-Shannon distances of those distributions. The lower triangular part represents the p-values of the one-sided K-S test (cumulative distribution of the label of column greater than that of the row). Entries shown as 0 are actually very small, with values less than $10^{-307}$.

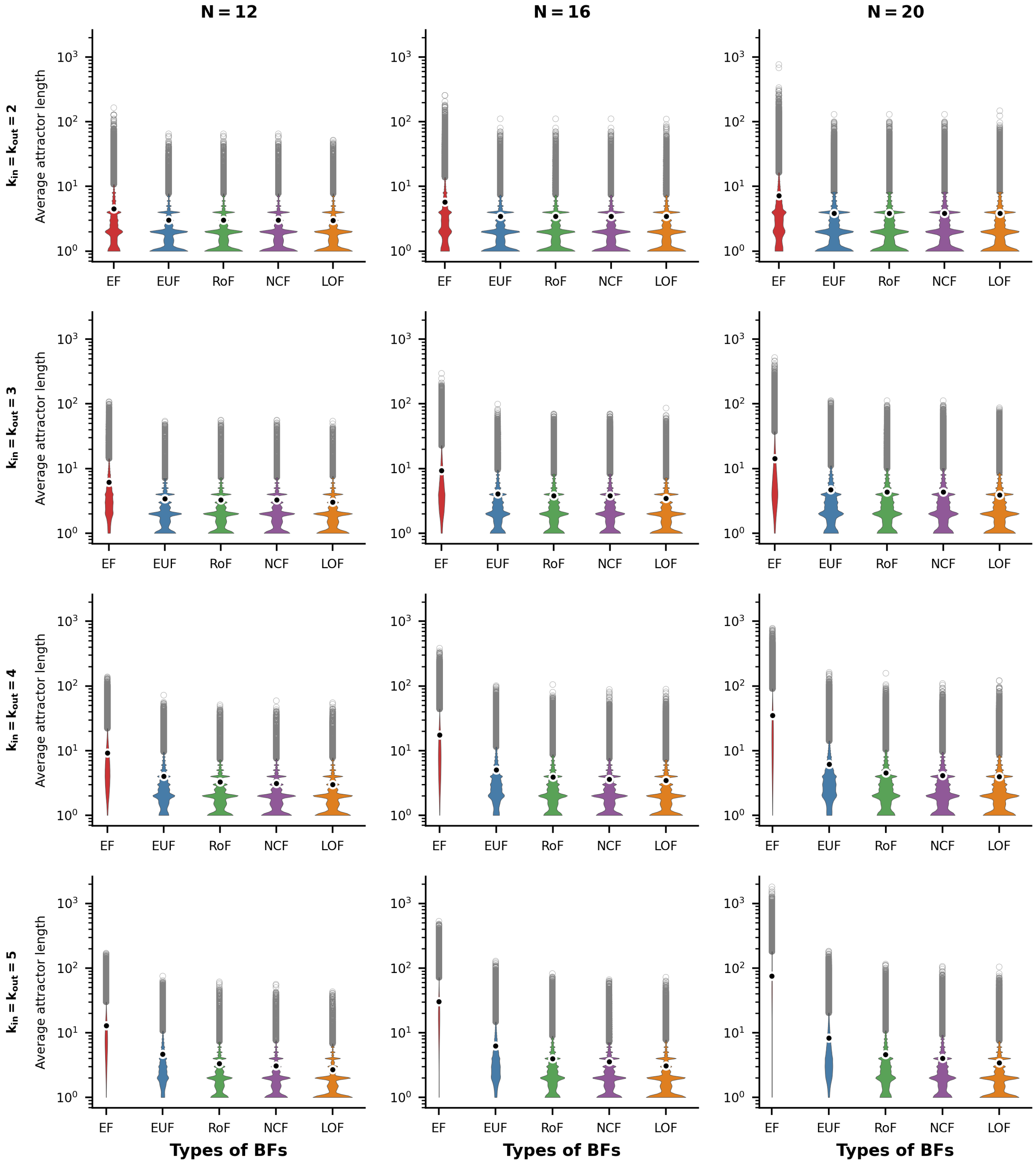


FIG. S29. **Distributions of average attractor lengths for R-R networks.** The violin plots display the distribution of the average attractor length for the R-R network topology. The sub-figures are organized into 3 columns, corresponding to the network sizes $N = 12, 16$ and 20 respectively. Each row represents a different number of inputs (and outputs) per node, ranging from 2 to 5. Within each sub-figure, there are five violins representing the distributions of the average attractor length for the different ensembles of models, namely EF, EUF, RoF, NCF and LOF. Each violin distribution is based on $10^6$ data points. Outliers of each distribution are shown as gray rings while the means are shown as black dots encircled with white rings. Note the logarithmic scale for the y-axis.

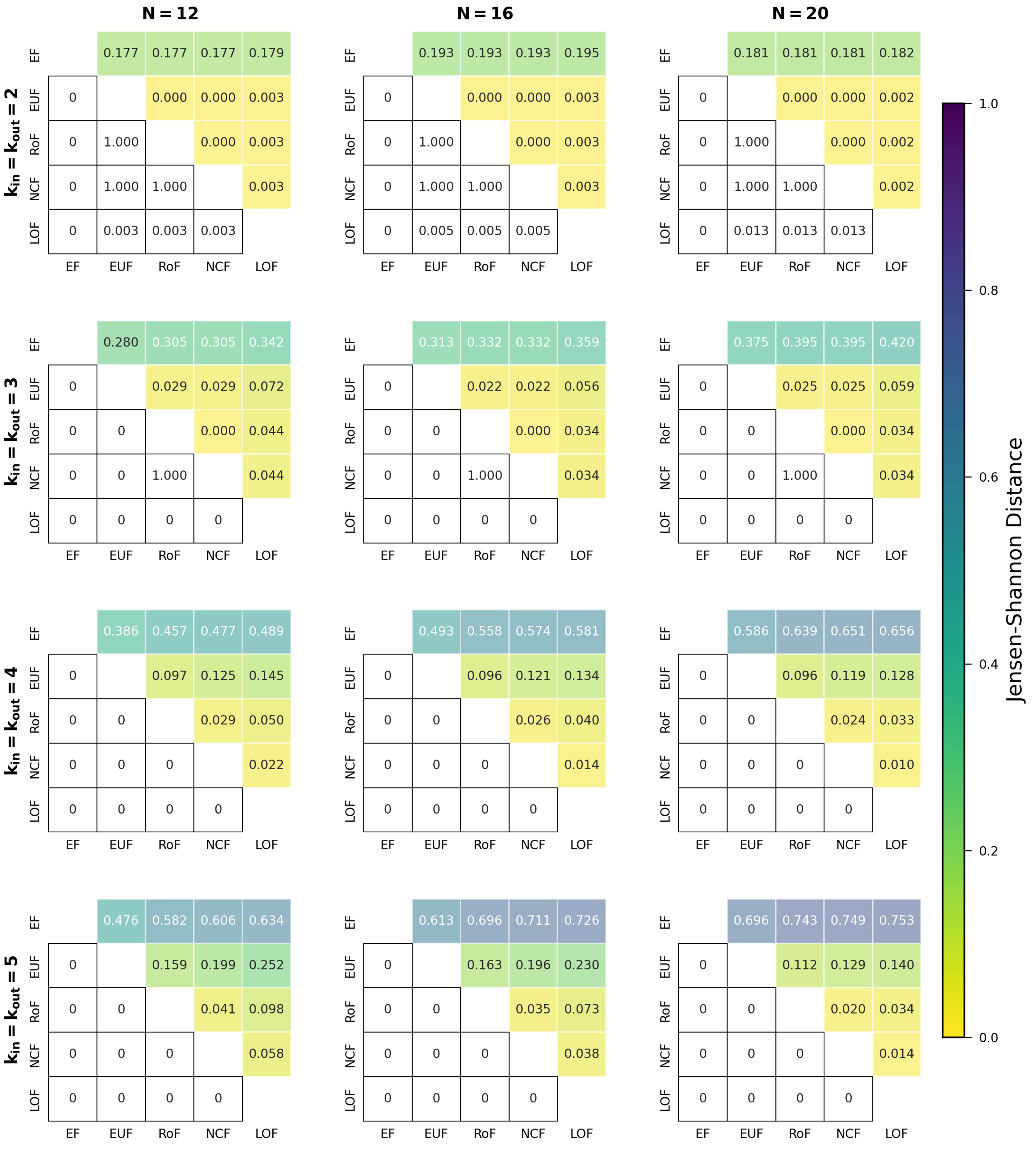


FIG. S30. **Comparing distributions for the average attractor lengths when using different classes of BFs within the R-R networks.** The different sub-figures are organised according to the values of the in-degree and out-degree ($k_{in}$ and $k_{out}$ both equal), and $N$ (the number of nodes in the network). At given $k_{in}$ (or $k_{out}$) and $N$, we determined the distribution of the average attractor lengths for each of the 5 classes of BFs. In each sub-figure, the upper triangular part displays the pairwise Jensen-Shannon distances of those distributions. The lower triangular part represents the p-values of the one-sided K-S test (cumulative distribution of the label of column less than that of the row). Entries shown as 0 are actually very small, with values less than $10^{-307}$.

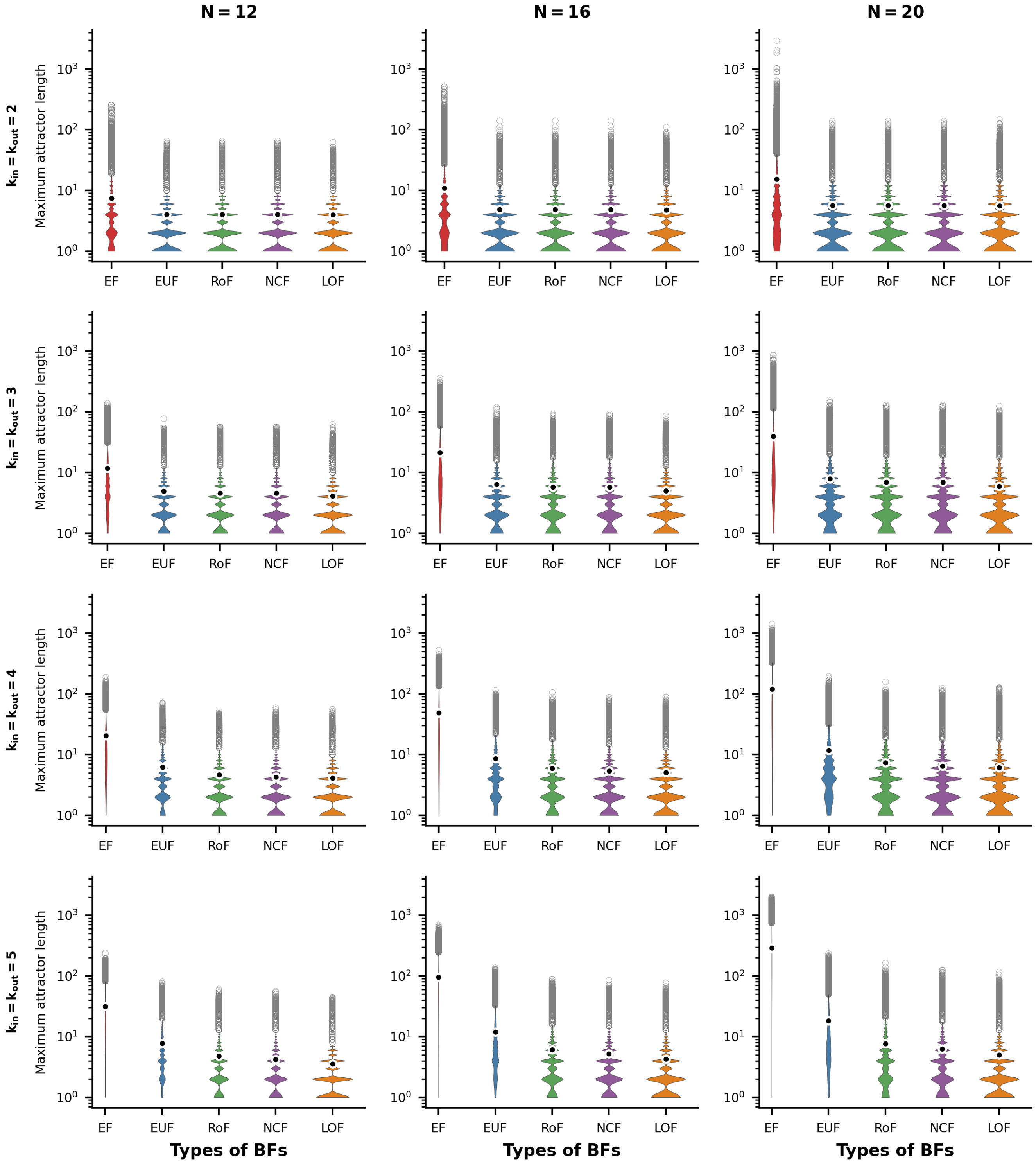


FIG. S31. **Distributions of Maximum attractor lengths for R-R networks.** The violin plots display the distribution of the maximum attractor length for the R-R network topology. The sub-figures are organized into 3 columns, corresponding to the network sizes $N = 12, 16$ and 20 respectively. Each row represents a different number of inputs (and outputs) per node, ranging from 2 to 5. Within each sub-figure, there are five violins representing the distributions of the maximum attractor length for the different ensembles of models, namely EF, EUF, RoF, NCF and LOF. Each violin distribution is based on $10^6$ data points. Outliers of each distribution are shown as gray rings while the means are shown as black dots encircled with white rings. Note the logarithmic scale for the y-axis.

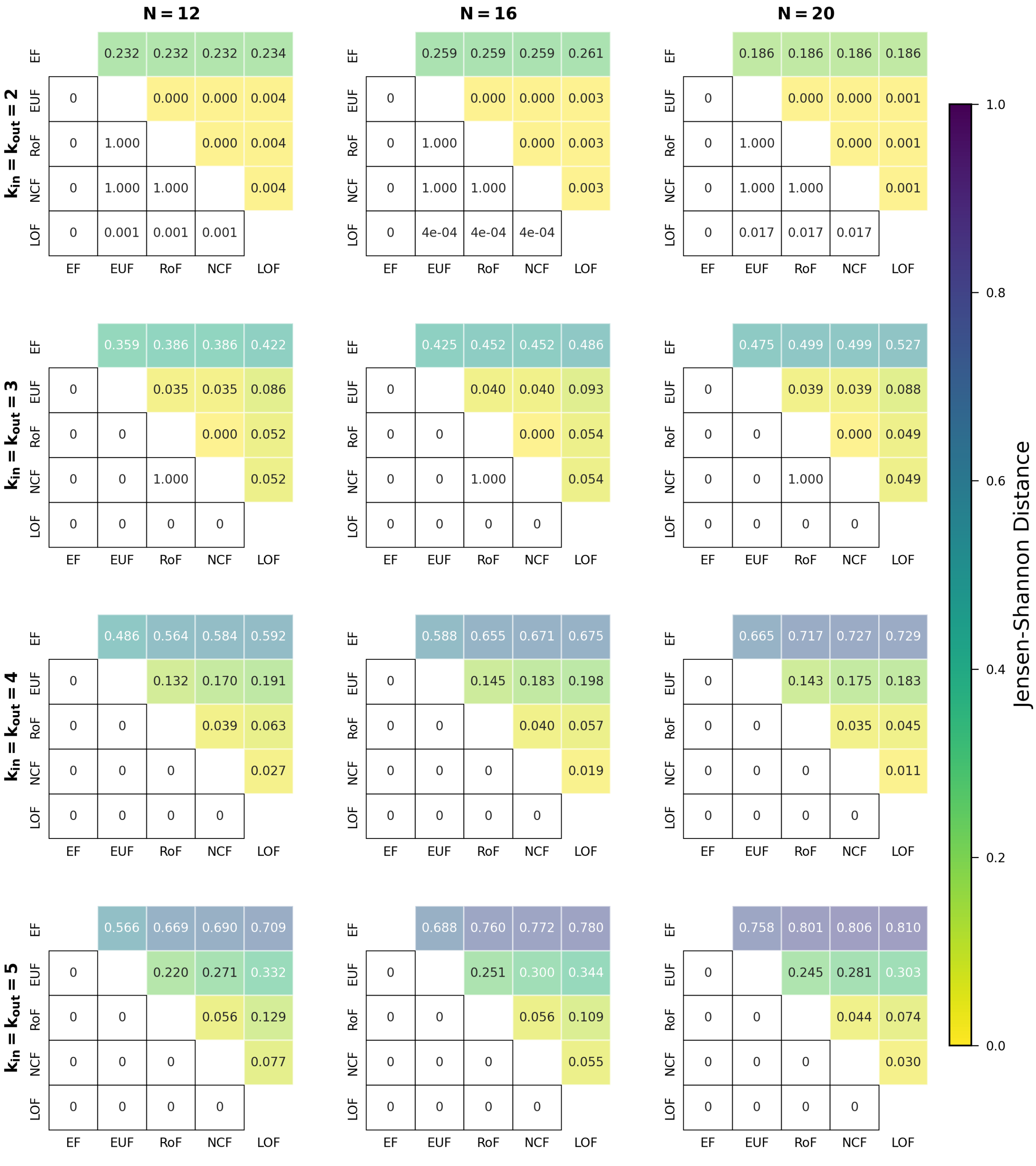


FIG. S32. **Comparing distributions for the maximum attractor lengths when using different classes of BFs within the R-R networks.** The different sub-figures are organised according to the values of the in-degree and out-degree ($k_{in}$ and $k_{out}$ both equal), and $N$ (the number of nodes in the network). At given $k_{in}$ (or $k_{out}$) and $N$, we determined the distribution of the maximum attractor lengths for each of the 5 classes of BFs. In each sub-figure, the upper triangular part displays the pairwise Jensen-Shannon distances of those distributions. The lower triangular part represents the p-values of the one-sided K-S test (cumulative distribution of the label of column less than that of the row). Entries shown as 0 are actually very small, with values less than $10^{-307}$.

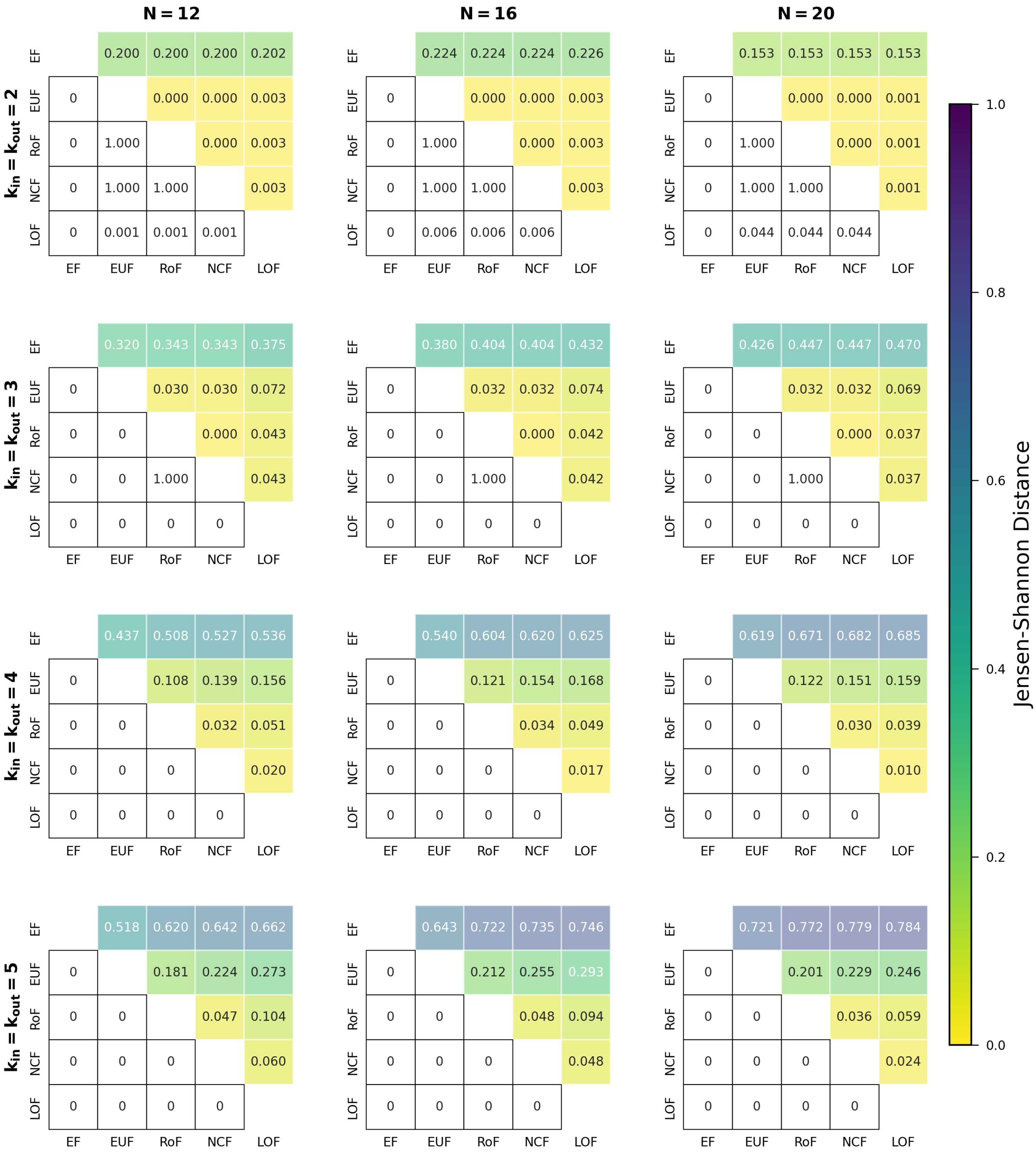


FIG. S33. **Comparing distributions for the weighted average attractor lengths when using different classes of BFs within the R-R networks.** The different sub-figures are organised according to the values of the in-degree and out-degree ($k_{in}$ and $k_{out}$ both equal), and $N$ (the number of nodes in the network). At given $k_{in}$ (or $k_{out}$) and $N$, we determined the distribution of the weighted average attractor lengths for each of the 5 classes of BFs. In each sub-figure, the upper triangular part displays the pairwise Jensen-Shannon distances of those distributions. The lower triangular part represents the p-values of the one-sided K-S test (cumulative distribution of the label of column less than that of the row). Entries shown as 0 are actually very small, with values less than $10^{-307}$.

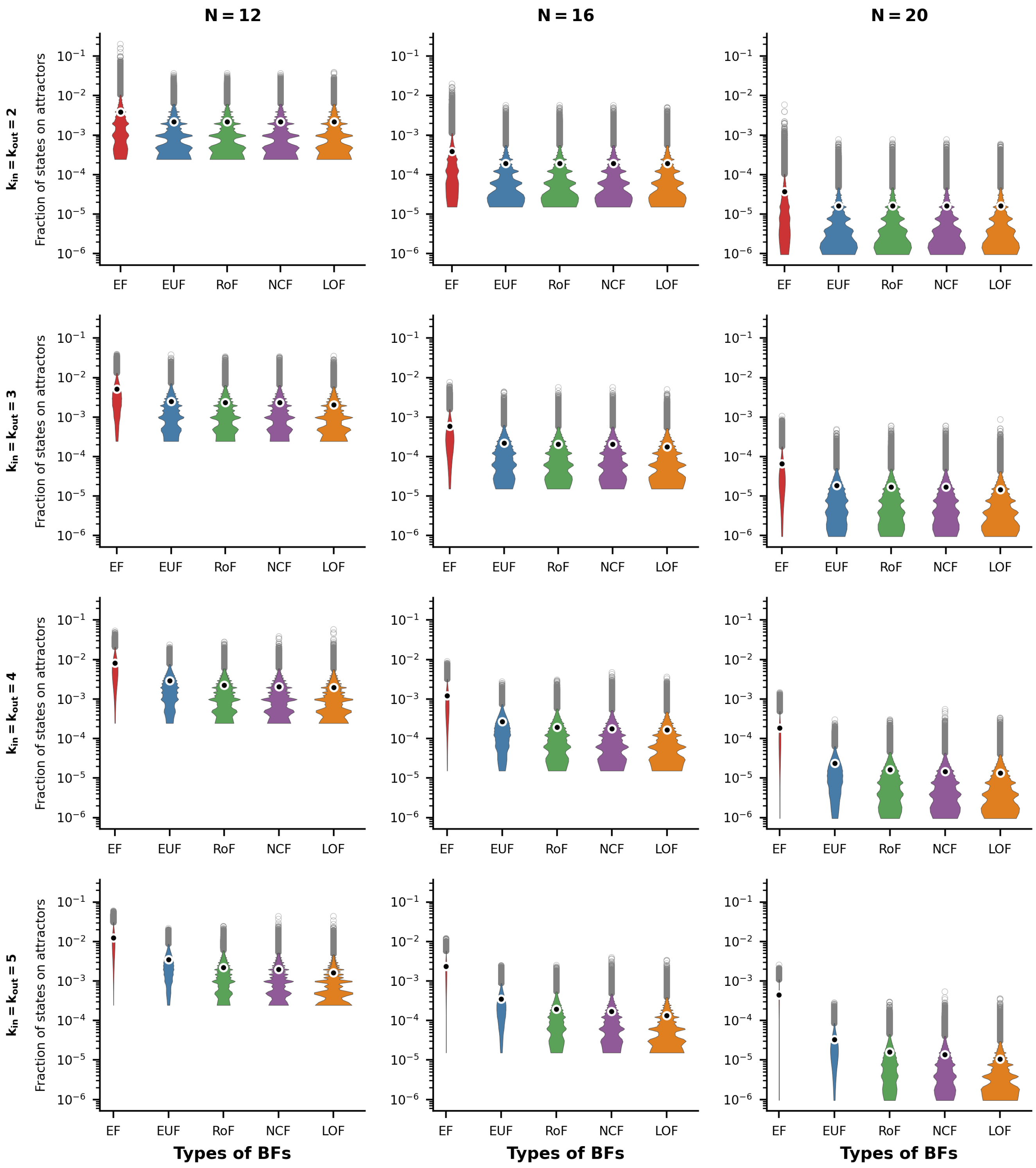


FIG. S34. **Distributions of fraction of states on attractors for R-R networks.** The violin plots display the distribution of the fraction of states on attractors for the R-R network topology. The sub-figures are organized into 3 columns, corresponding to the network sizes $N = 12, 16$ and 20 respectively. Each row represents a different number of inputs (and outputs) per node, ranging from 2 to 5. Within each sub-figure, there are five violins representing the distributions of the fraction of states on attractors for the different ensembles of models, namely EF, EUF, RoF, NCF and LOF. Each violin distribution is based on $10^6$ data points. Outliers of each distribution are shown as gray rings while the means are shown as black dots encircled with white rings. Note the logarithmic scale for the y-axis.

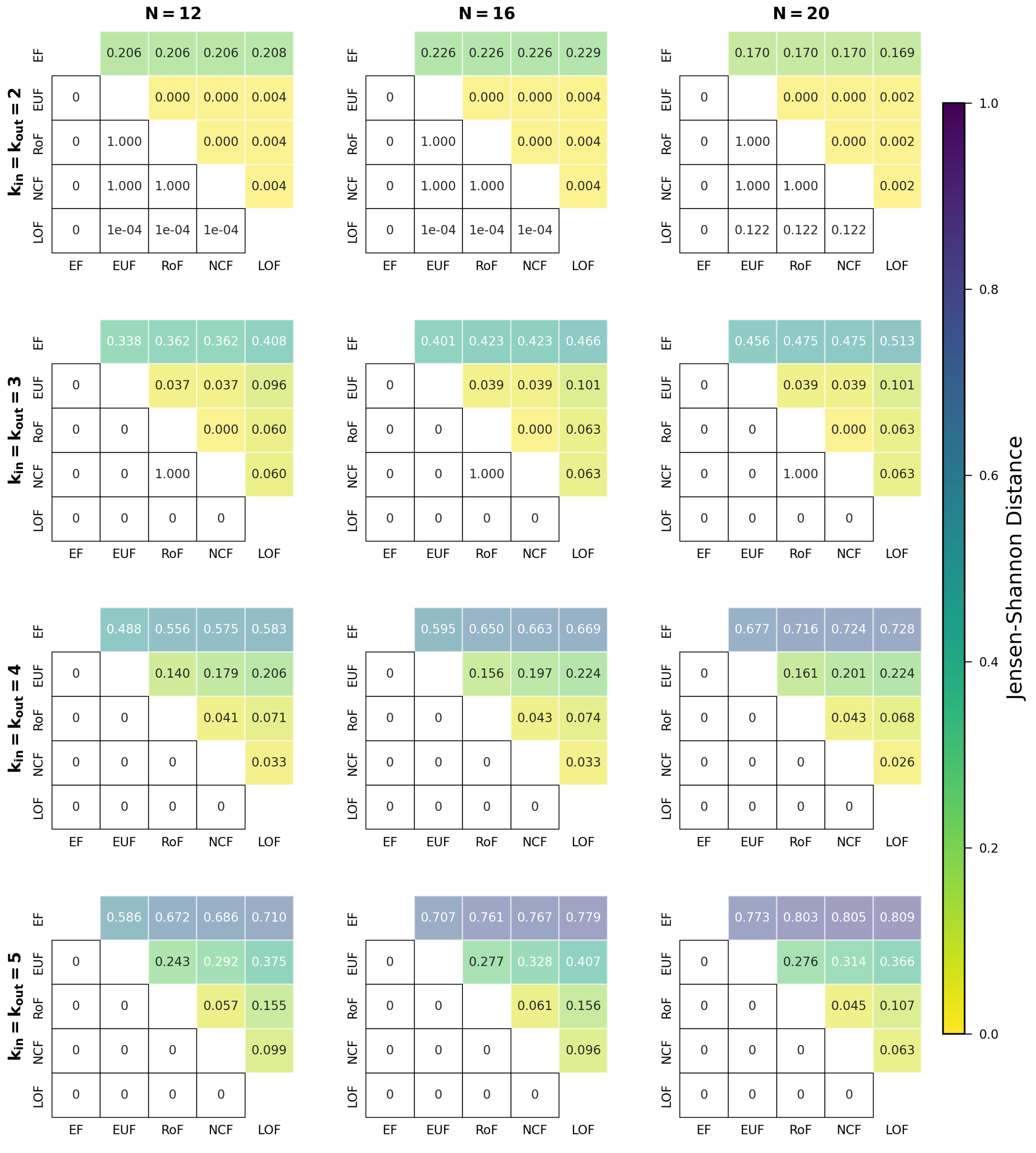

FIG. S35. **Comparing distributions for the fraction of states on attractors when using different classes of BFs within the R-R networks.** The different sub-figures are organised according to the values of the in-degree and out-degree ($k_{in}$ and $k_{out}$ both equal), and $N$ (the number of nodes in the network). At given $k_{in}$ (or $k_{out}$) and $N$, we determined the distribution of the fraction of states on attractors for each of the 5 classes of BFs. In each sub-figure, the upper triangular part displays the pairwise Jensen-Shannon distances of those distributions. The lower triangular part represents the p-values of the one-sided K-S test (cumulative distribution of the label of column less than that of the row). Entries shown as 0 are actually very small, with values less than $10^{-307}$.

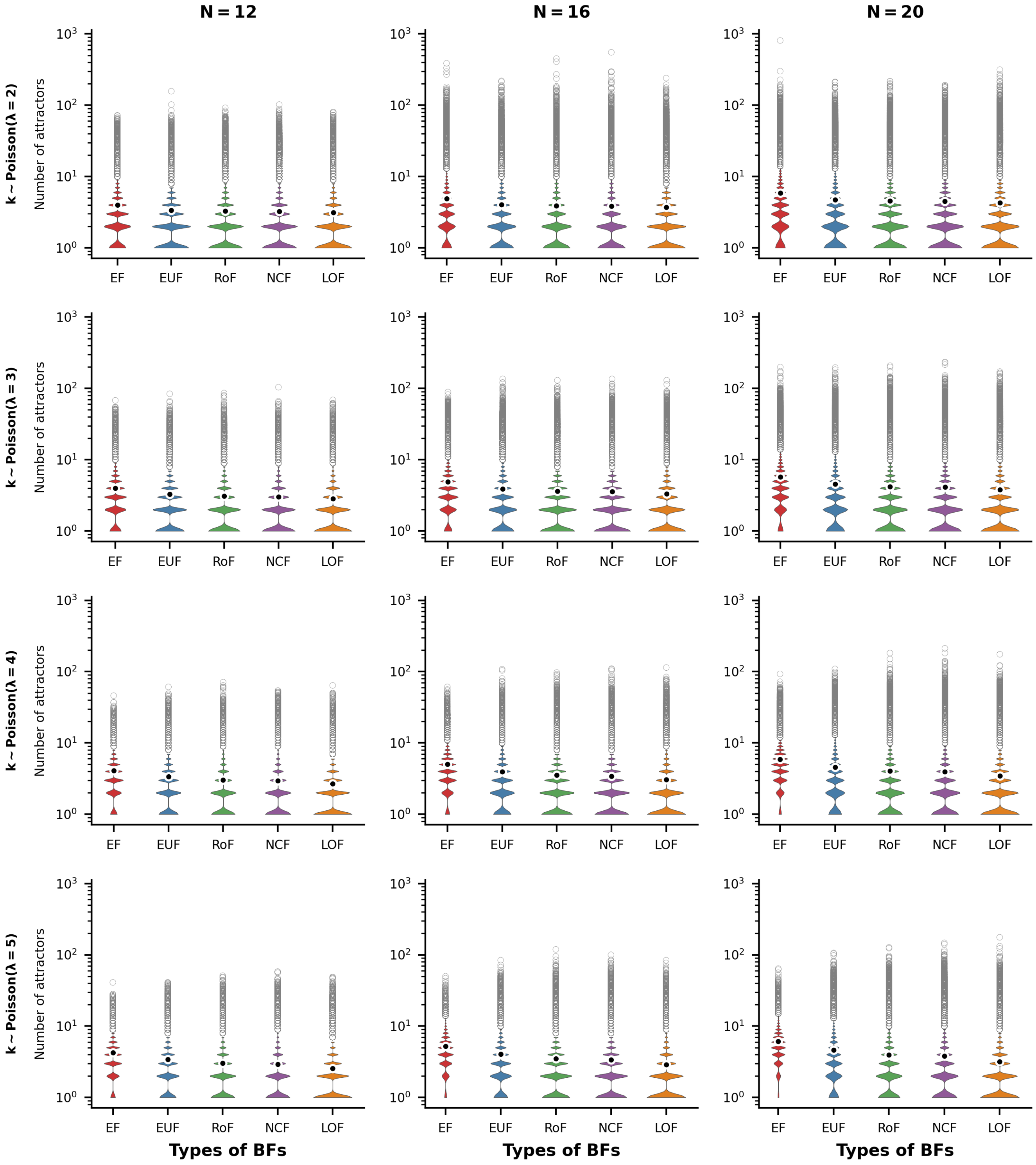


FIG. S36. **Distributions of number of attractors for P-P networks.** The violin plots display the distribution of the number of attractors for the P-P network topology. The sub-figures are organized into 3 columns, corresponding to the network sizes $N = 12, 16$ and 20 respectively. Each row represents a different average number of inputs per node, with the degrees drawn from Poisson distributions with means $2, 3, 4$ and 5. Within each sub-figure, there are five violins representing the distributions of the number of attractors for the different ensembles of models, namely EF, EUF, RoF, NCF and LOF. Each violin distribution is based on $10^6$ data points. Outliers of each distribution are shown as gray rings while the means are shown as black dots encircled with white rings. Note the logarithmic scale for the y-axis.

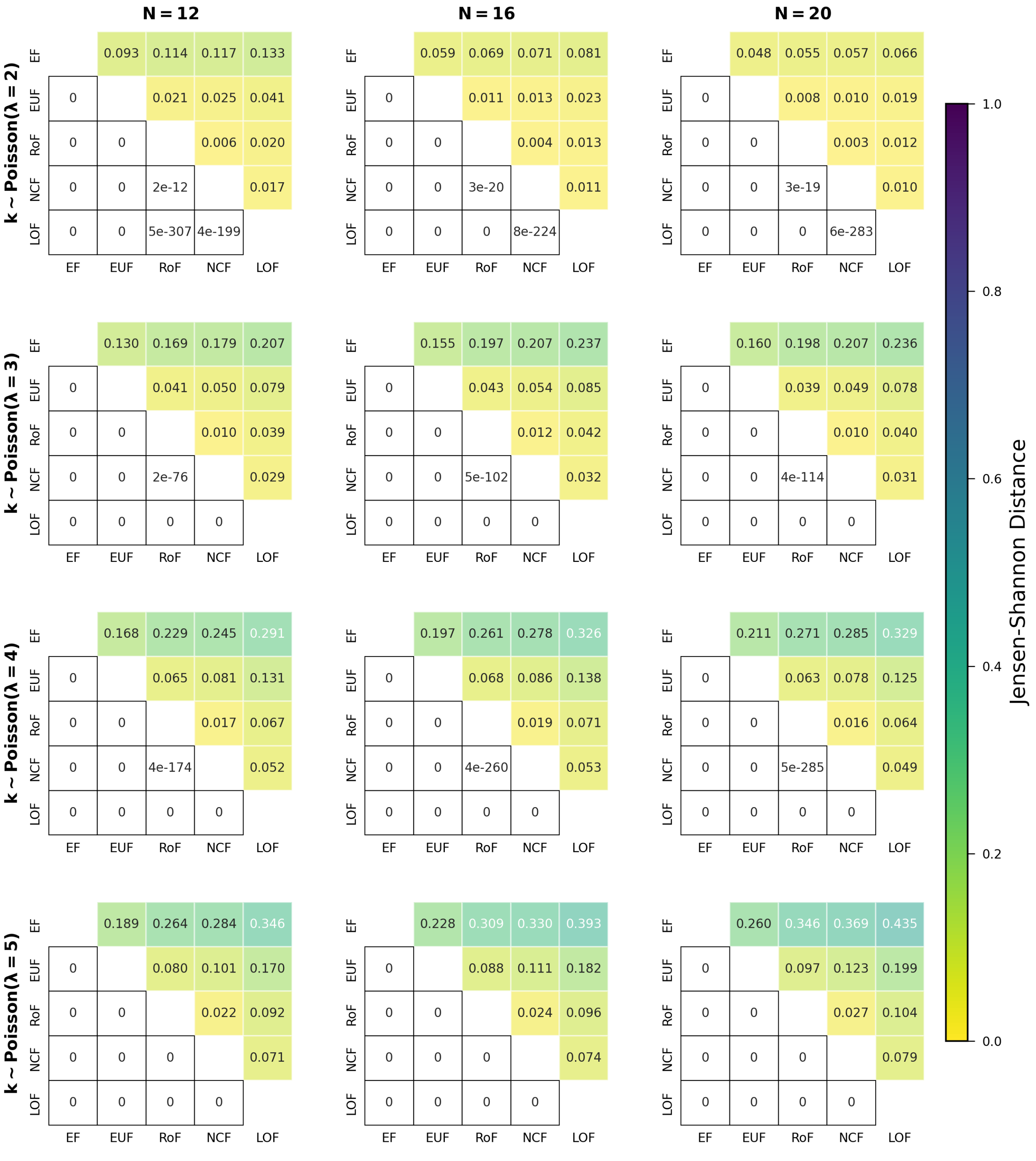


FIG. S37. **Comparing distributions for the number of attractors when using different classes of BFs within the P-P networks.** The different sub-figures are organised according to the values of $N$ (the number of nodes in the network) and $\lambda$ ($\lambda = 2, 3, 4$ and 5). The degrees of each node of the networks are drawn randomly from a Poisson distributions with mean $= \lambda$. At a given $\lambda$ and $N$, we determined the distribution of the number of attractors for each of the 5 classes of BFs. In each sub-figure, the upper triangular part displays the pairwise Jensen-Shannon distances of those distributions. The lower triangular part represents the p-values of the one-sided K-S test (cumulative distribution of the label of column less than that of the row). Entries shown as 0 are actually very small, with values less than $10^{-307}$.

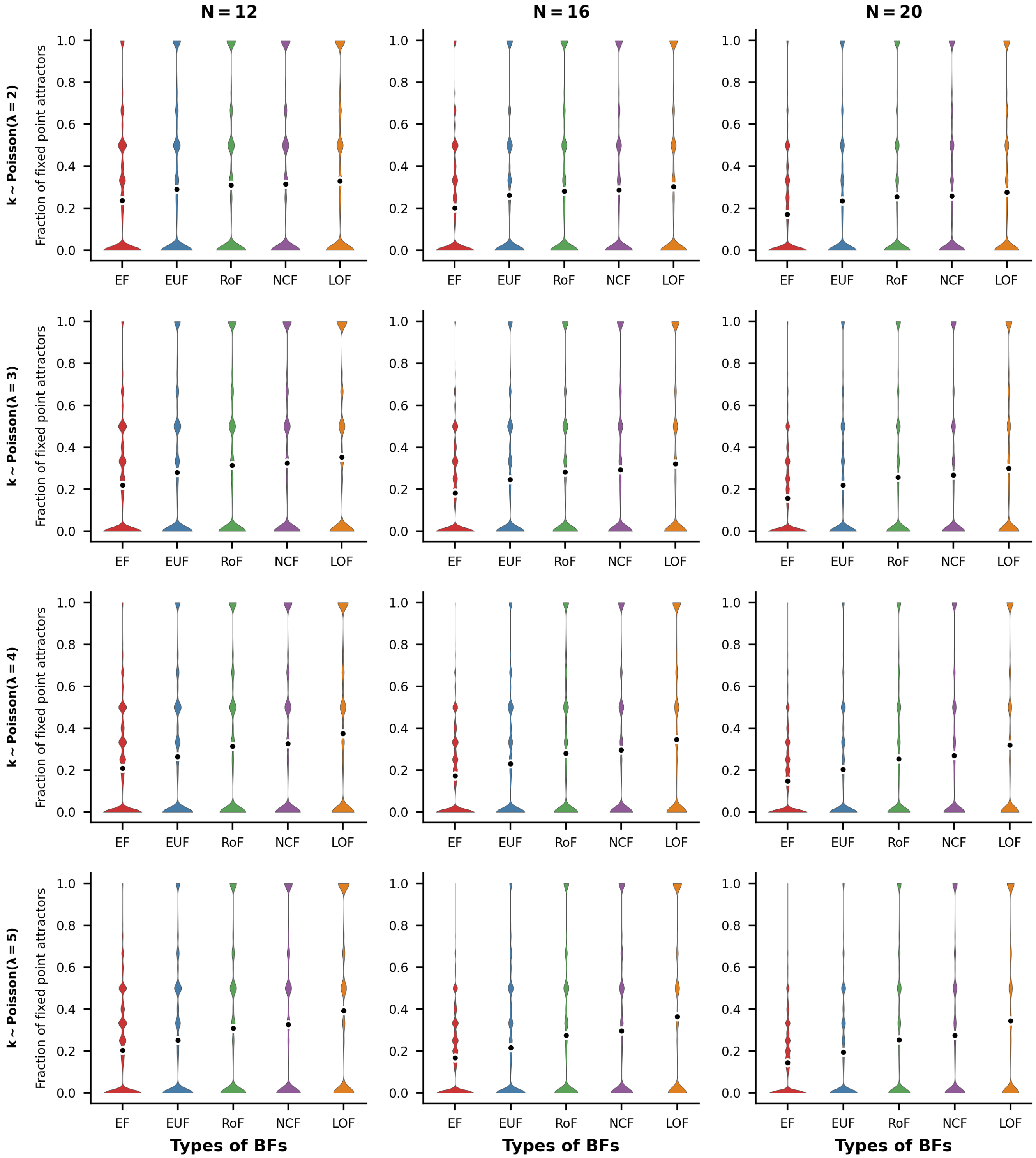


FIG. S38. **Distributions of fraction of fixed point attractors for P-P networks.** The violin plots display the distribution of the fraction of fixed point attractors for the P-P network topology. The sub-figures are organized into 3 columns, corresponding to the network sizes $N = 12, 16$ and $20$ respectively. Each row represents a different average number of inputs per node, with the degrees drawn from Poisson distributions with means $2, 3, 4$ and $5$. Within each sub-figure, there are five violins representing the distributions of the fraction of fixed point attractors for the different ensembles of models, namely EF, EUF, RoF, NCF and LOF. Each violin distribution is based on $10^6$ data points. Outliers of each distribution are shown as gray rings while the means are shown as black dots encircled with white rings.

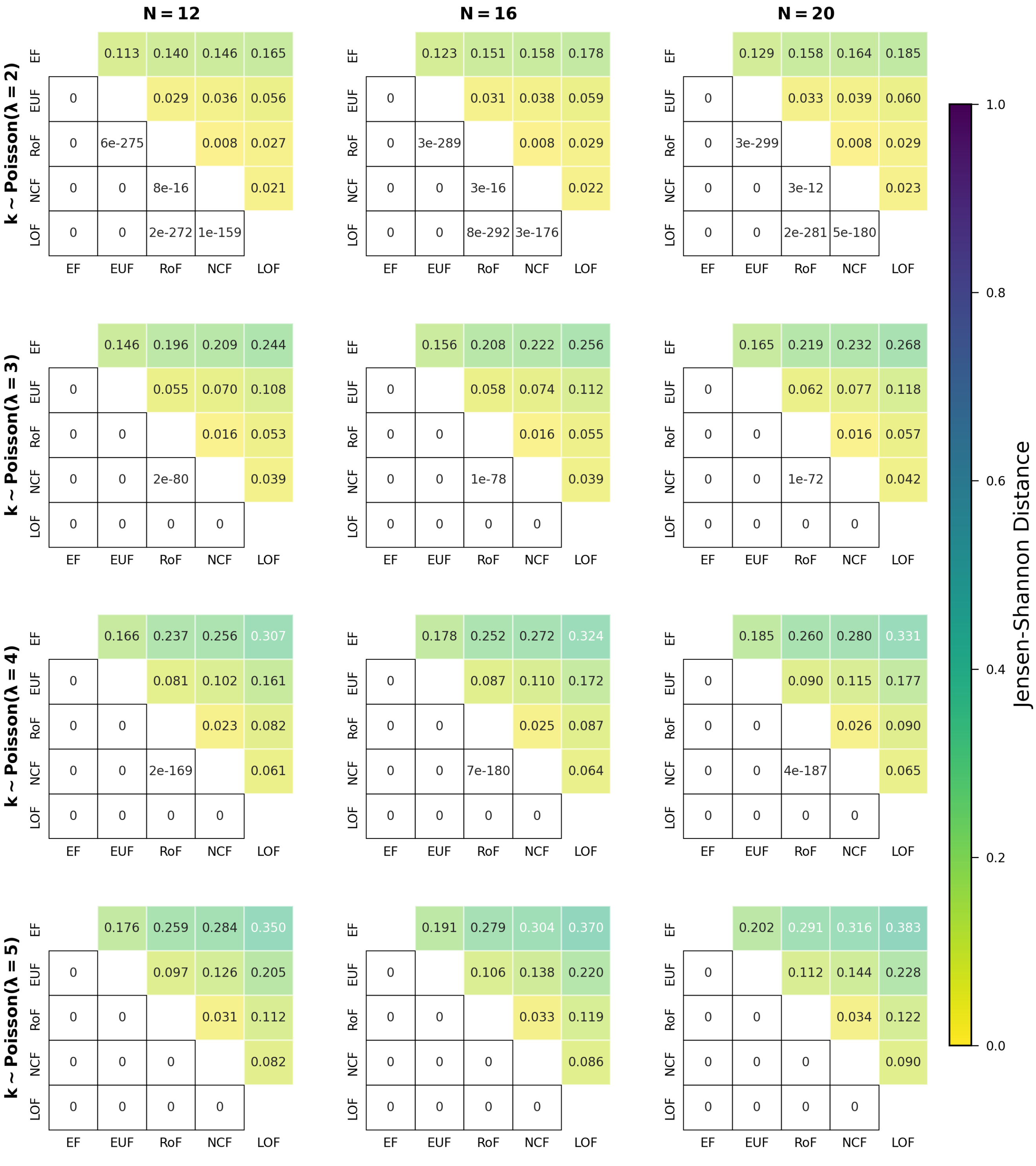


FIG. S39. **Comparing distributions for the fraction of fixed point attractors when using different classes of BFs within the P-P networks.** The different sub-figures are organised according to the values of $N$ (the number of nodes in the network) and $\lambda$ ($\lambda = 2, 3, 4$ and 5). The degrees of each node of the networks are drawn randomly from a Poisson distributions with mean $= \lambda$. At a given $\lambda$ and $N$, we determined the distribution of the fraction of fixed point attractors for each of the 5 classes of BFs. In each sub-figure, the upper triangular part displays the pairwise Jensen-Shannon distances of those distributions. The lower triangular part represents the p-values of the one-sided K-S test (cumulative distribution of the label of column greater than that of the row). Entries shown as 0 are actually very small, with values less than $10^{-307}$.

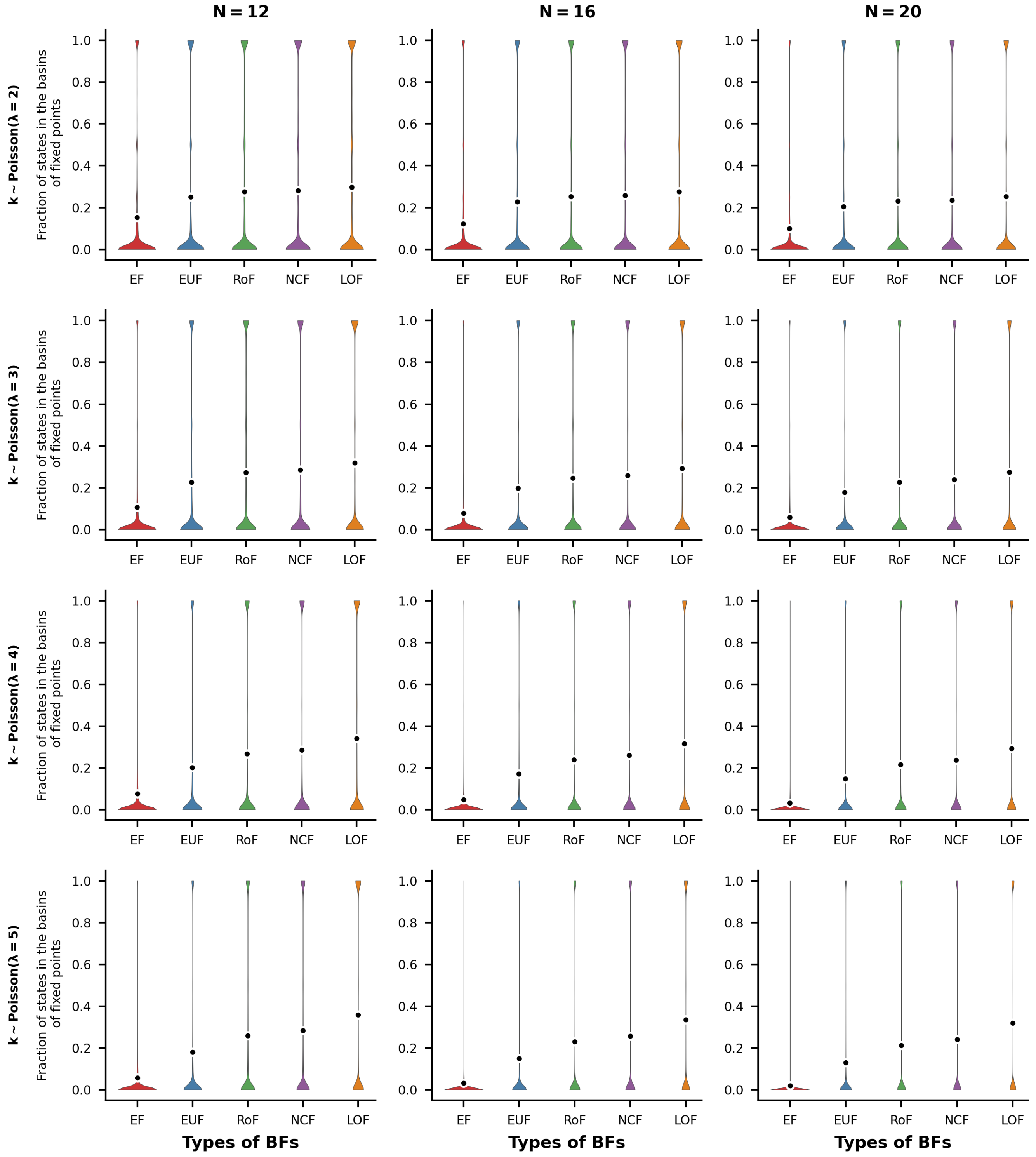


FIG. S40. **Distributions of Fraction of states in the basins of fixed points for P-P networks** The violin plots display the distribution of the fraction of states in the basins of fixed point attractors for the P-P network topology. The sub-figures are organized into 3 columns, corresponding to the network sizes $N = 12, 16$ and 20 respectively. Each row represents a different average number of inputs per node, with the degrees drawn from Poisson distributions with means $2, 3, 4$ and 5. Within each sub-figure, there are five violins representing the distributions of the fraction of states in the basins of fixed point attractors for the different ensembles of models, namely EF, EUF, RoF, NCF and LOF. Each violin distribution is based on $10^6$ data points. Outliers of each distribution are shown as gray rings while the means are shown as black dots encircled with white rings.

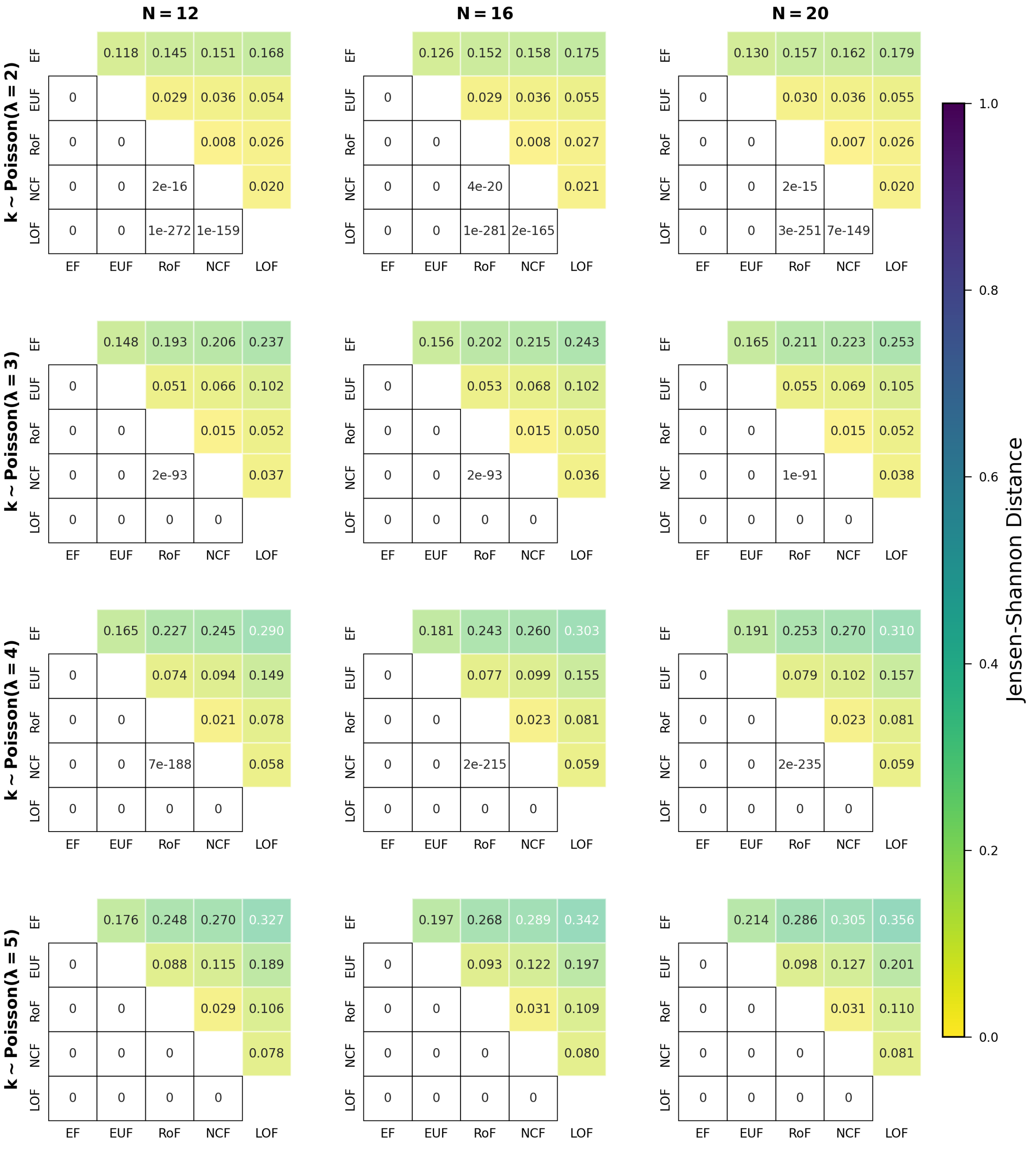


FIG. S41. **Comparing distributions for the fraction of states in the basins of fixed points when using different classes of BFs within the P-P networks.** The different sub-figures are organised according to the values of $N$ (the number of nodes in the network) and $\lambda$ ($\lambda = 2, 3, 4$ and 5). The degrees of each node of the networks are drawn randomly from a Poisson distributions with mean $= \lambda$. At a given $\lambda$ and $N$, we determined the distribution of the fraction of states in the basins of fixed points for each of the 5 classes of BFs. In each sub-figure, the upper triangular part displays the pairwise Jensen-Shannon distances of those distributions. The lower triangular part represents the p-values of the one-sided K-S test (cumulative distribution of the label of column greater than that of the row). Entries shown as 0 are actually very small, with values less than $10^{-307}$.

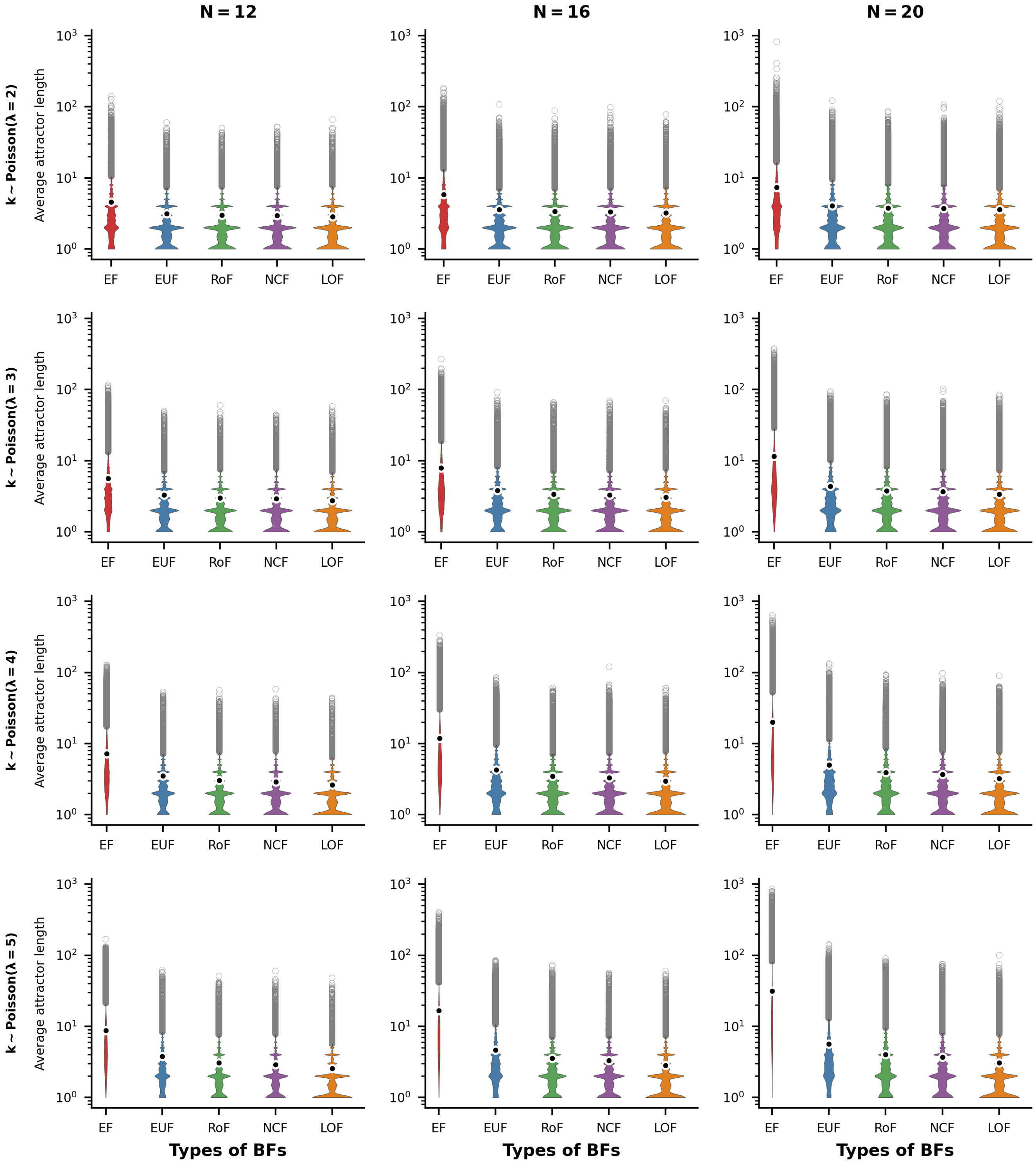


FIG. S42. **Distributions of average attractor lengths for P-P networks.** The violin plots display the distribution of the average attractor length for the P-P network topology. The sub-figures are organized into 3 columns, corresponding to the network sizes $N = 12, 16$ and 20 respectively. Each row represents a different average number of inputs per node, with the degrees drawn from Poisson distributions with means $2, 3, 4$ and 5. Within each sub-figure, there are five violins representing the distributions of the average attractor length for the different ensembles of models, namely EF, EUF, RoF, NCF and LOF. Each violin distribution is based on $10^6$ data points. Outliers of each distribution are shown as gray rings while the means are shown as black dots encircled with white rings. Note the logarithmic scale for the y-axis.

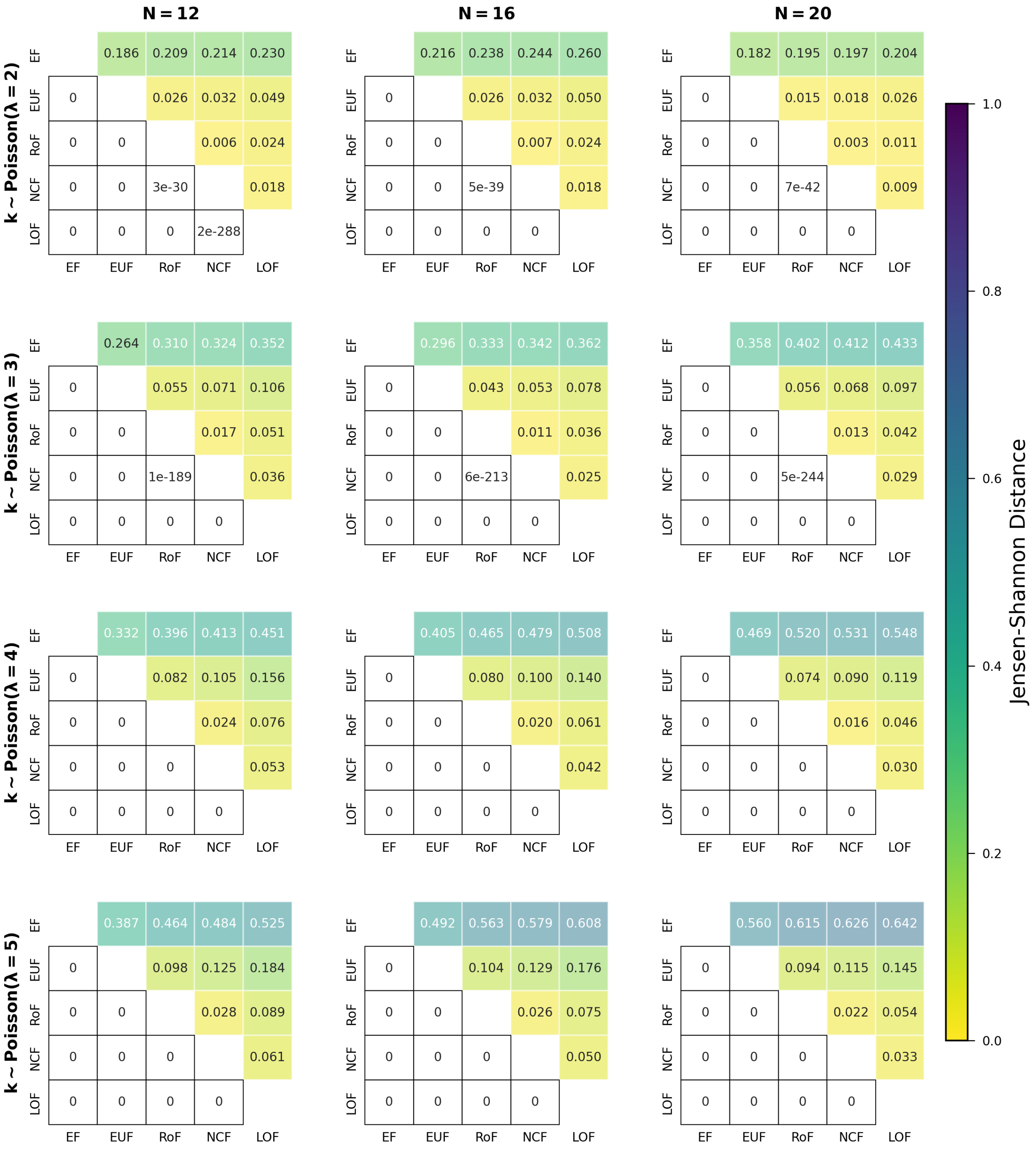


FIG. S43. **Comparing distributions for the average attractor lengths when using different classes of BFs within the P-P networks.** The different sub-figures are organised according to the values of $N$ (the number of nodes in the network) and $\lambda$ ($\lambda = 2, 3, 4$ and 5). The degrees of each node of the networks are drawn randomly from a Poisson distributions with mean $= \lambda$. At a given $\lambda$ and $N$, we determined the distribution of the average attractor lengths for each of the 5 classes of BFs. In each sub-figure, the upper triangular part displays the pairwise Jensen-Shannon distances of those distributions. The lower triangular part represents the p-values of the one-sided K-S test (cumulative distribution of the label of column less than that of the row). Entries shown as 0 are actually very small, with values less than $10^{-307}$.

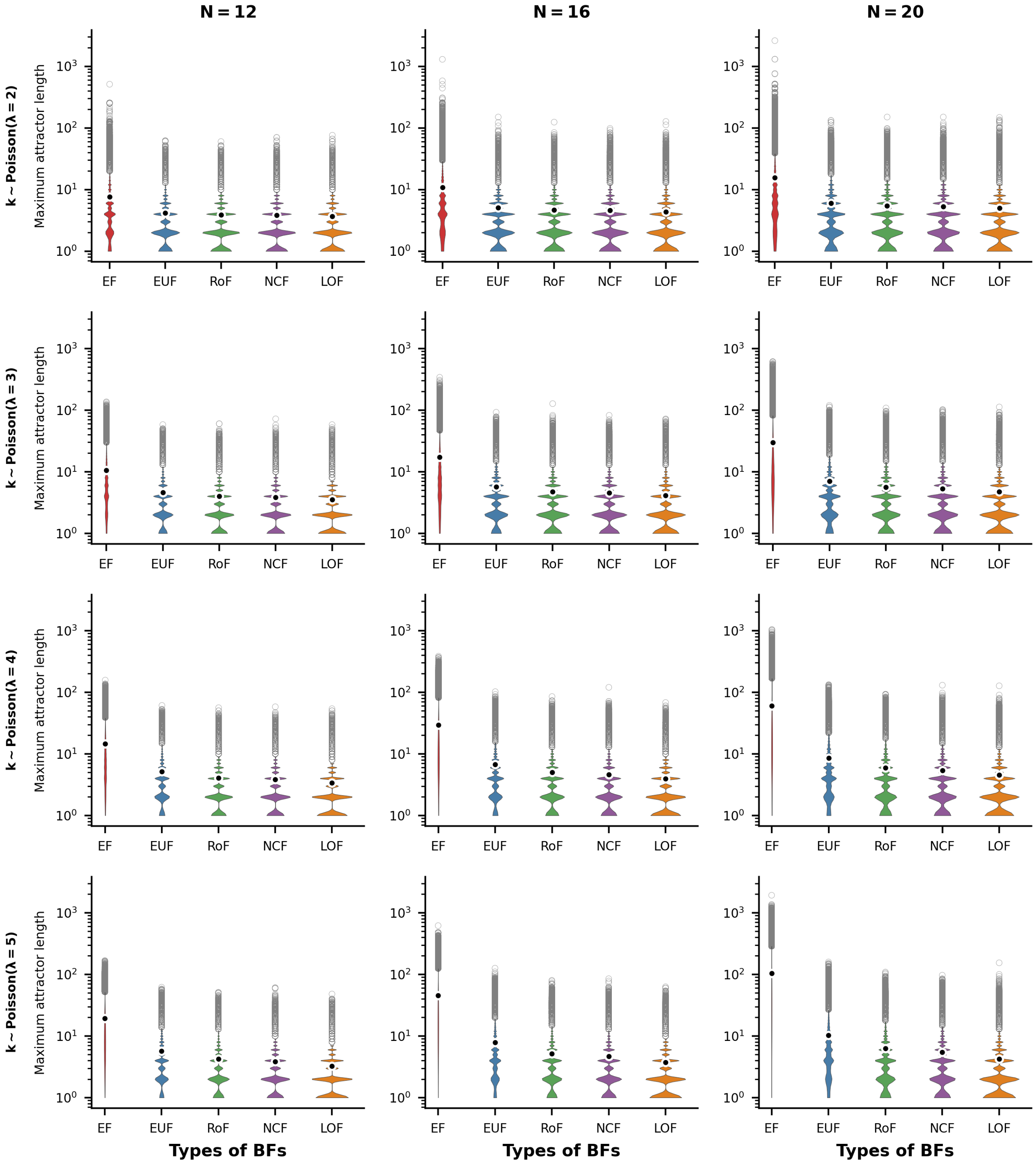


FIG. S44. **Distributions of Maximum attractor lengths for P-P networks.** The violin plots display the distribution of the maximum attractor length for the P-P network topology. The sub-figures are organized into 3 columns, corresponding to the network sizes $N = 12, 16$ and 20 respectively. Each row represents a different average number of inputs per node, with the degrees drawn from Poisson distributions with means $2, 3, 4$ and 5. Within each sub-figure, there are five violins representing the distributions of the maximum attractor length for the different ensembles of models, namely EF, EUF, RoF, NCF and LOF. Each violin distribution is based on $10^6$ data points. Outliers of each distribution are shown as gray rings while the means are shown as black dots encircled with white rings. Note the logarithmic scale for the y-axis.

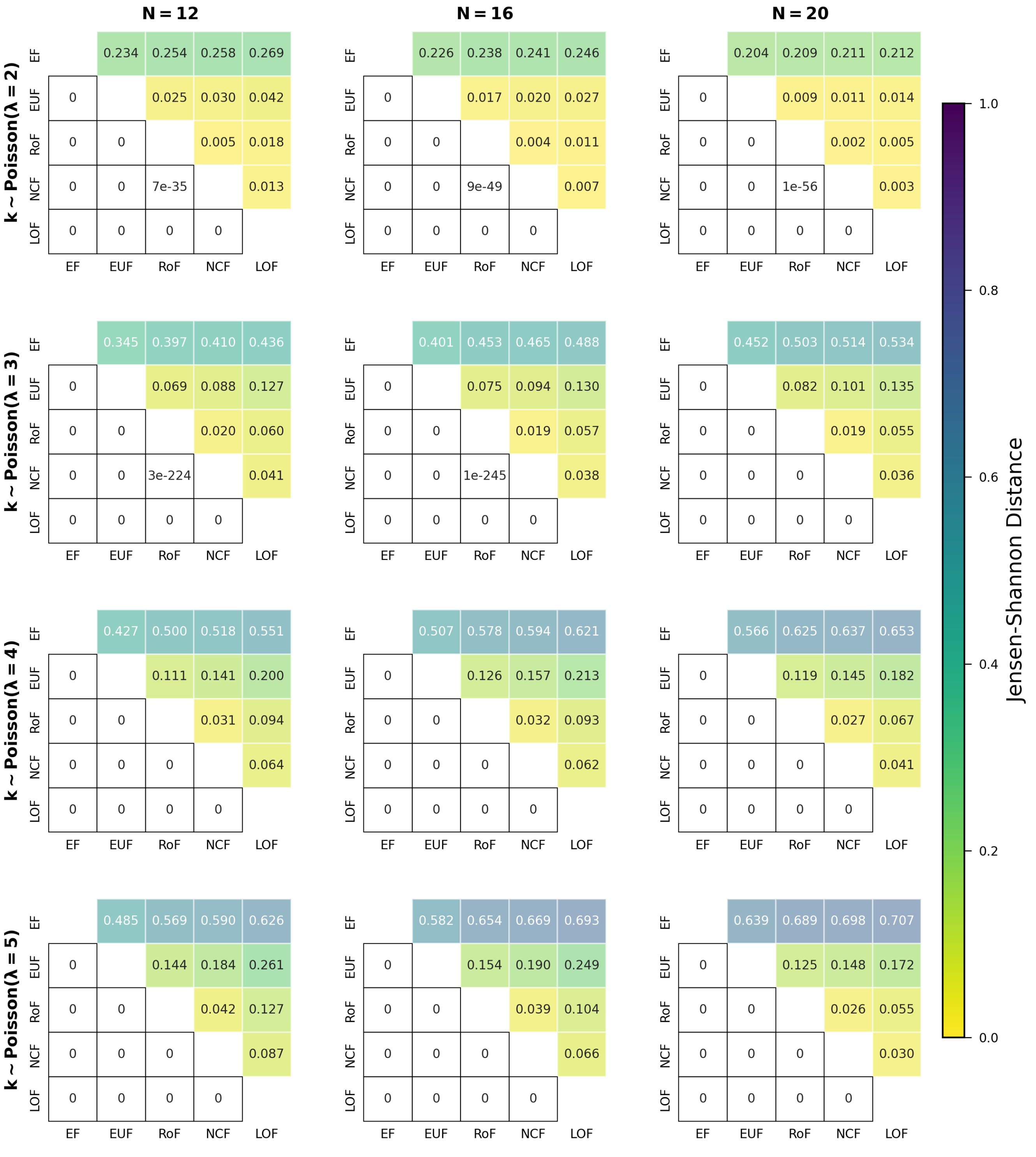


FIG. S45. **Comparing distributions for the maximum attractor lengths when using different classes of BFs within the P-P networks.** The different sub-figures are organised according to the values of $N$ (the number of nodes in the network) and $\lambda$ ($\lambda = 2, 3, 4$ and 5). The degrees of each node of the networks are drawn randomly from a Poisson distributions with mean $= \lambda$. At a given $\lambda$ and $N$, we determined the distribution of the maximum attractor lengths for each of the 5 classes of BFs. In each sub-figure, the upper triangular part displays the pairwise Jensen-Shannon distances of those distributions. The lower triangular part represents the p-values of the one-sided K-S test (cumulative distribution of the label of column less than that of the row). Entries shown as 0 are actually very small, with values less than $10^{-307}$.

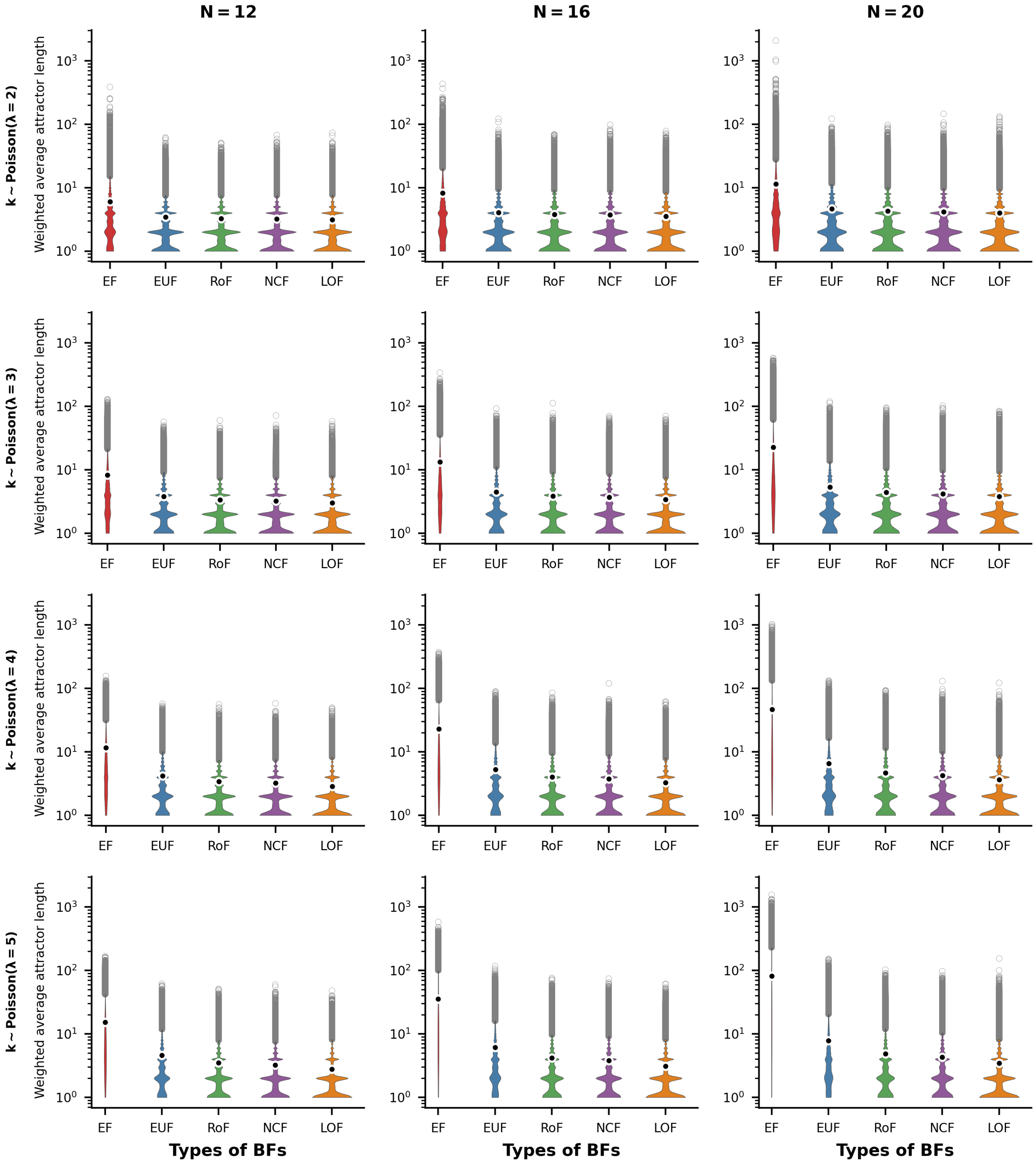


FIG. S46. **Distributions of weighted average attractor lengths for P-P networks.** The violin plots display the distribution of the weighted average attractor length for the P-P network topology. The sub-figures are organized into 3 columns, corresponding to the network sizes $N = 12, 16$ and 20 respectively. Each row represents a different average number of inputs per node, with the degrees drawn from Poisson distributions with means $2, 3, 4$ and 5. Within each sub-figure, there are five violins representing the distributions of the weighted average attractor length for the different ensembles of models, namely EF, EUF, RoF, NCF and LOF. Each violin distribution is based on $10^6$ data points. Outliers of each distribution are shown as gray rings while the means are shown as black dots encircled with white rings. Note the logarithmic scale for the y-axis.

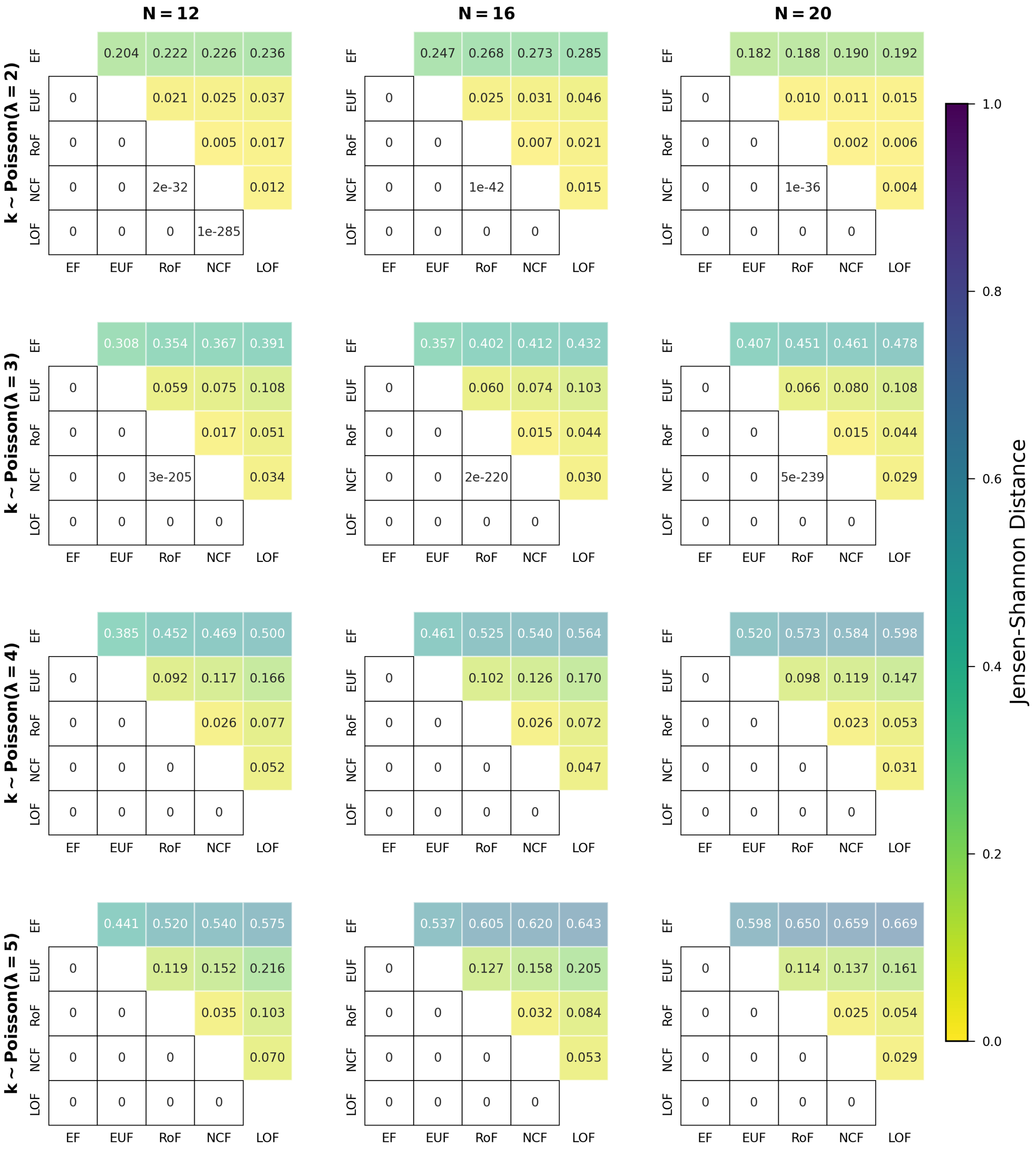

FIG. S47. **Comparing distributions for the weighted average attractor lengths when using different classes of BFs within the P-P networks.** The different sub-figures are organised according to the values of $N$ (the number of nodes in the network) and $\lambda$ ($\lambda = 2, 3, 4$ and 5). The degrees of each node of the networks are drawn randomly from a Poisson distributions with mean $= \lambda$. At a given $\lambda$ and $N$, we determined the distribution of the weighted average attractor lengths for each of the 5 classes of BFs. In each sub-figure, the upper triangular part displays the pairwise Jensen-Shannon distances of those distributions. The lower triangular part represents the p-values of the one-sided K-S test (cumulative distribution of the label of column less than that of the row). Entries shown as 0 are actually very small, with values less than $10^{-307}$.

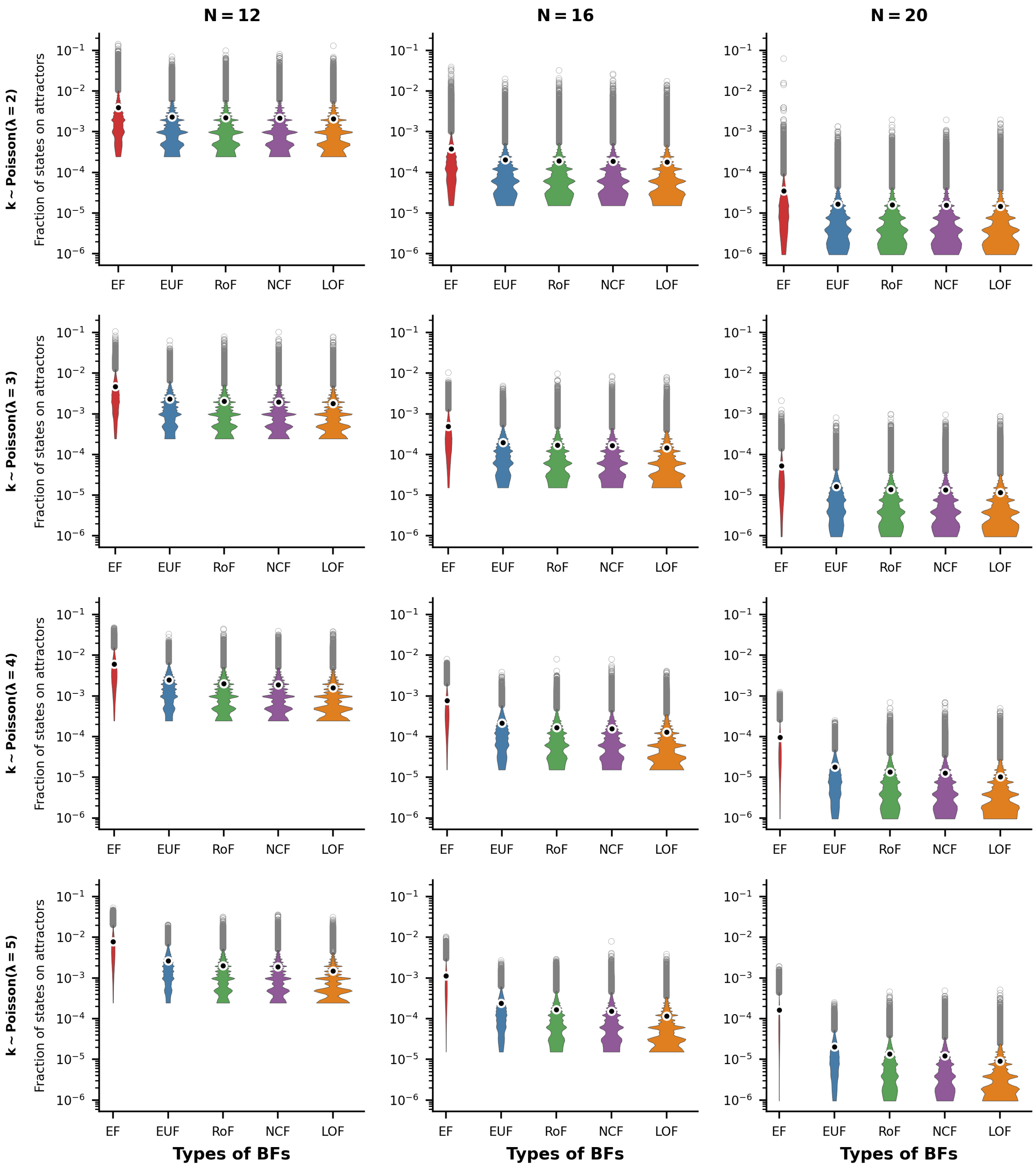


FIG. S48. **Distributions of fraction of states on attractors for P-P networks.** The violin plots display the distribution of the fraction of states on attractors for the P-P network topology. The sub-figures are organized into 3 columns, corresponding to the network sizes $N = 12, 16$ and 20 respectively. Each row represents a different average number of inputs per node, with the degrees drawn from Poisson distributions with means $2, 3, 4$ and 5. Within each sub-figure, there are five violins representing the distributions of the fraction of states on attractors for the different ensembles of models, namely EF, EUF, RoF, NCF and LOF. Each violin distribution is based on $10^6$ data points. Outliers of each distribution are shown as gray rings while the means are shown as black dots encircled with white rings. Note the logarithmic scale for the y-axis.

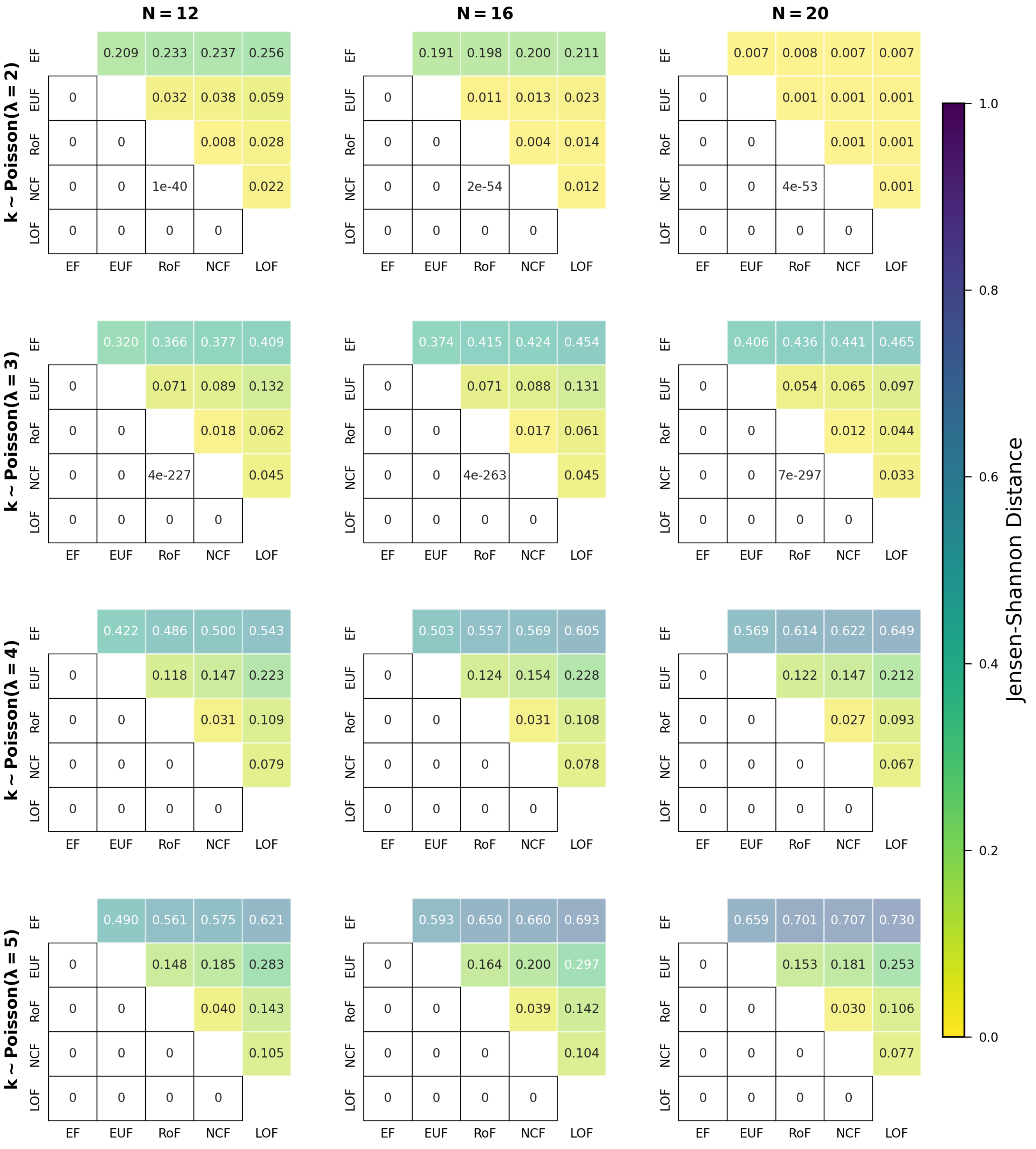


FIG. S49. **Comparing distributions for the fraction of states on attractors when using different classes of BFs within the P-P networks.** The different sub-figures are organised according to the values of $N$ (the number of nodes in the network) and $\lambda$ ($\lambda = 2, 3, 4$ and 5). The degrees of each node of the networks are drawn randomly from a Poisson distributions with mean $= \lambda$. At a given $\lambda$ and $N$, we determined the distribution of the fraction of states on attractors for each of the 5 classes of BFs. In each sub-figure, the upper triangular part displays the pairwise Jensen-Shannon distances of those distributions. The lower triangular part represents the p-values of the one-sided K-S test (cumulative distribution of the label of column less than that of the row). Entries shown as 0 are actually very small, with values less than $10^{-307}$.

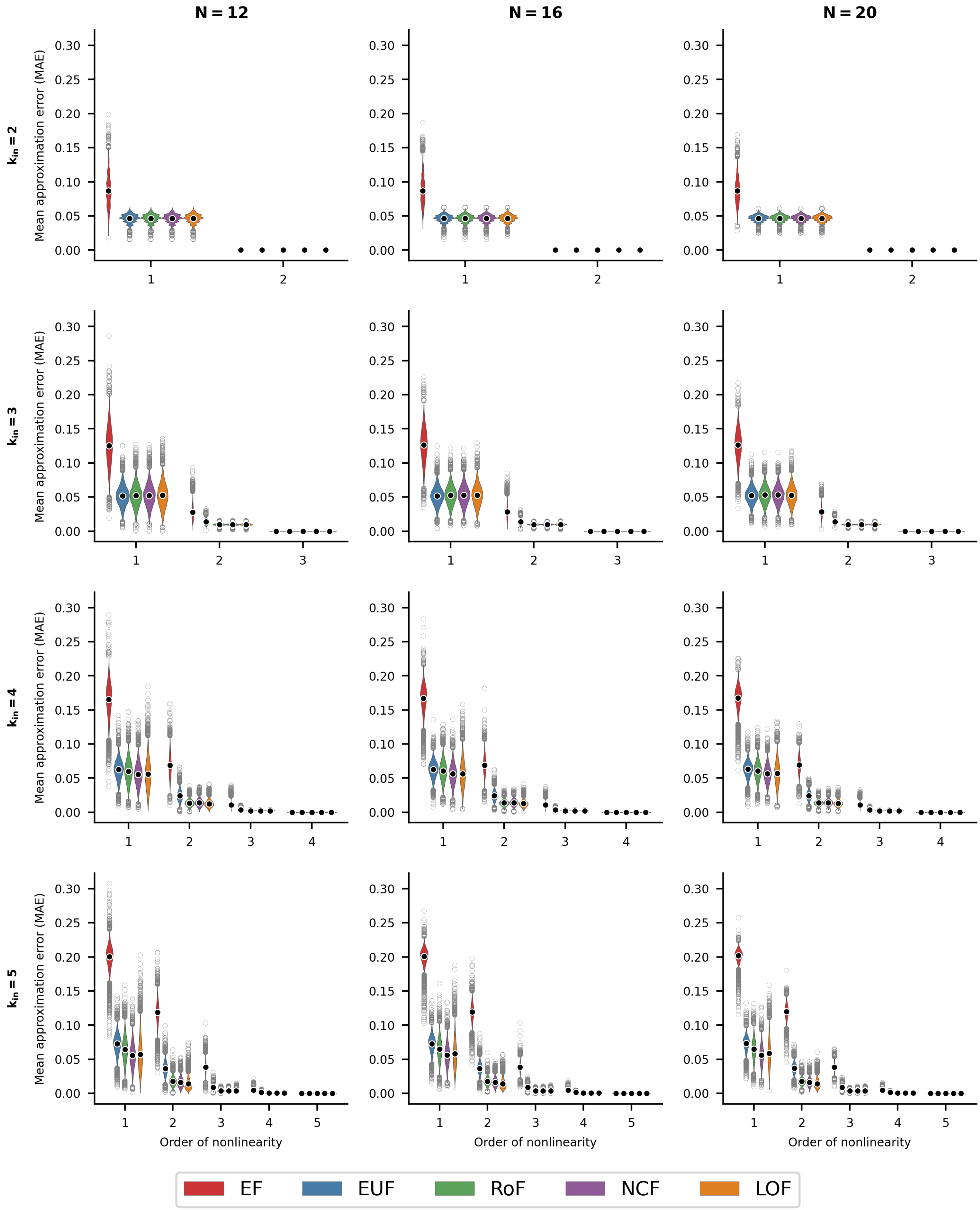


FIG. S50. **Distributions of mean approximation error (MAE) for R-P networks.** The violin plots display the distribution of the MAE for the R-P network topology. The sub-figures are organized into 3 columns, corresponding to the network sizes N=12, 16 and 20 respectively. Each row represents a different number of inputs per node, ranging from 2 to 5. Within each sub-figure, there are grouped violins that show the distributions for the different ensembles of models, namely EF, EUF, RoF, NCF and LOF for different orders of nonlinearity. Each violin distribution is based on $10^4$ data points. Outliers of each distribution are shown as gray rings while the means are shown as black dots encircled with white rings.

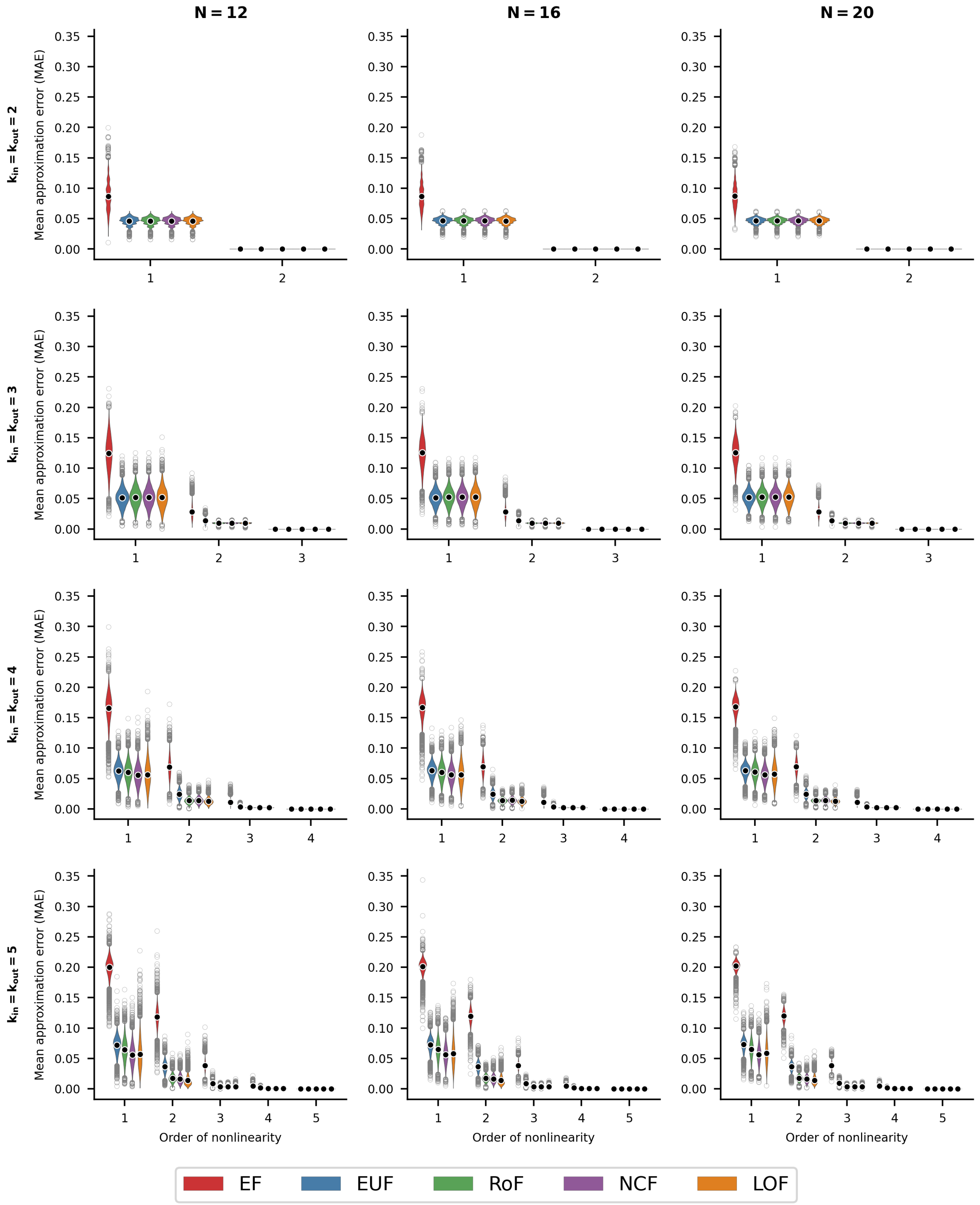


FIG. S51. **Distributions of mean approximation error (MAE) for R-R networks.** The violin plots display the distribution of the MAE for the R-R network topology. The sub-figures are organized into 3 columns, corresponding to the network sizes N=12, 16 and 20 respectively. Each row represents a different number of inputs per node, ranging from 2 to 5. Within each sub-figure, there are grouped violins that show the distributions for the different ensembles of models, namely EF, EUF, RoF, NCF and LOF for different orders of nonlinearity. Each violin distribution is based on $10^4$ data points. Outliers of each distribution are shown as gray rings while the means are shown as black dots encircled with white rings.

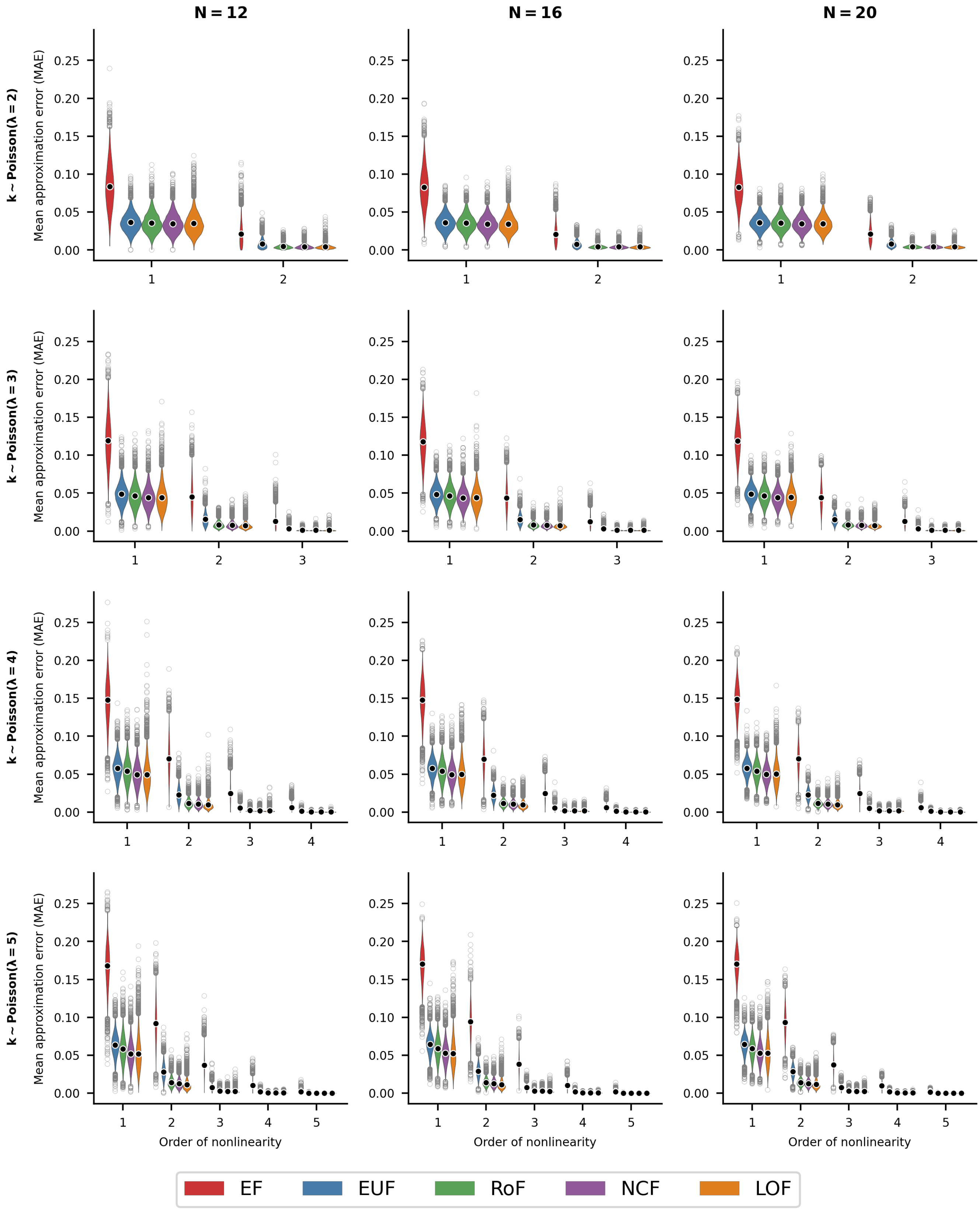


FIG. S52. **Distributions of mean approximation error (MAE) for P-P networks.** The violin plots display the distribution of the MAE for the P-P network topology. The sub-figures are organized into 3 columns, corresponding to the network sizes N=12, 16 and 20 respectively. Each row represents a different average number of inputs per node, with the degrees drawn from Poisson distributions with means 2, 3, 4 and 5. Within each sub-figure, there are grouped violins that show the distributions for the different ensembles of models, namely EF, EUF, RoF, NCF and LOF for different orders of nonlinearity. Each violin distribution is based on $10^4$ data points. Outliers of each distribution are shown as gray rings while the means are shown as black dots encircled with white rings.

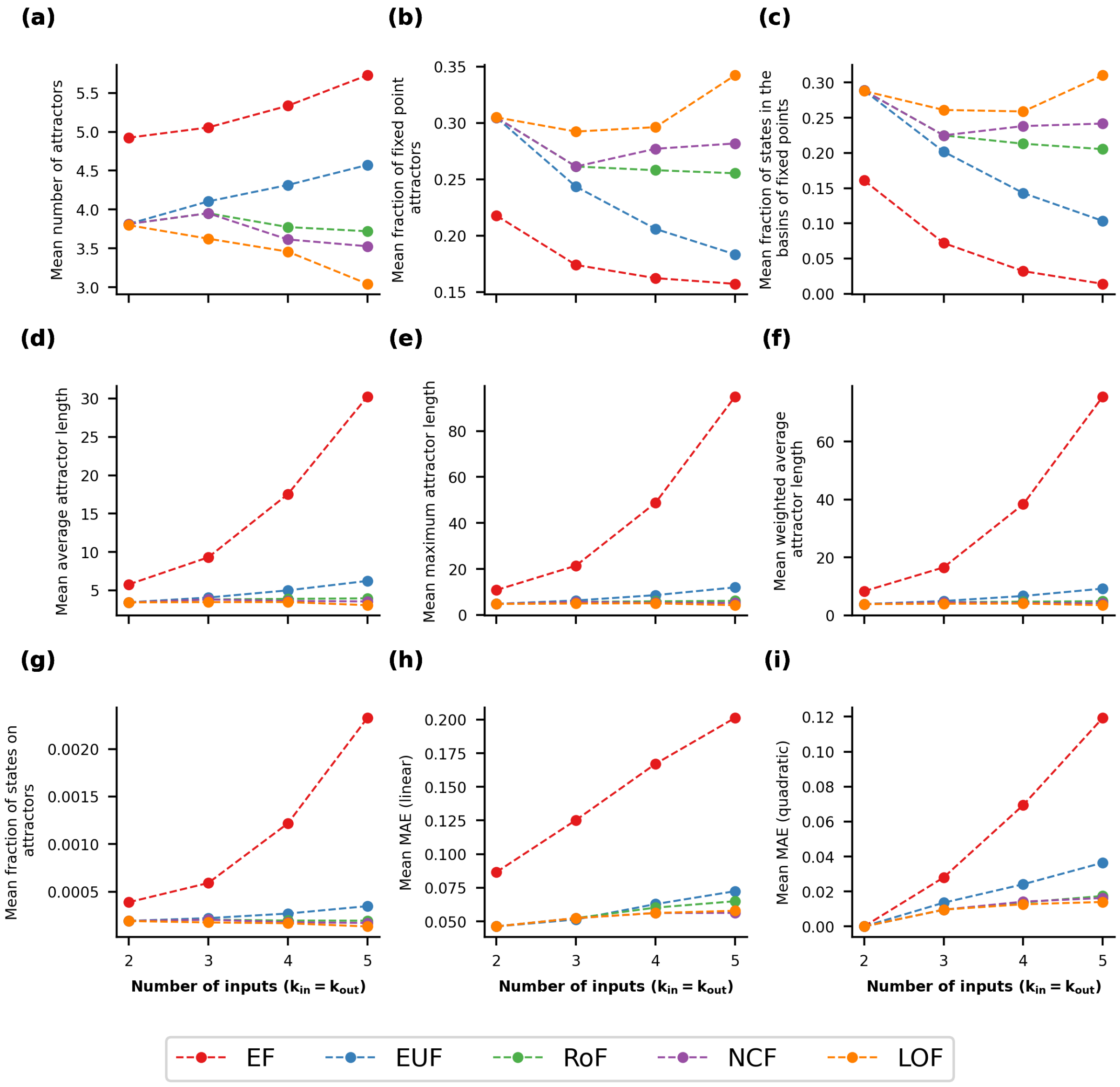


FIG. S53. **Mean values of 9 stability observables as a function of $k_{in}(= k_{out})$ for different classes of BFs, in the case of R-R networks at $N = 16$.** Within each sub-figure, the 5 curves correspond to the five different classes of BFs. The data displayed represent the mean value of the corresponding observable at a given value of $k_{in}$ for R-R network topology.

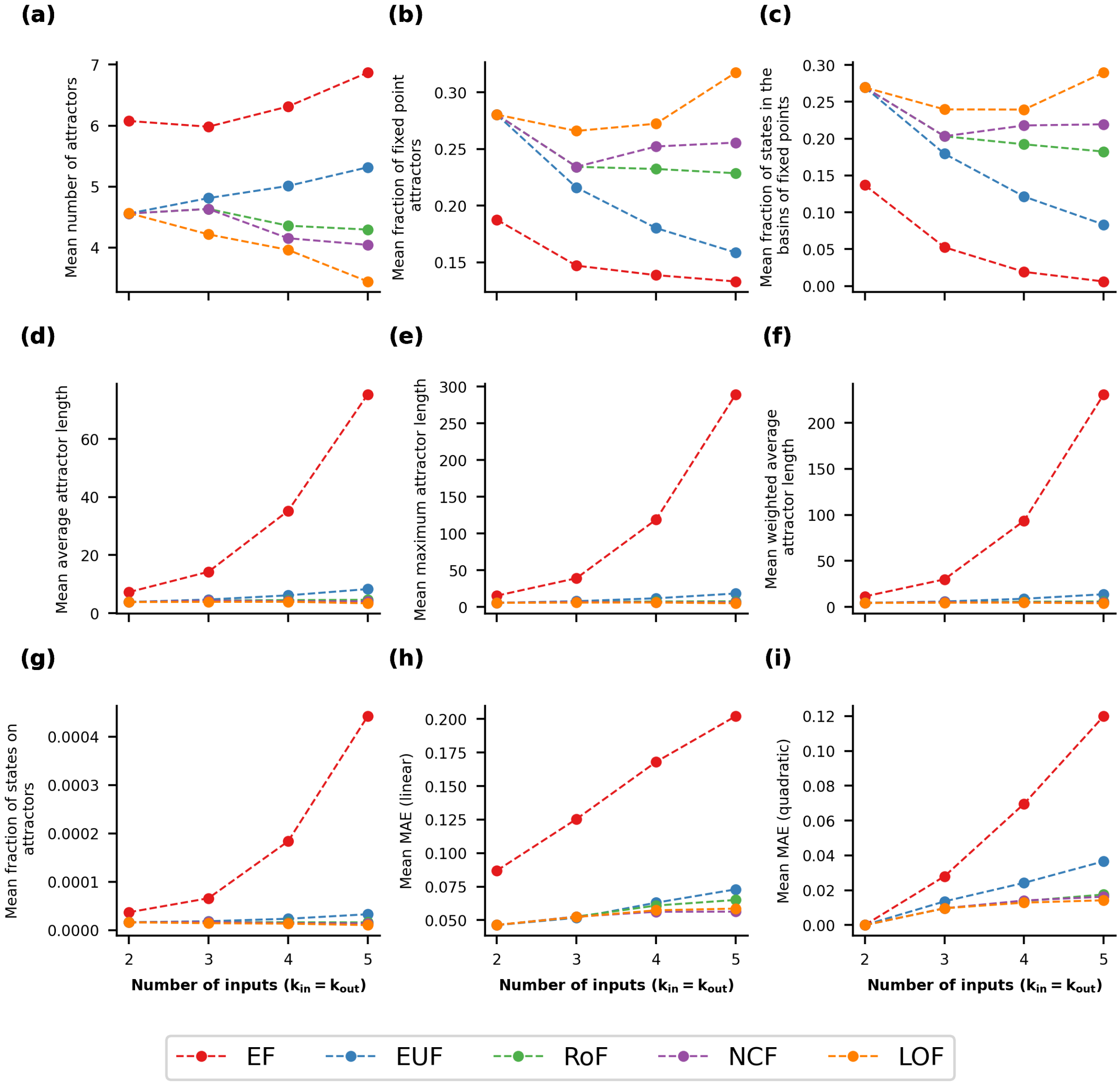


FIG. S54. **Mean values of 9 stability observables as a function of $k_{in}(= k_{out})$ for different classes of BFs, in the case of R-R networks at $N = 20$.** Within each sub-figure, the 5 curves correspond to the five different classes of BFs. The data displayed represent the mean value of the corresponding observable at a given value of $k_{in}$ for R-R network topology.

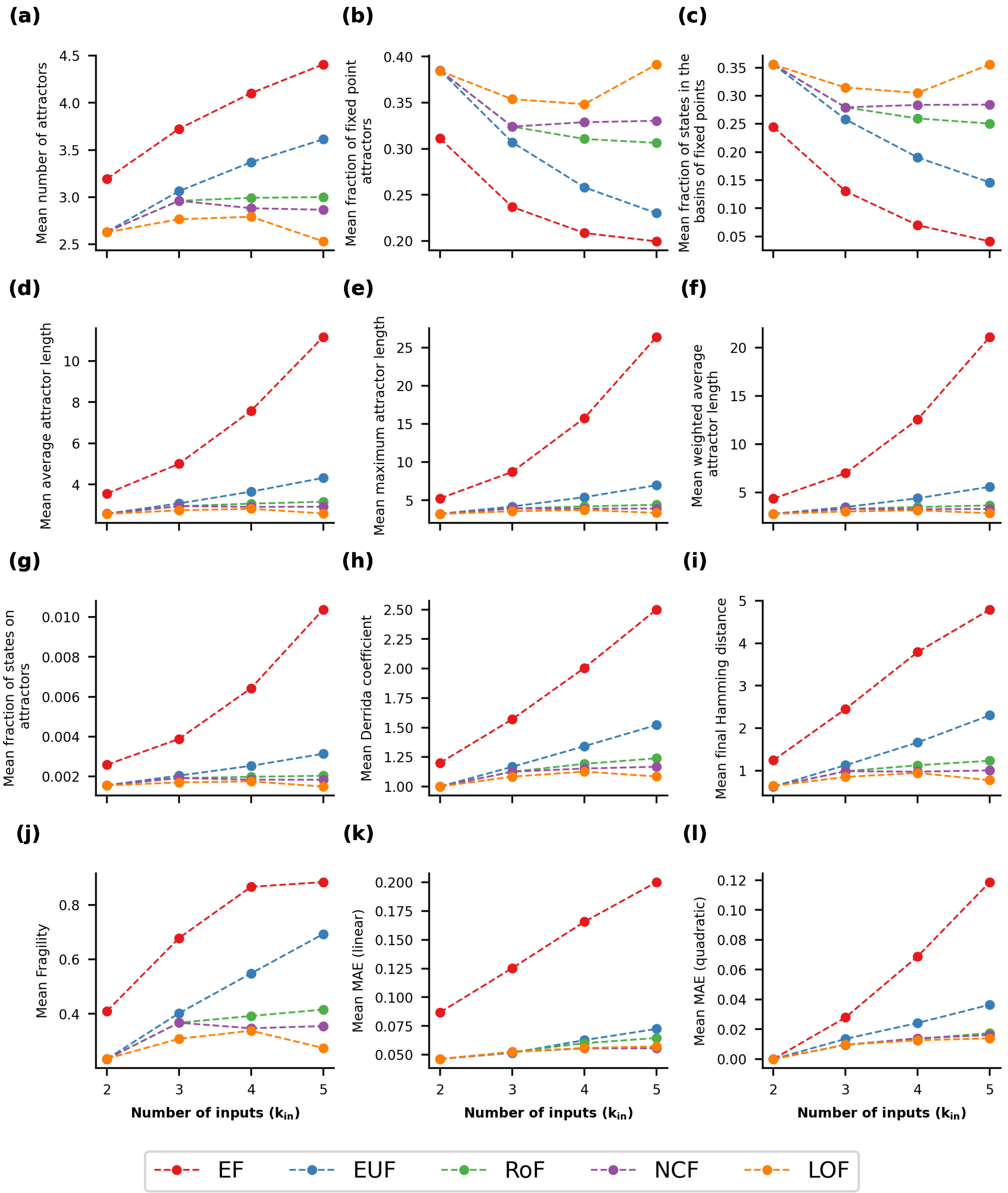


FIG. S55. **Mean values of our 12 stability observables as a function of $k_{in}$ for different classes of BFs, in the case of R-P networks at $N = 12$.** Within each sub-figure, the 5 curves correspond to the five different classes of BFs. The data displayed represent the mean value of the corresponding observable at a given value of $k_{in}$ for R-P network topology.

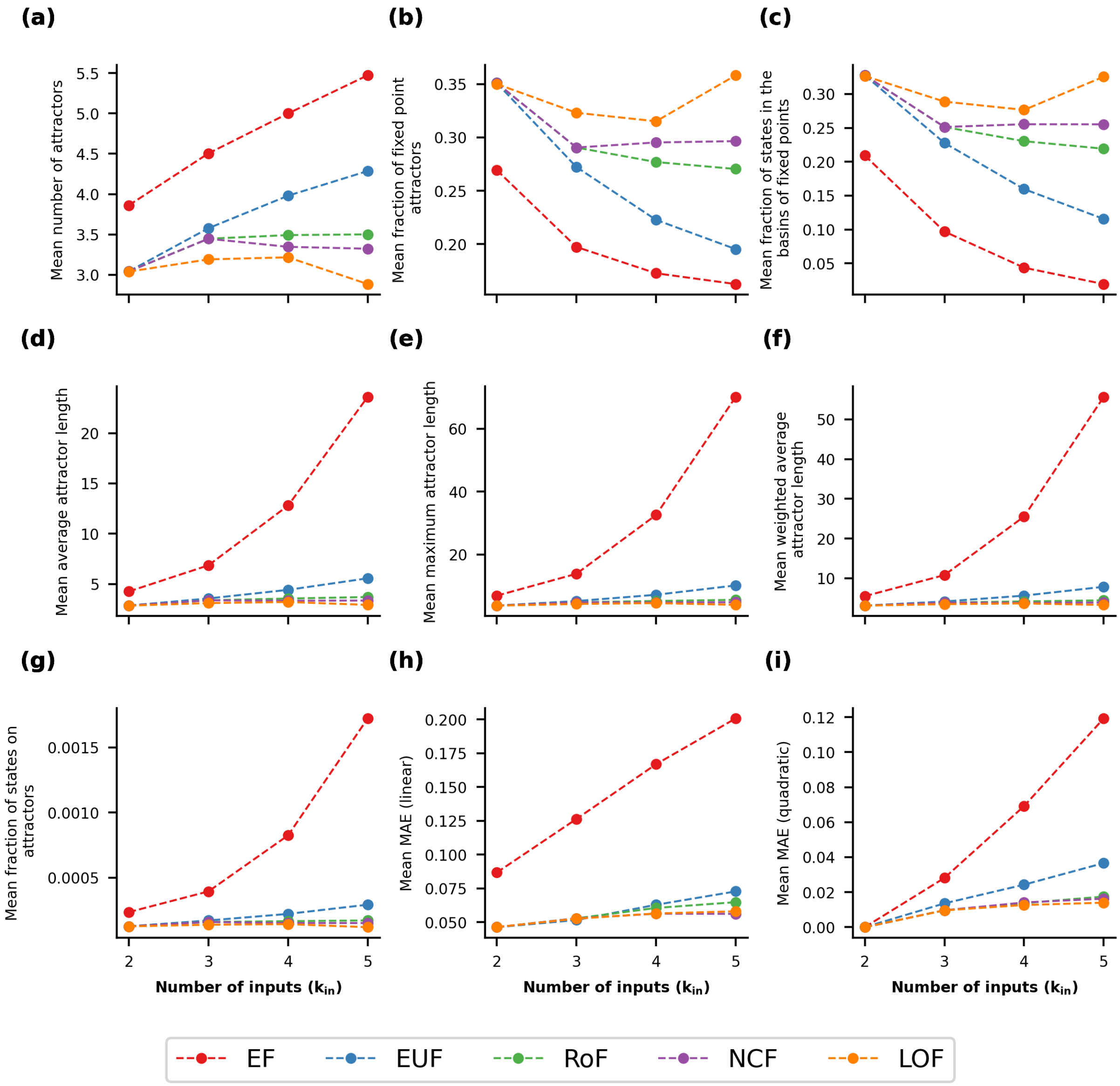


FIG. S56. **Mean values of 9 stability observables as a function of $k_{in}$ for different classes of BFs, in the case of R-P networks at $N = 16$.** Within each sub-figure, the 5 curves correspond to the five different classes of BFs. The data displayed represent the mean value of the corresponding observable at a given value of $k_{in}$ for R-P network topology.

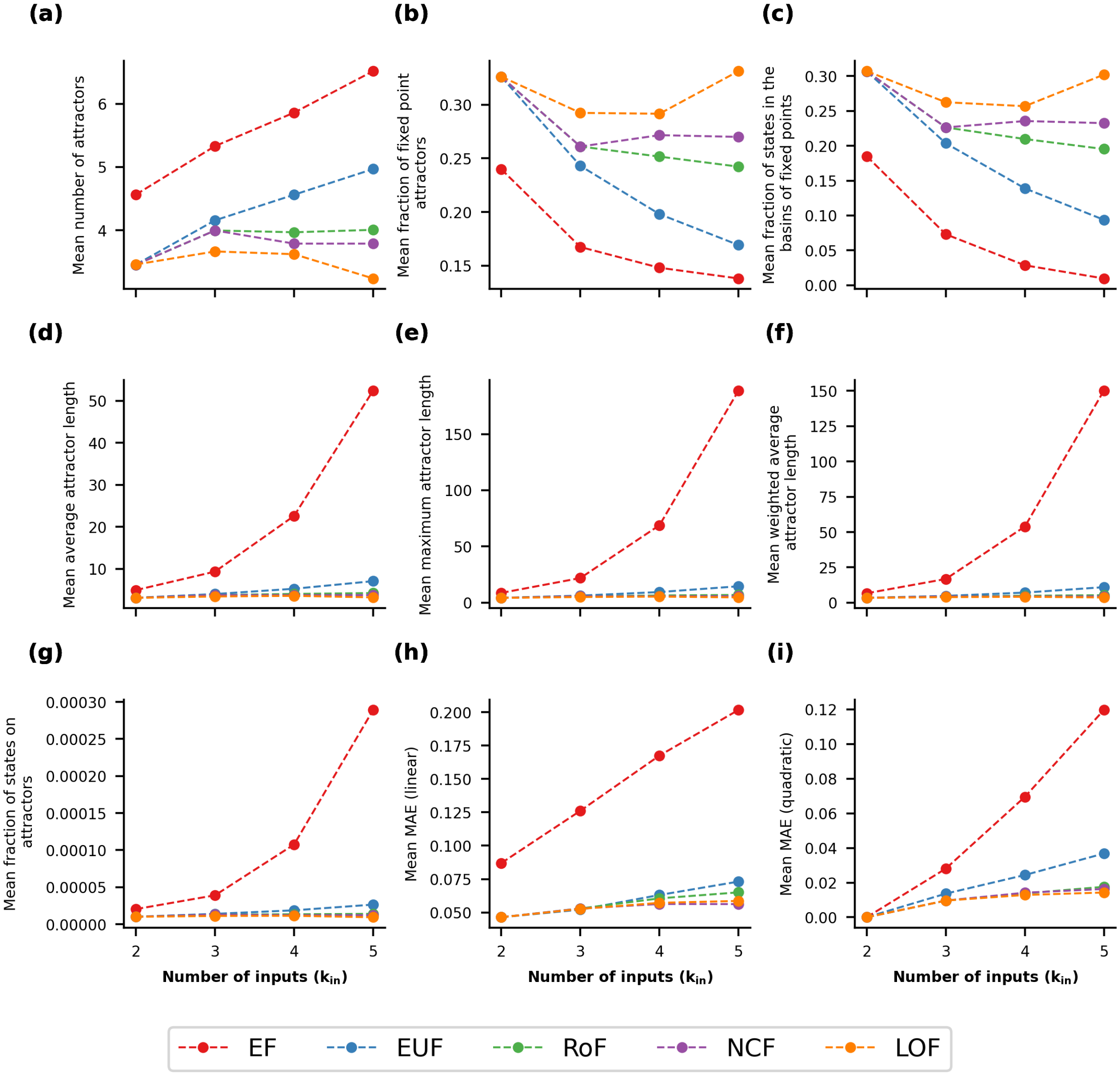


FIG. S57. **Mean values of 9 stability observables as a function of $k_{in}$ for different classes of BFs, in the case of R-P networks at $N = 20$.** Within each sub-figure, the 5 curves correspond to the five different classes of BFs. The data displayed represent the mean value of the corresponding observable at a given value of $k_{in}$ for R-P network topology.

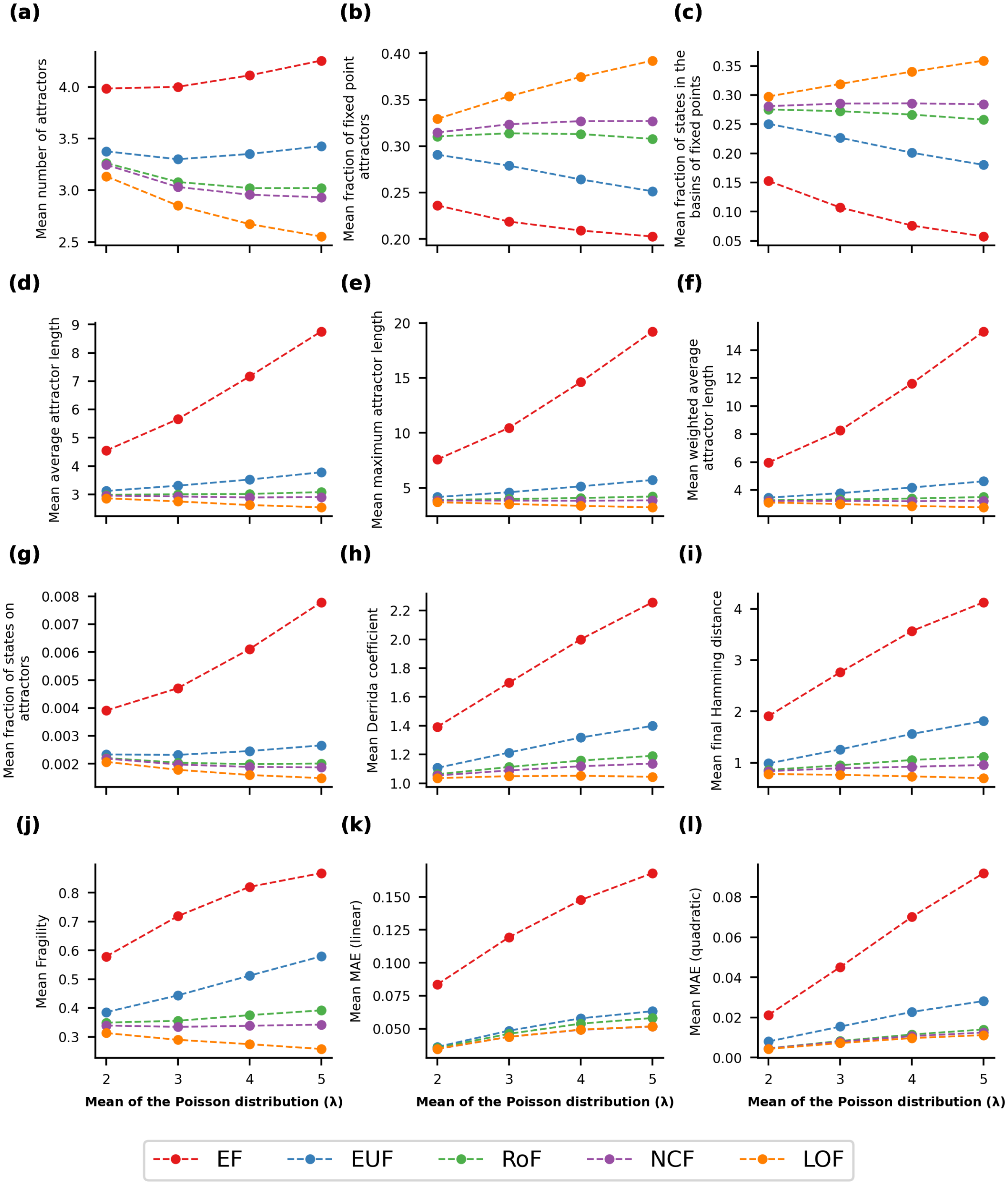


FIG. S58. **Mean values of our 12 stability observables as a function of $\lambda$ for different classes of BFs, in the case of P-P networks at $N = 12$.** Within each sub-figure, the 5 curves correspond to the five different classes of BFs. The data displayed represent the mean value of the corresponding observable at a given value of $\lambda$ (the degrees of each node in the networks are drawn randomly from a Poisson distribution with mean = $\lambda$) for P-P network topology.

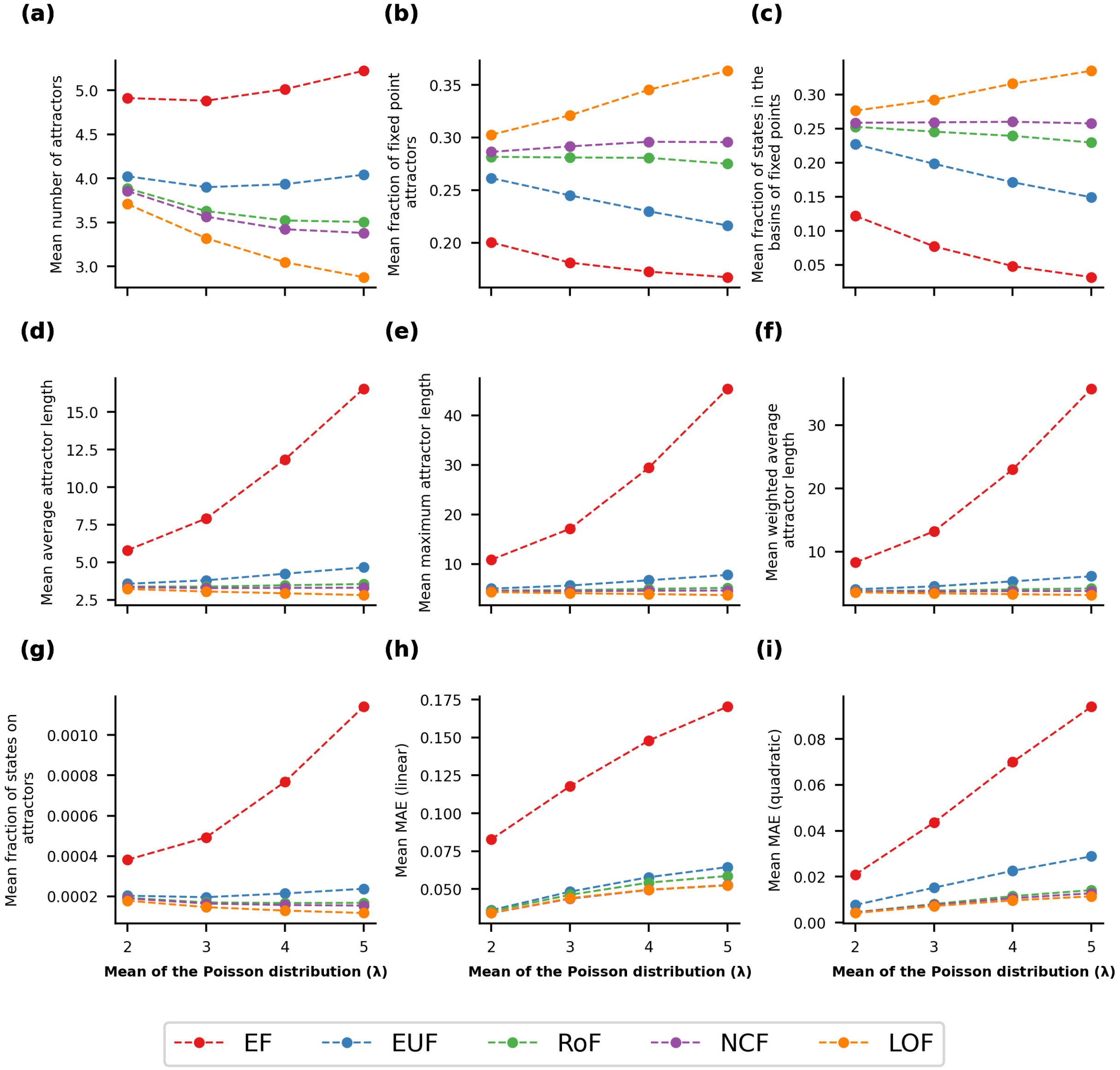


FIG. S59. **Mean values of 9 stability observables as a function of $\lambda$ for different classes of BFs, in the case of P-P networks at $N = 16$.** Within each sub-figure, the 5 curves correspond to the five different classes of BFs. The data displayed represent the mean value of the corresponding observable at a given value of $\lambda$ (the degrees of each node in the networks are drawn randomly from a Poisson distribution with mean = $\lambda$) for P-P network topology.

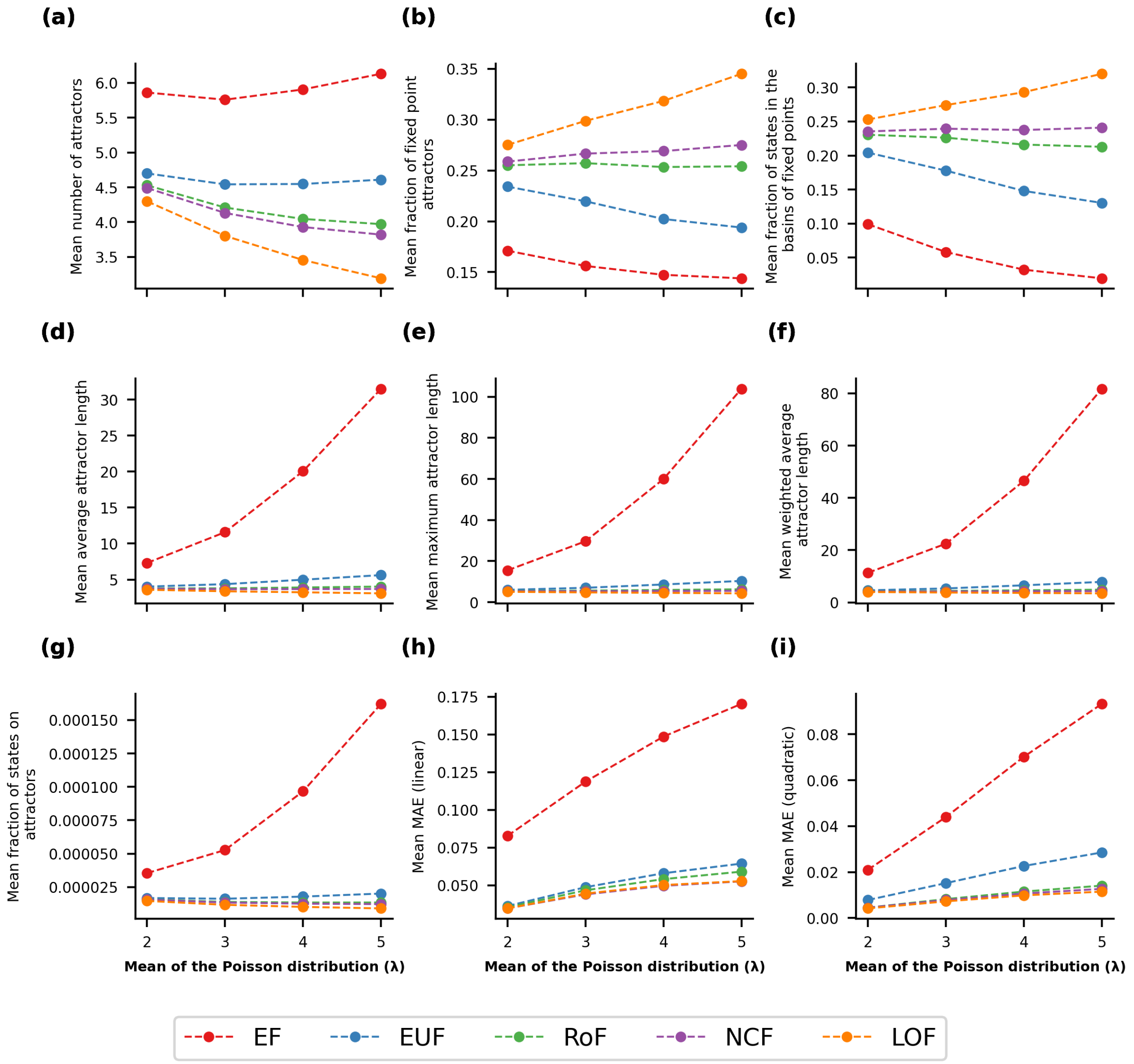


FIG. S60. **Mean values of 9 stability observables as a function of $\lambda$ for different classes of BFs, in the case of P-P networks at $N = 20$.** Within each sub-figure, the 5 curves correspond to the five different classes of BFs. The data displayed represent the mean value of the corresponding observable at a given value of $\lambda$ (the degrees of each node in the networks are drawn randomly from a Poisson distribution with mean = $\lambda$) for P-P network topology.

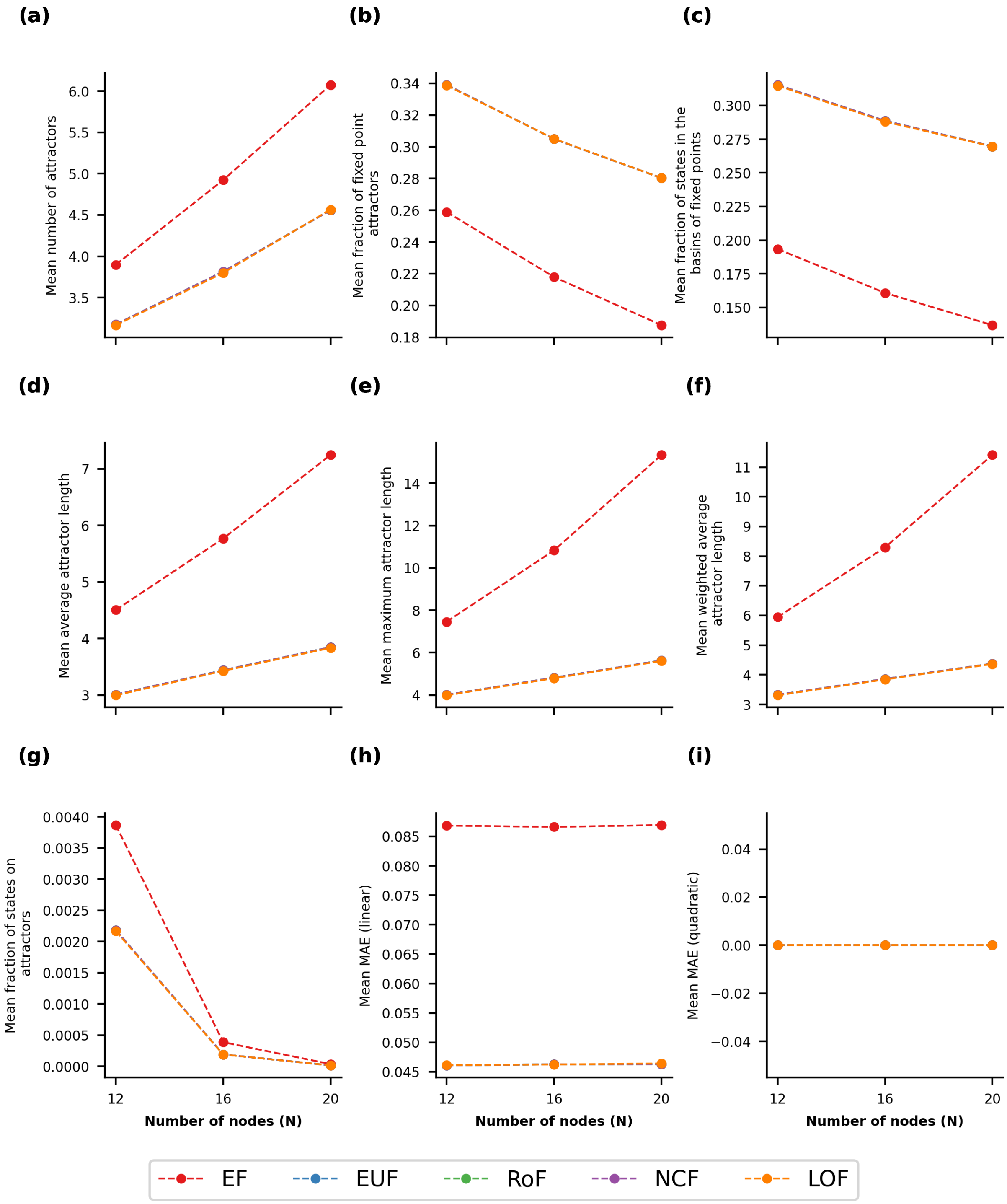


FIG. S61. **Mean of the stability observables for different values of $N$ in the case of the R-R topology with $k_{in} = k_{out} = 2$.** Within each sub-figure, the 5 curves correspond to the 5 different classes of BFs. The data displayed represent the mean value of the corresponding observable at a given value of $N$ for the R-R network topology with $k_{in} = k_{out} = 2$

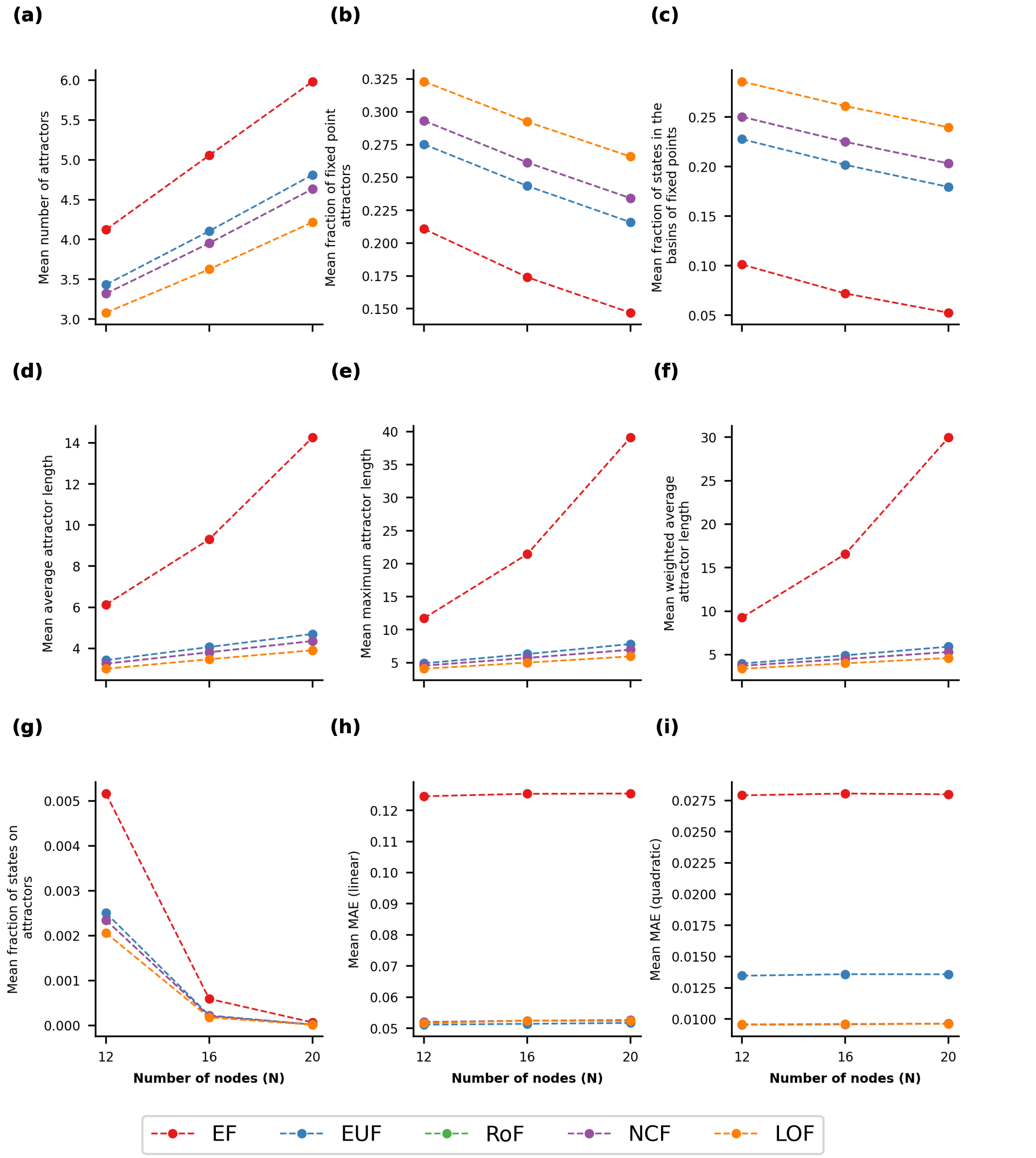


FIG. S62. **Mean of the stability observables for different values of $N$ in the case of the R-R topology with $k_{in} = k_{out} = 3$.** Within each sub-figure, the 5 curves correspond to the 5 different classes of BFs. The data displayed represent the mean value of the corresponding observable at a given value of $N$ for the R-R network topology with $k_{in} = k_{out} = 3$

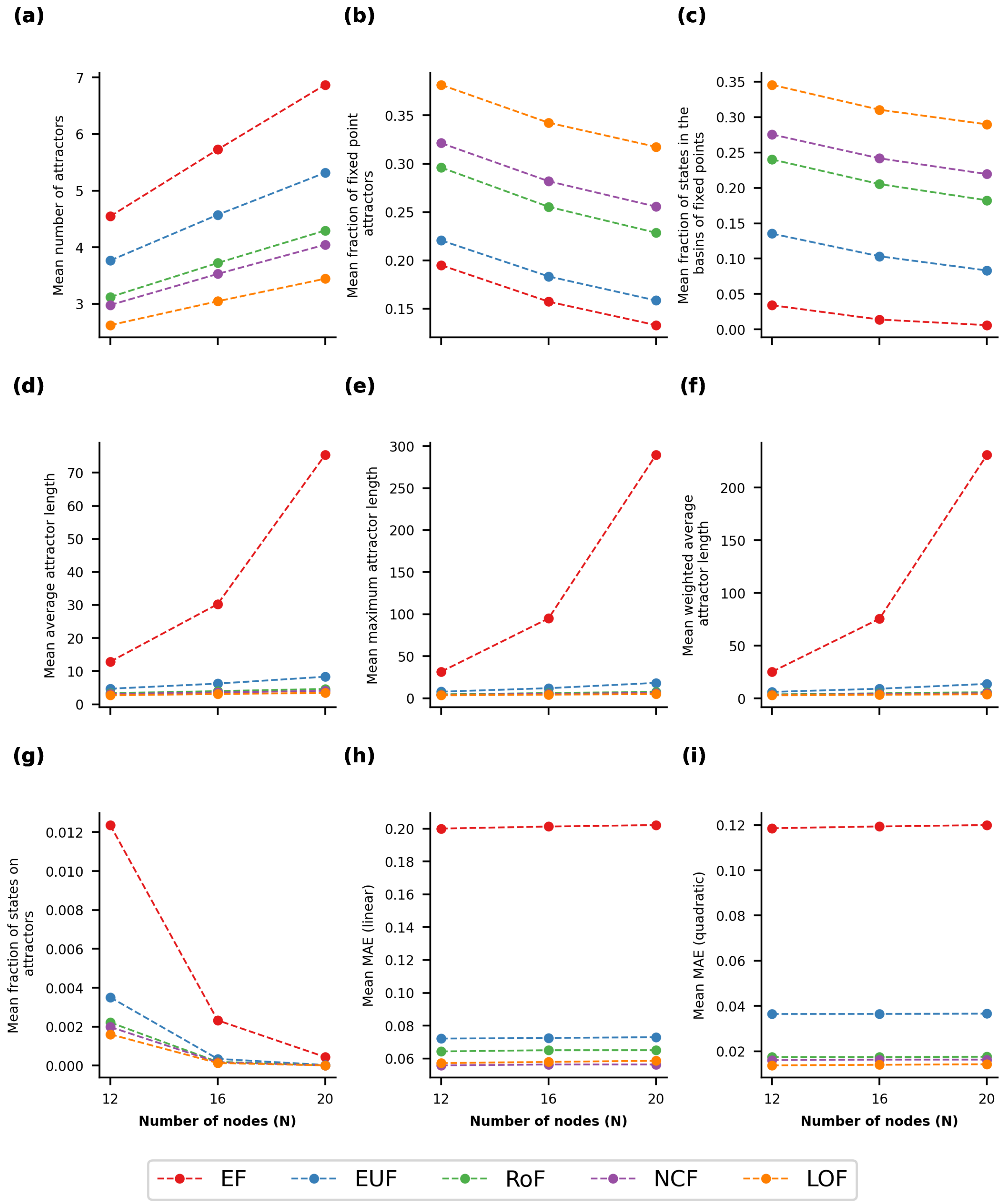


FIG. S63. **Mean of the stability observables for different values of $N$ in the case of the R-R topology with $k_{in} = k_{out} = 5$.** Within each sub-figure, the 5 curves correspond to the 5 different classes of BFs. The data displayed represent the mean value of the corresponding observable at a given value of $N$ for the R-R network topology with $k_{in} = k_{out} = 5$

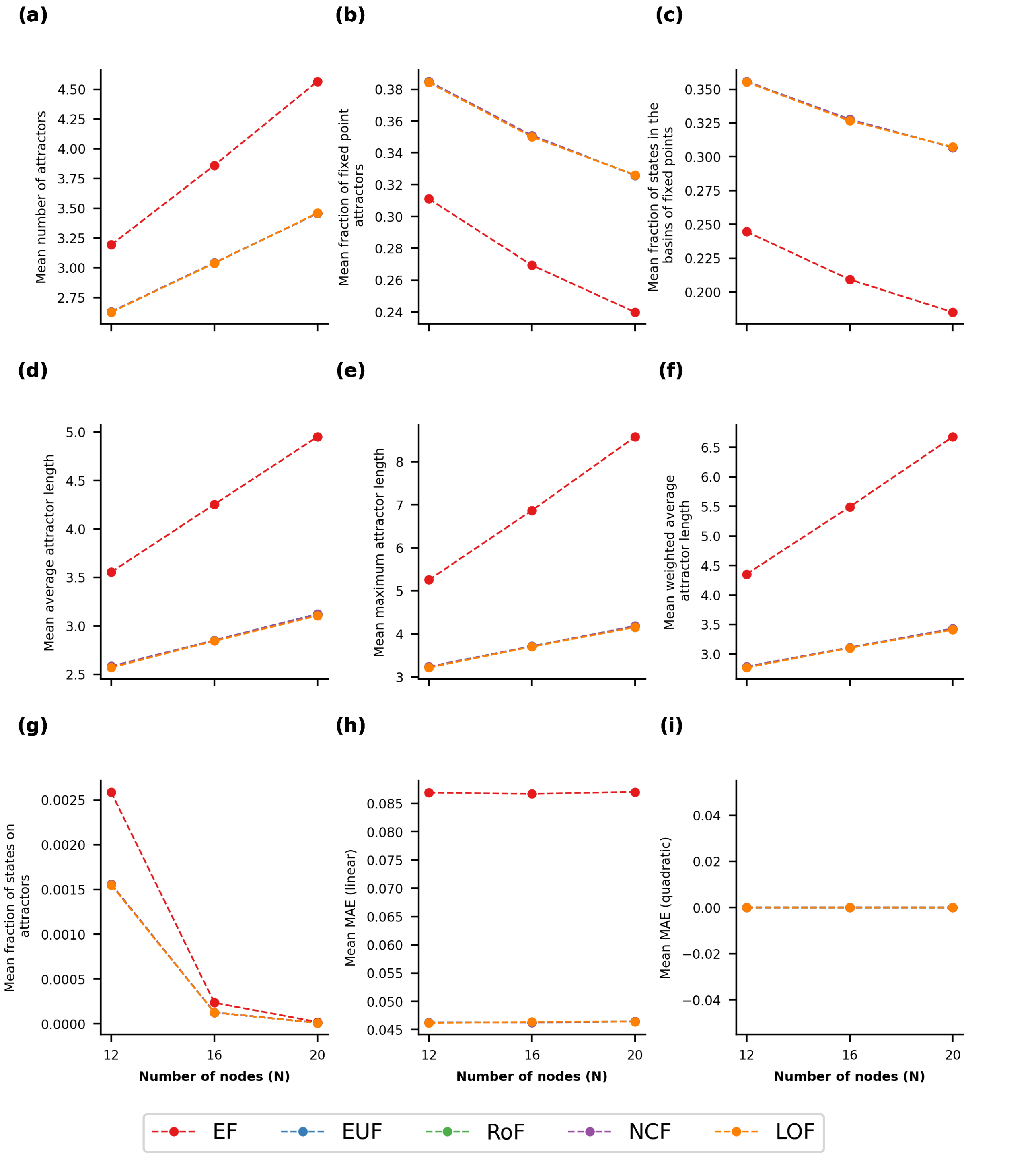


FIG. S64. **Mean of the stability observables for different values of $N$ in the case of the R-P topology with $k_{in} = 2$.** Within each sub-figure, the 5 curves correspond to the 5 different classes of BFs. The data displayed represent the mean value of the corresponding observable at a given value of $N$ for the R-P network topology with $k_{in} = 2$

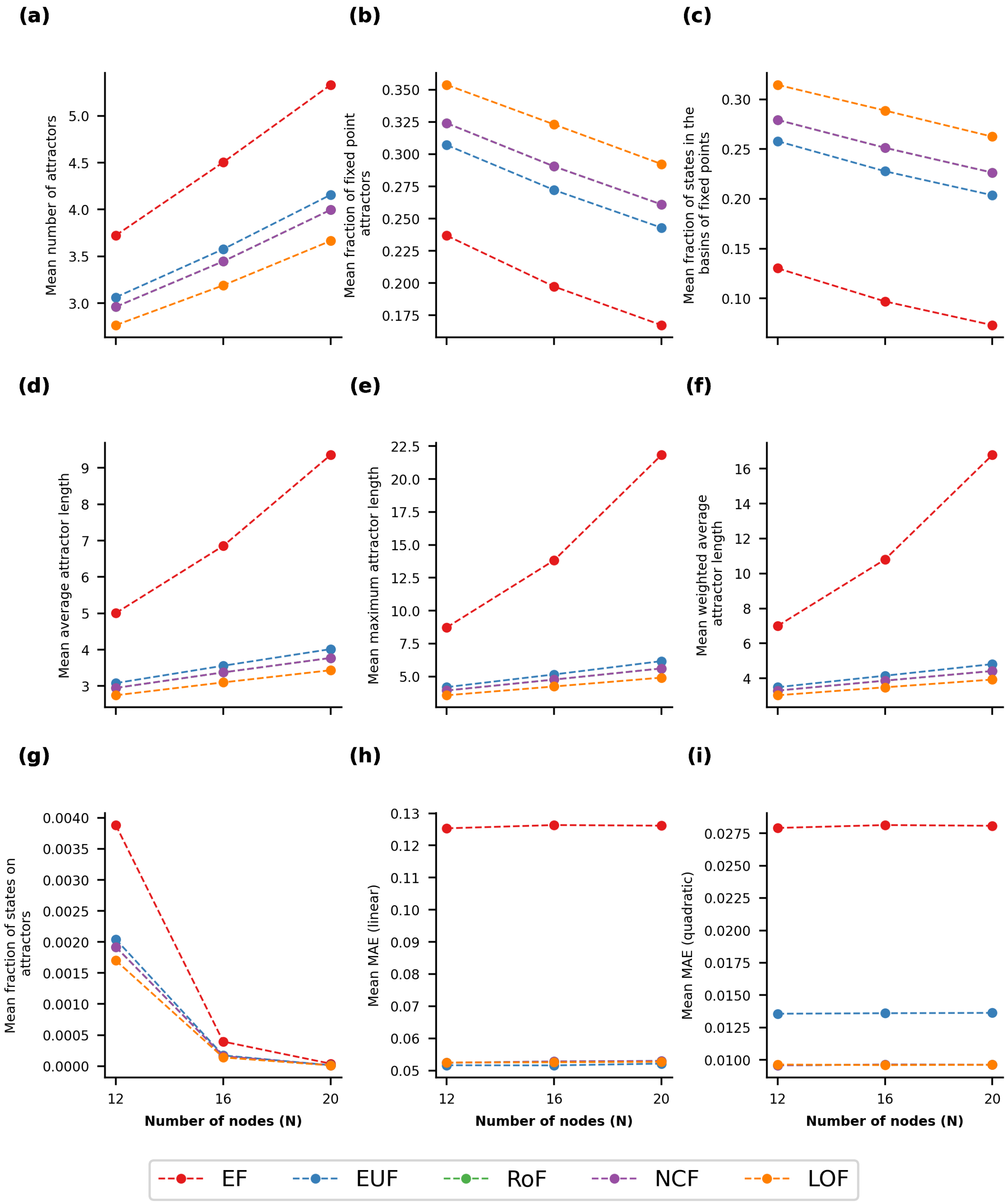


FIG. S65. **Mean of the stability observables for different values of $N$ in the case of the R-P topology with $k_{in} = 3$.** Within each sub-figure, the 5 curves correspond to the 5 different classes of BFs. The data displayed represent the mean value of the corresponding observable at a given value of $N$ for the R-P network topology with $k_{in} = 3$

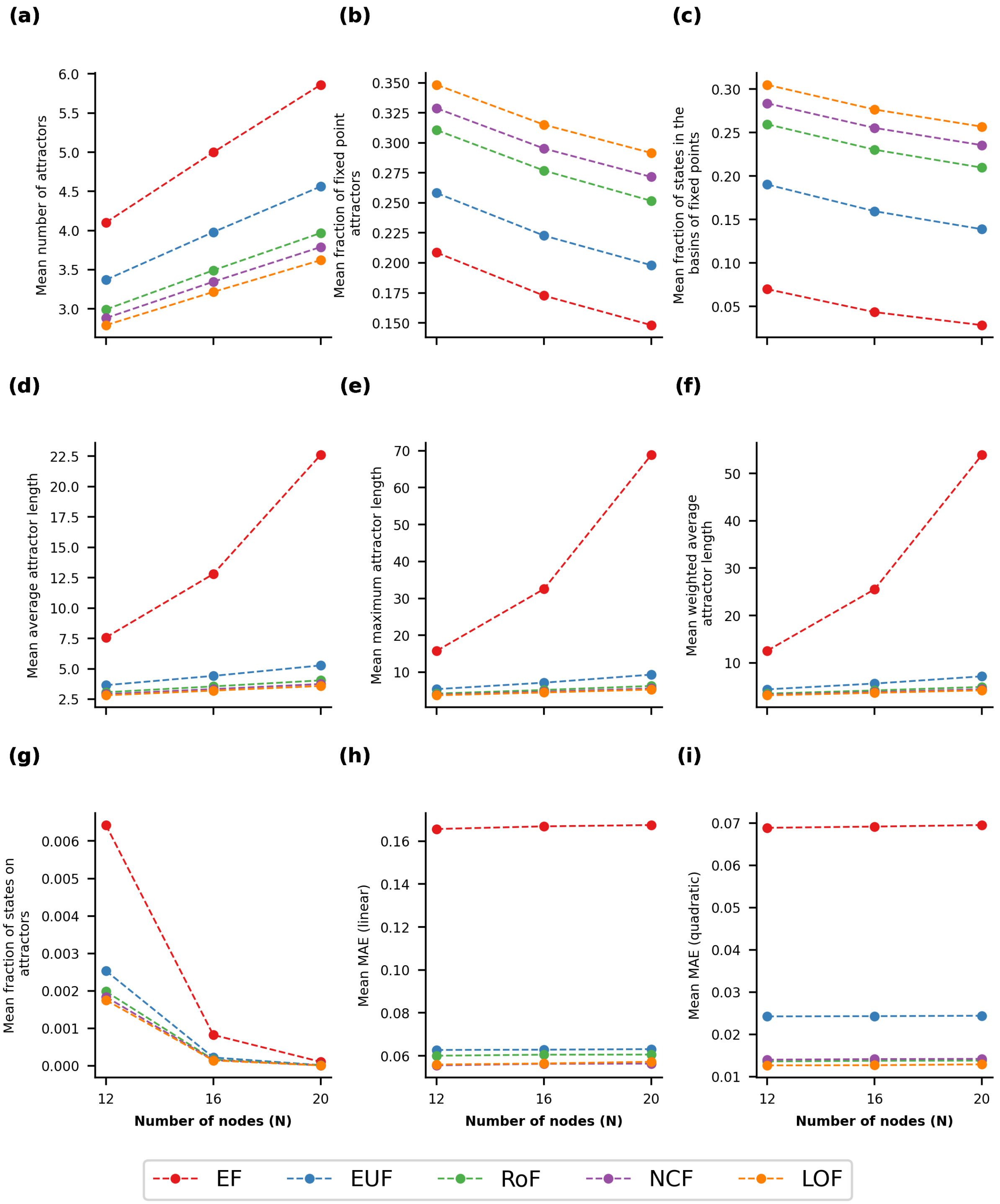


FIG. S66. **Mean of the stability observables for different values of $N$ in the case of the R-P topology with $k_{in} = 4$.** Within each sub-figure, the 5 curves correspond to the 5 different classes of BFs. The data displayed represent the mean value of the corresponding observable at a given value of $N$ for the R-P network topology with $k_{in} = 4$

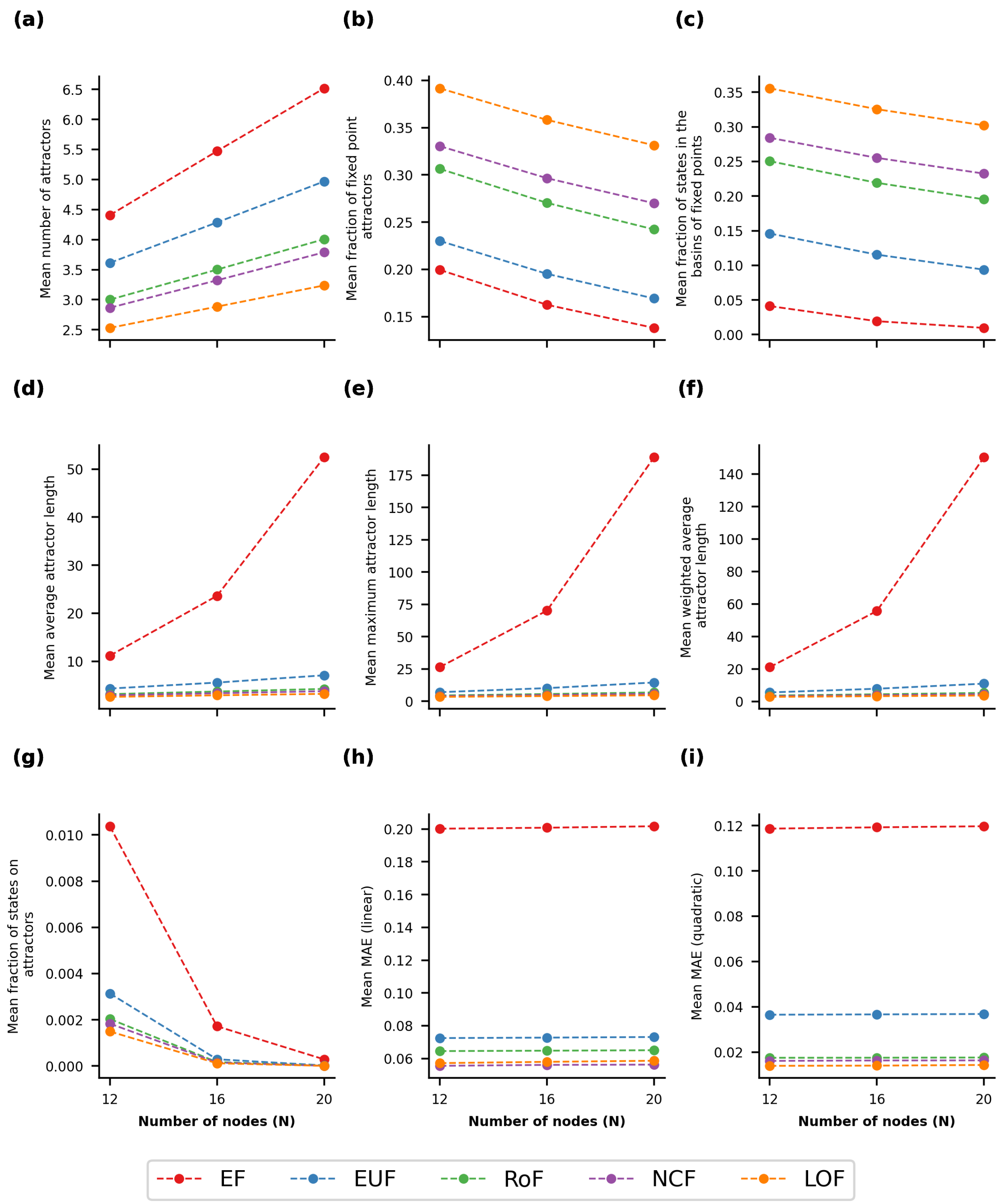


FIG. S67. **Mean of the stability observables for different values of $N$ in the case of the R-P topology with $k_{in} = 5$.** Within each sub-figure, the 5 curves correspond to the 5 different classes of BFs. The data displayed represent the mean value of the corresponding observable at a given value of $N$ for the R-P network topology with $k_{in} = 5$

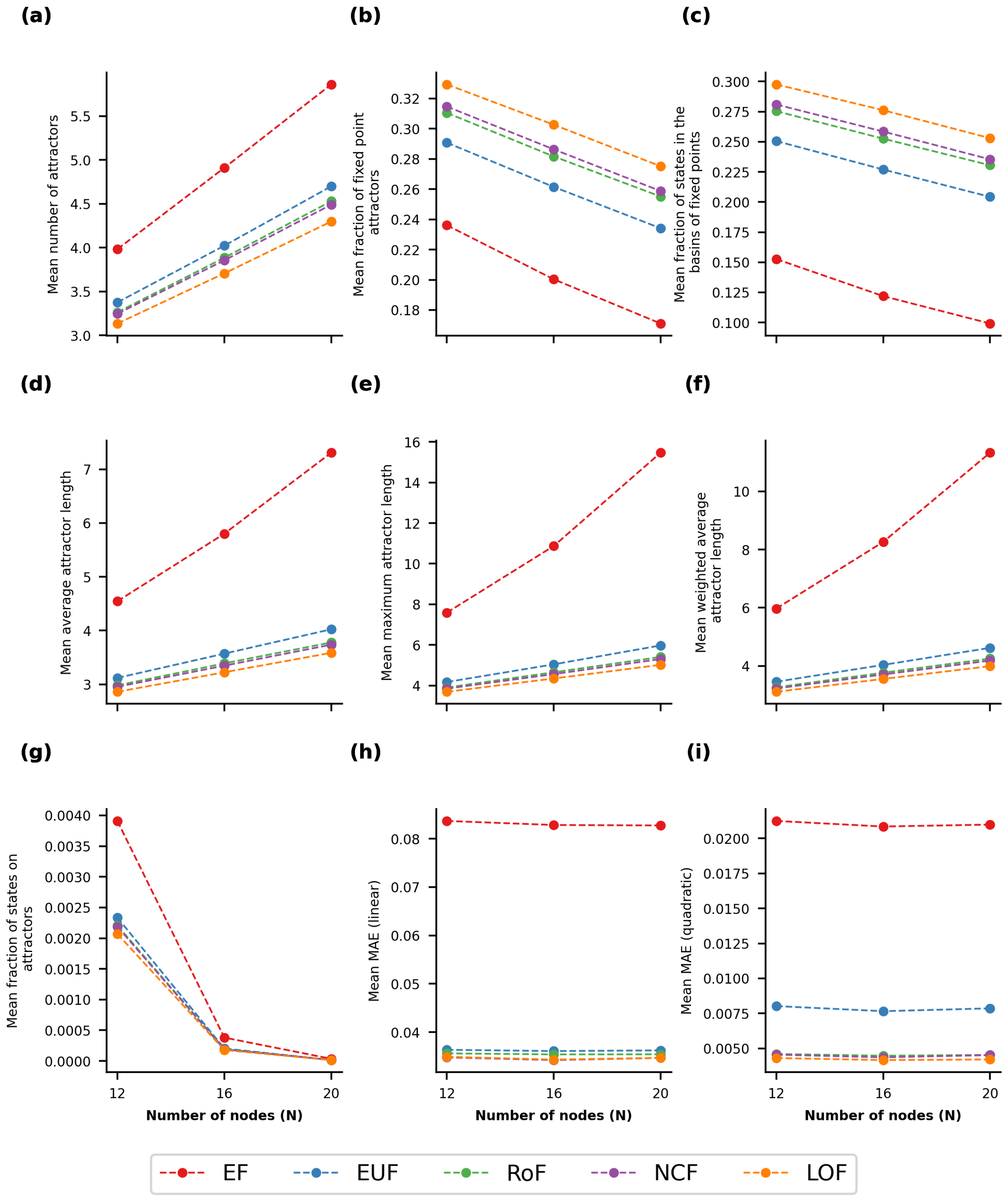


FIG. S68. **Mean of the stability observables for different values of $N$ in the case of the P-P topology with $\lambda = 2$.** Within each sub-figure, the 5 curves correspond to the 5 different classes of BFs. The data displayed represent the mean value of the corresponding observable at a given value of $N$ for the P-P network topology with $\lambda = 2$ (the degrees of each node in the networks are drawn randomly from a Poisson distribution with mean $\lambda = 2$)

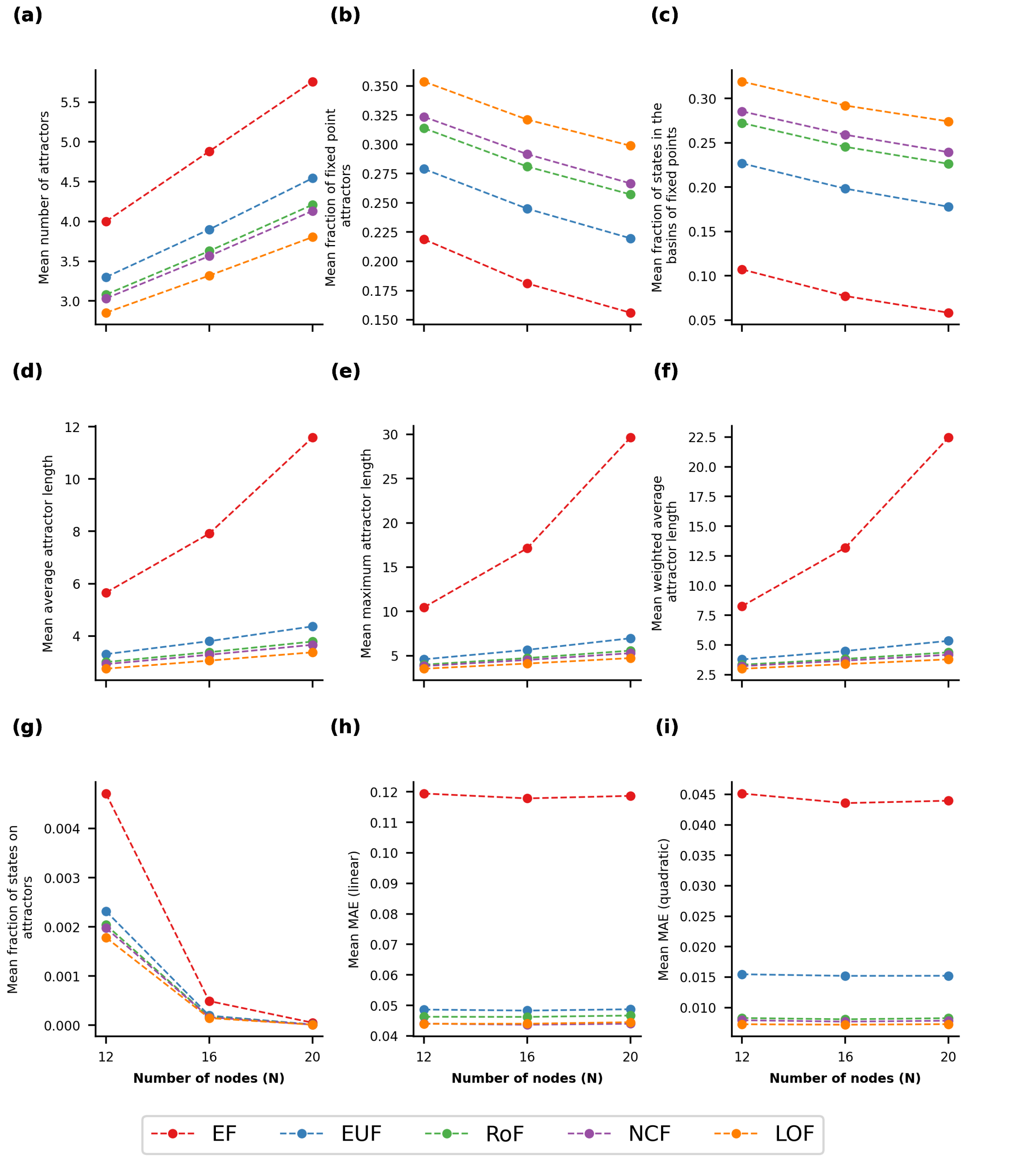


FIG. S69. **Mean of the stability observables for different values of $N$ in the case of the P-P topology with $\lambda = 3$.** Within each sub-figure, the 5 curves correspond to the 5 different classes of BFs. The data displayed represent the mean value of the corresponding observable at a given value of $N$ for the P-P network topology with $\lambda = 3$ (the degrees of each node in the networks are drawn randomly from a Poisson distribution with mean $\lambda = 3$)

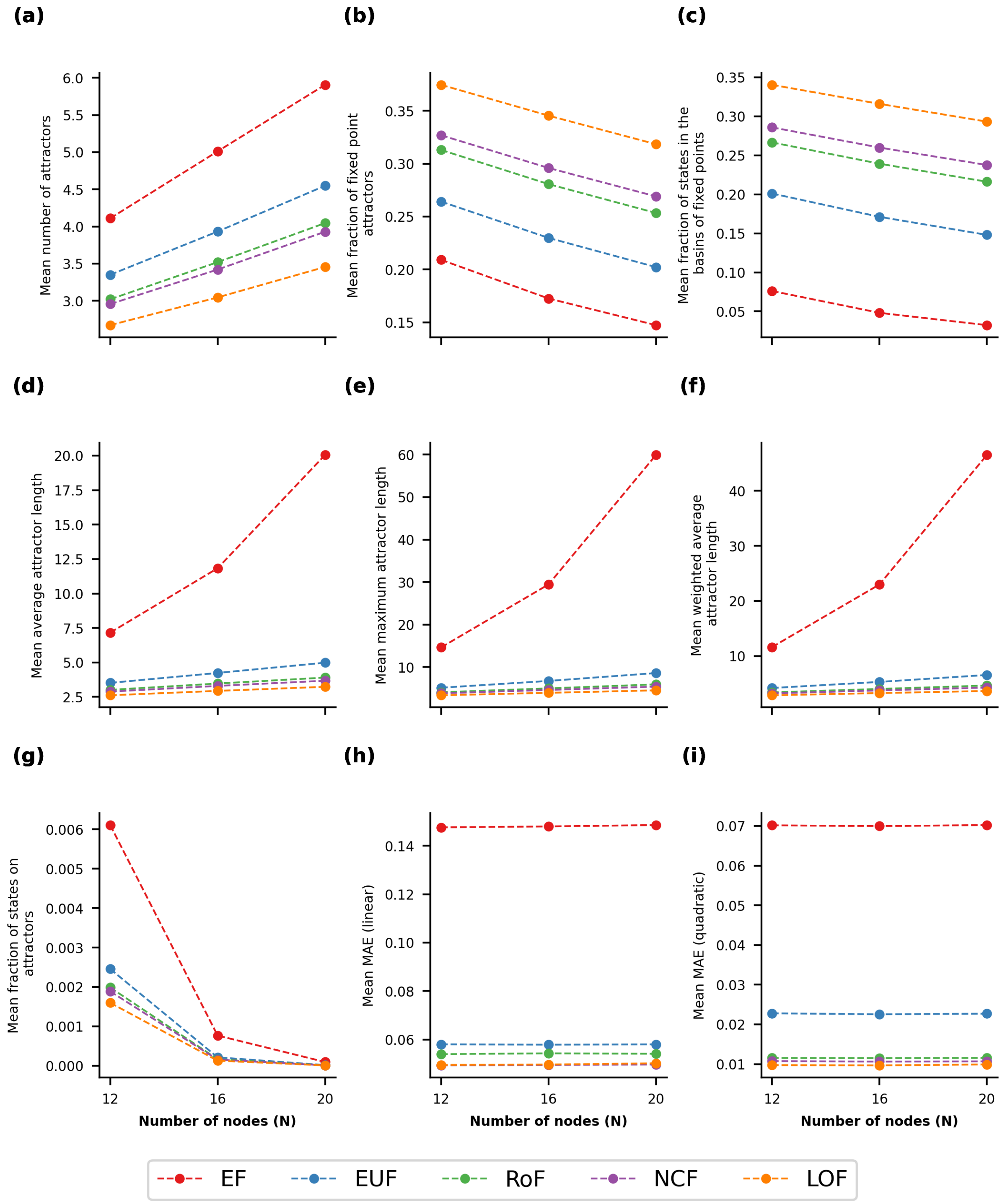


FIG. S70. **Mean of the stability observables for different values of $N$ in the case of the P-P topology with $\lambda = 4$.** Within each sub-figure, the 5 curves correspond to the 5 different classes of BFs. The data displayed represent the mean value of the corresponding observable at a given value of $N$ for the P-P network topology with $\lambda = 4$ (the degrees of each node in the networks are drawn randomly from a Poisson distribution with mean $\lambda = 4$)

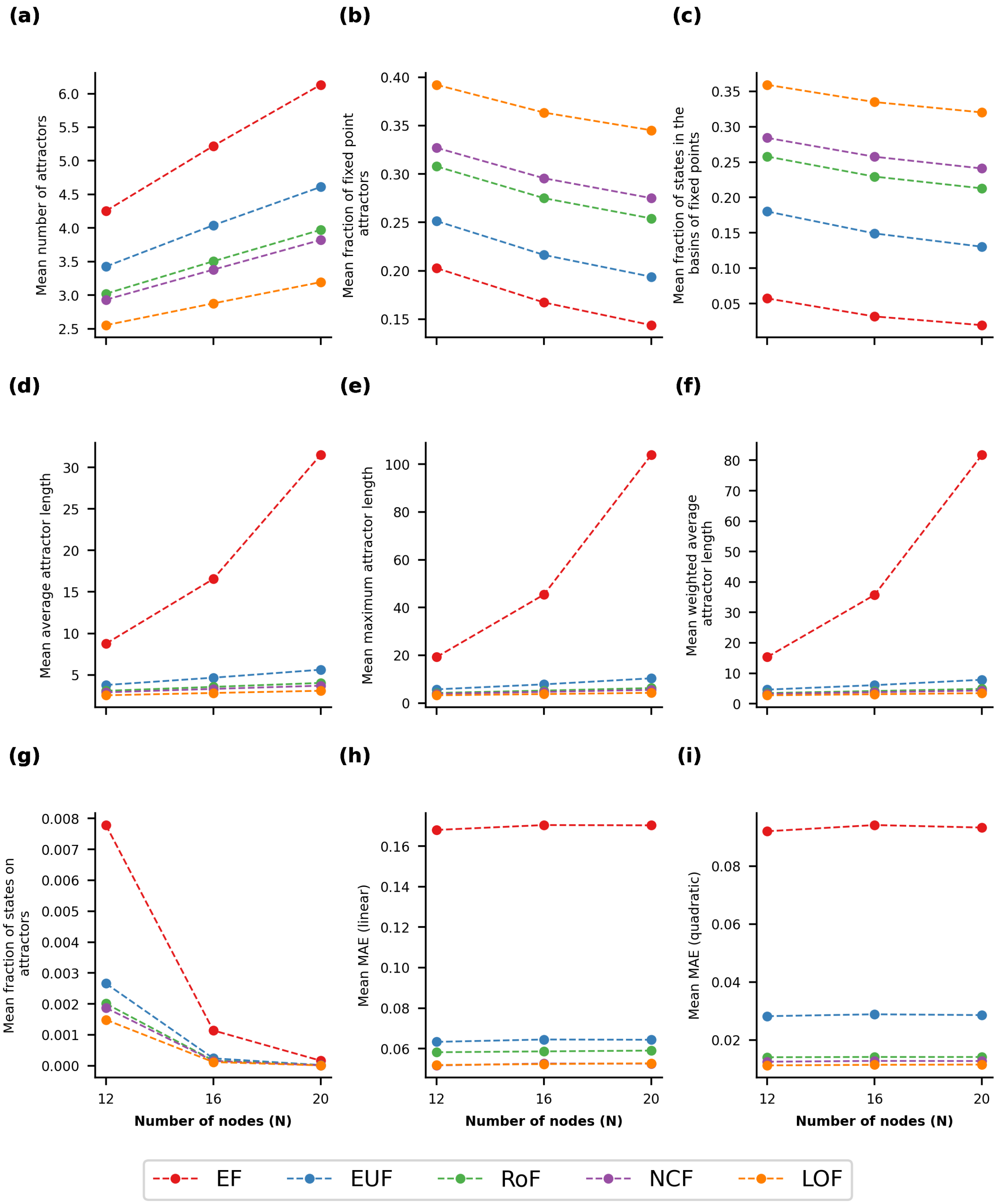


FIG. S71. **Mean of the stability observables for different values of $N$ in the case of the P-P topology with $\lambda = 5$.** Within each sub-figure, the 5 curves correspond to the 5 different classes of BFs. The data displayed represent the mean value of the corresponding observable at a given value of $N$ for the P-P network topology with $\lambda = 5$ (the degrees of each node in the networks are drawn randomly from a Poisson distribution with mean $\lambda = 5$)
